# Conserved strategies underpin the replicated evolution of aridity tolerance in trees

**DOI:** 10.1101/2025.03.06.641812

**Authors:** Ben Halliwell, Chris J. Blackman, Rachael V. Gallagher, Luke A. Yates, Travis G. Britton, Rebecca C. Jones, Dean Nicolle, Gabrielle E. Hartill, Ian J. Wright, Barbara R. Holland, Timothy J. Brodribb

## Abstract

Predicting forest responses to climate change requires a detailed understanding of trait-environment coordination. Adaptation to environment comprises both conserved and labile components of trait variation. However, few studies explore the decomposition of trait-environment relationships in a rigorous phylogenetic framework. Combining a revised phylogeny with trait, climate and soil data to achieve unprecedented species coverage (∼85%), we identified patterns of replicated evolution that allowed the evergreen tree genus, *Eucalyptus*, to rapidly radiate across Australia in response to aridification. Eucalypts from arid regions are short, produce dense wood, and have small, physically robust leaves with high nitrogen content, promoting hydraulic safety and economies in photosynthetic water use. Phylogenetic modelling reveals strong niche conservatism, with adaptation to aridity occurring primarily via clade-level divergences, followed by phylogenetically independent adjustments to local conditions. Ancestral state reconstructions accounting for trends in the paleoclimate record indicate that transitions in aridity tolerance are associated with distinct signals of environmental filtering and directional selection on functional trait variation. However, astonishing repeatability of trait changes in different clades reveals a narrow optimal solution to water availability, opening a path to predict future species distributions from phylogenetically structured trait data and signalling major implications for functional and species diversity under progressive climate change.

## Introduction

Global increases in tree mortality^1,2^ and forest die-back events^3-6^ bring an urgency to understand the process of climatic adaptation in woody plants^7,8^. To predict ecosystem responses to climate change, we must understand which species traits confer resilience to temperature and drought stress, including the decomposition of factors that contribute to this functional trait variation across species and environments^9-11^. This knowledge is critical to effective planning and protection of natural and managed forest systems into the future, both of which are vulnerable to climate induced increases in tree mortality^12,13^.

Comparative analyses emphasise unifying trait spectra that relate plant functional variation to species environmental niches^14-17^. Trait coordination can be understood in terms of economic and physiological trade-offs, such as between tissue lifespan and productivity^17,18^, and the safety and efficiency of water transport^19,20^. Interspecific variation in functional traits therefore reflects the evolutionary outcomes of these trade-offs for fitness across environmental gradients^21,22^. In woody species, adaptation to aridity is thought to involve coordination among trait spectra, leading to whole-plant strategies that integrate evolutionary responses in hydraulic, economic and photosynthetic function^23-26^. However, a major challenge for predicting forest responses to environmental change is that trait-environment relationships vary considerably^27^ due to inherent biological and biogeographic differences between plant clades ^28-31^. One consequence is that broad-scale trait-environment relationships are often poor predictors of local-scale plant responses^9,32^. Phylogenetic models of trait-environment coordination within prominent clades offer a promising avenue for overcoming these challenges of model transferability^33,34^. However, few studies have attempted to decompose trait-trait and trait-environment relationships in such a rigorous phylogenetic framework.

Genus *Eucalyptus* presents a model system to study adaptation to aridity in evergreen trees. Emerging 50-60 Mya in the warm and humid Paleogene^35^, fossil evidence suggests eucalypts became increasingly abundant during the Miocene (23-5 Mya), coincident with the onset of progressive climatic drying and depletion of soil nutrients across the Australian continent^36,37^. This rapid environmental change imposed strong selection and environmental filtering on ancestral eucalypts, leading to multiple independent transitions into arid environments in different lineages (figure 2). These recent, replicated evolutionary events, along with the functional and ecological diversity of eucalypts, present an outstanding opportunity to decompose drivers of trait coordination during adaptation to aridity^38^. Recent drought-induced canopy die-back of eucalypt forest in Australia^39-41^, promoting catastrophic wildfires^42,43^, underscores the urgency of predicting future changes in eucalypt forest to mitigate impacts of climate change on both natural and managed systems.

Drawing on the latest phylogenetic evidence for *Eucalyptus*^44^, we apply multivariate phylogenetic models to assess trait-trait and trait-environment coordination. We leverage a dataset of >22,500 trait observations covering ∼85% of *Eucalyptus* species^45^, by combining AusTraits^46^ data records with targeted sampling of >200 species (table 1). We combine these trait data with environmental data observed at two spatial scales (species mean and site-level) in a single analysis to simultaneously assess 1) the independent effects of climate and soils on trait variation; 2) the decomposition of trait-trait and trait-environment relationships across phylogenetic and non-phylogenetic variance components; 3) whether past changes in trait values represent causes or consequences of transitions to more arid environments.

**Table 1.** Sample size information and summary statistics for each functional trait and environmental niche variable within the dataset used to fit MR-PMM. N_obs_ = total number of observations across species, N_species_ = number of species with data. The mean number of observations per species (and per taxonomic series) for each trait demonstrates good within-species replication across the 767 species included in analyses.

| | $N_{\text{obs}}$ | $N_{\text{species}}$ | proportion<br>of species | mean number of<br>obs per species | mean number of<br>obs per series | min<br>value | max<br>value | units |
| --- | --- | --- | --- | --- | --- | --- | --- | --- |
| leaf area | 8727 | 766 | 0.99 | 11.4 | 72.1 | 155 | 16422 | mm<sup>3</sup> |
| LMA | 6820 | 625 | 0.81 | 10.9 | 60.9 | 42 | 932 | g/m<sup>2</sup> |
| $N_{\text{area}}$ | 2521 | 504 | 0.66 | 5 | 25.2 | 0.17 | 6.19 | g/m<sup>2</sup> |
| $\delta^{13}\text{C}$ | 1860 | 505 | 0.66 | 3.7 | 18.8 | -31.9 | -23.2 | ‰ |
| wood density | 1930 | 385 | 0.50 | 5 | 19.5 | 0.4 | 1.19 | mg/mm<sup>3</sup> |
| max height | 761 | 761 | 0.99 | - | - | 1 | 99.6 | m |
| temperature | 767 | 767 | 1 | - | - | 6.3 | 27.8 | °C |
| moisture | 767 | 767 | 1 | - | - | 0.06 | 2.25 | unitless |
| soil P | 767 | 767 | 1 | - | - | 0.007 | 0.033 | ‰ |
| soil bulk density | 767 | 767 | 1 | - | - | 1.08 | 1.7 | g/cm<sup>3</sup> |

## Results

### Adaptation to aridity represents a whole-plant strategy

We focus on 6 traits with well-established links to hydraulic, economic and photosynthetic function: leaf area, leaf mass per area (LMA), leaf N per area (N_area_), discrimination between ^13^C and ^12^C carbon isotopes in leaf tissue (δ^13^C), wood density, and species maximum potential height (max height); and 4 key dimensions of species environmental niche: mean annual temperature (temperature), mean aridity index (moisture), mean soil phosphorus concentration (soil P) and mean soil bulk density (bulk density).

Two principal components explained 66.7% of the phenotypic variation across the 6 traits and 4 environmental niche variables (figure 1; table S1; S2). PC1 (49.3%) captured variation in moisture, soil P and a suite of functional traits, whereas PC2 (17.4%) primarily captured variation in temperature, soil bulk density and leaf N_area_. In contrast to results from global analyses reporting independent axes of variation in size and economic traits^14,15^, our targeted analysis of *Eucalyptus* reveals extensive coordination between these traits, consistent with variation along a whole-plant fast-slow economic spectrum (figure 1). Eucalypts occupying hot dry environments are characterised by short stature, dense wood, small leaves, as well as high LMA, N_area_ and δ^13^C (figure 1). The phylogenetic distribution of functional trait variation indicates that divergence along this trait spectrum has occurred repeatedly during adaptation to aridity gradients within different eucalypt clades (figure 2).

**Figure 1.**
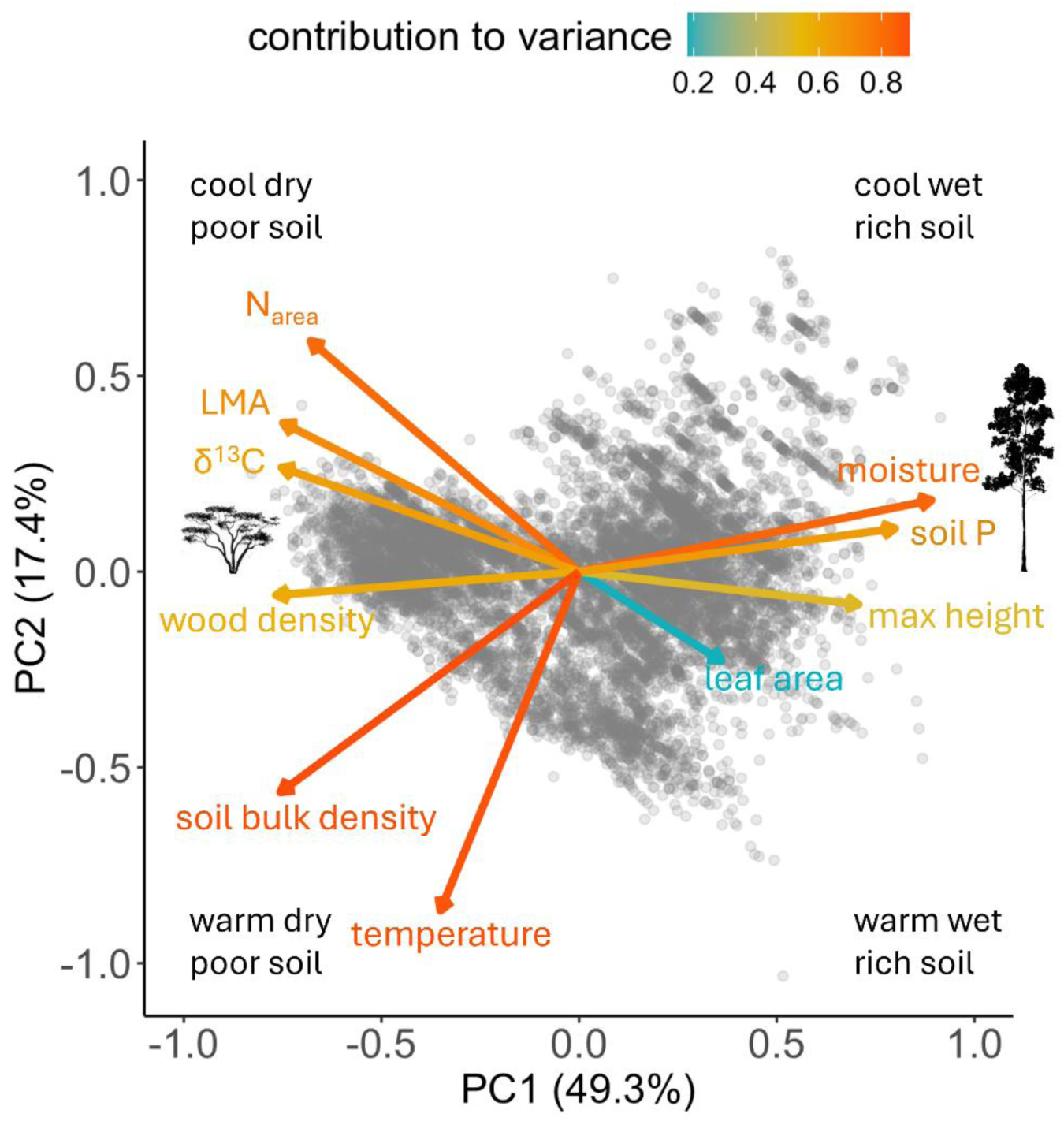
Adaptation to aridity in *Eucalyptus* represents a whole-plant strategy. Principal components analysis of functional traits and environmental factors using data gap-filled with predictions from the fitted MR-PMM. PC1 (49.3% of variance) primarily captured trade-offs in functional traits along an axis of variation in moisture index and soil variables. PC2 (17.4% of variance) primarily captured variation in temperature, soil bulk density and Narea. PC loadings and cumulative variance are shown in tables S1; S2.

**Figure 2.**
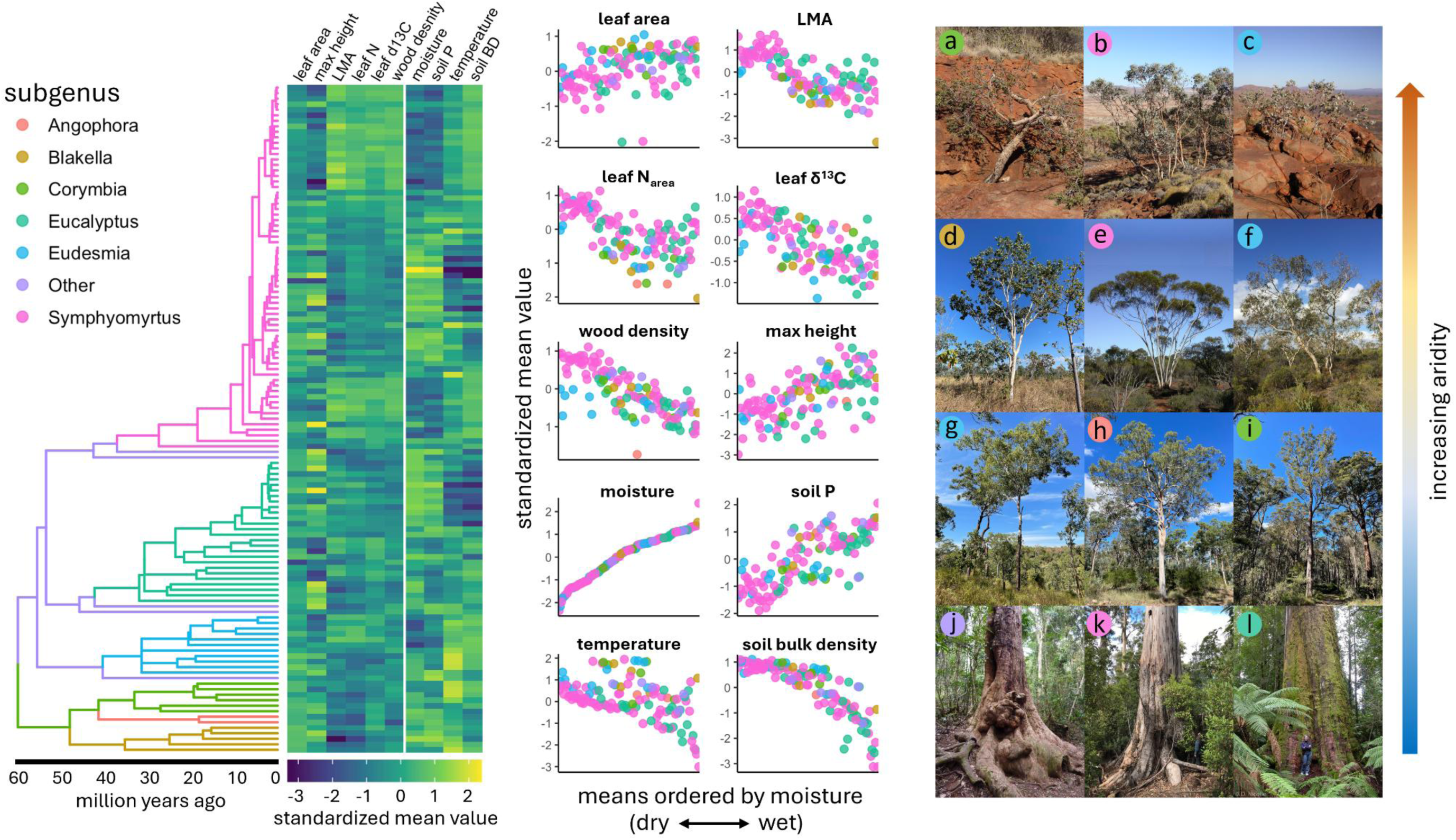
Conserved strategies underpin the replicated evolution of aridity tolerance in different *Eucalyptus* clades. Phylogenetic distribution of functional trait values and environmental niche variables for *Eucalyptus* (N_species_=767) using gap-filled data imputed by MR-PMM. Taxonomic series-level topology (left) coloured by subgenera is plotted against a matrix of series-level means (centre) for each trait and environmental niche variable. Scatterplots (centre) show series-level means for each trait and niche variable ordered by moisture. The phylogenetic distribution of trait values reveals that multiple clades have evolved the same set of trait values to cope with dry climates, evident as blocks of similar trait values among clades characterised by low moisture. Remarkable convergence in whole-plant phenotype to moisture availability is observed across multiple independent origins of aridity tolerance in different clades (right): a) *E. deserticola*, b) *E. repullulans*, c) *E. gamophylla*, d) *E. grandifolia,* e) *E. pileata,* f) *E. erythrocorys,* g) *E. miniata,* h) *E. leiocarpa,* i) *E. gummifera,* j) *E. microcorys,* k) *E. globulus,* l) *E. regnans* (photos by Dean Nicolle).

### Decomposing trait-environment relationships

Understanding the process of climatic adaptation at different biological scales requires that we separate trait variance due to local environmental conditions from that due to conserved trait-environment relationships^47^. The multi-response phylogenetic mixed model (MR-PMM)^48^ presented here leverages an unprecedented dataset for *Eucalyptus*, both in terms of phylogenetic coverage and within-species replication of functional traits (table 1), to jointly estimate and partition these effects (figure 3). These analyses allow us to decompose correlations between species traits and environmental niche variables into phylogenetic and non-phylogenetic components. Furthermore, for trait observations matched with environmental data from the site individuals were sampled, we calculated environmental anomalies (standardized differences between species mean environmental niche values and site-level environmental conditions) for each species-site combination. By fitting species-specific environmental anomalies as fixed effects in the MR-PMM, we can separate the influence of site-level environmental conditions on trait variation *within-species*, when estimating correlations between traits and niche variables *between-species*. We first consider the effect of environmental anomalies, then explore the decomposition of trait-trait and trait-environment correlations within components of variance that remain unexplained by these site-level effects.

**Figure 3.**
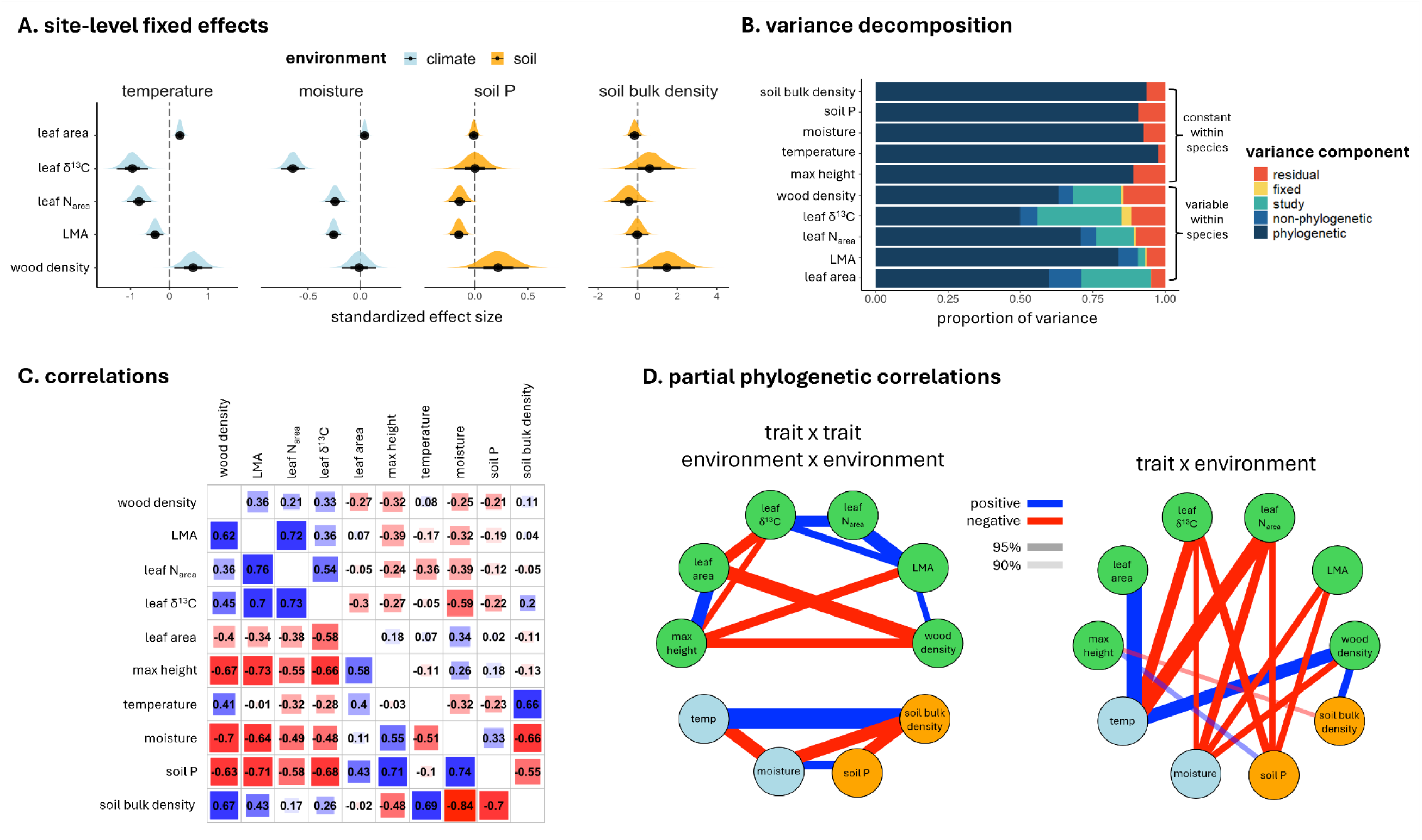
Within-species, traits vary with site-level environmental conditions, but effects are relatively weak. Decomposition of the remaining trait (co)variance across species reveals strong phylogenetic signal underlying a conserved network of trait-trait and trait-environment relationships. Results from MR-PMM fit to 100 candidate series-level topologies (N_species_ = 767): **A** Posterior fixed effect estimates of site-level environmental anomalies for each functional trait associated with site-level data. **B** Variance decomposition for each functional trait and environmental niche variable. Traits that are constant for a species are partitioned into phylogenetic and residual components, while traits varying within-species are partitioned into phylogenetic, non-phylogenetic, study, fixed, and residual components. **C** Phylogenetic (lower triangular) and non-phylogenetic (upper triangular) between-species correlations between response variables from MR-PMM. Values represent the posterior mean estimate for each correlation coefficient. **D** Network graphs showing significant (90% and 95% CI) partial phylogenetic correlations (positive = blue edges, negative = red edges) between functional traits (green), climate (light blue) and soil (orange) niche variables. Edge widths are proportional to partial correlation coefficients. All correlation estimates derive from the same MR-PMM model fit but are grouped and plotted in separate networks for illustrative purposes. Note that partial phylogenetic correlations between max height and environmental niche variables were markedly stronger when phylogenetic random effects were modelled using the species-level topology (figure S2D).

### Responses to local environment

Within-species variation in functional traits was significantly related to site-level environmental anomalies (figure 3A), indicating adjustments in trait values to deviations from species mean environmental conditions. On average, individuals from warmer sites have higher density wood, larger leaves, lower N_area_ and more negative δ^13^C, implying less efficient water use during photosynthesis. However, elevated temperature had similar effects to elevated moisture on trait values. For example, eucalypts respond to both hotter and wetter sites by decreasing N_area_ (figure 3A). This suggests a partly compensatory mix of trait responses to hotter and drier (i.e., more arid) sites, which predominate given strong negative covariance between temperature and moisture across the bioclimatic niche space of *Eucalyptus* (figure 2, figure 3C). Climate effects were particularly pronounced for leaf δ^13^C, highlighting adjustments in photosynthetic properties as a key response to site-level temperature and water availability. Soil effects were detected for LMA, N_area_ and wood density, indicating a shift to more conservative growth strategies in challenging soil conditions. Specifically, eucalypts respond to low relative soil P by increasing LMA and leaf N_area_, and to high relative soil bulk density by increasing wood density (figure 3A).

### Drivers of trait variation

Fixed effects of environmental anomalies explained considerably more variance in wood density and leaf δ^13^C than any other functional traits (figure 3B). However, these within-species, site-level effects explained only a tiny fraction (1-3%) of total trait variance (figure 3B; table S3). Study effects consistently explained a greater proportion of trait variance than fixed effects, likely due to characteristics of study sites not captured by our fixed effects (e.g., micro-climate, topography, biotic factors), as well as differences in methodology or measurement technique between studies. Non-phylogenetic between-species effects (i.e., effects of species identity after accounting for phylogenetic relationships) were small (5-11%) and generally attributed less variance than either study effects or residual errors, indicating that divergences between closely related species contribute relatively little to observed trait variation. In contrast, for every trait except leaf δ^13^C, the phylogenetic component of variance was greater than all other variance components combined (figure 3B). Indeed, phylogenetic signal (0 < λ < 1) for functional traits ranged from 0.48 (leaf δ^13^C) to 0.87 (max height), and was extremely high for species mean environmental variables (λ > 0.85), reflecting strong niche conservatism.

### Phylogenetic versus non-phylogenetic correlations

Decomposing between-species trait correlations into phylogenetic and non-phylogenetic components confirms that similar relationships operate on both levels (figure 3C), broadly reflecting patterns observed in PCA of phenotypic data (figure 1). This indicates a relatively scale-free evolutionary response to environmental gradients, with similar patterns of trait coordination in the climatic divergence of closely related species as in the climatic divergence of major clades over deeper time (figure 2, figure S1). However, phylogenetic correlations were considerably stronger (figure 3C, lower triangular), emphasising the ecological significance of clade-level divergences (figure 2).

### Identifying a core network of integrated trait responses

To identify candidate causal relationships between response variables within the major phylogenetic component of trait variance, we computed partial correlations from the fitted phylogenetic covariance matrix. These correlations revealed a highly integrated network of trait-trait and trait-environment relationships, consistent with a conserved, whole-plant response to environmental gradients (figure 3D). Trait coordination can be described as a trade-off between two modules of positively covarying traits: a ‘fast’ acquisitive trait module comprising size-related traits (leaf size and max height), and a ‘slow’ conservative trait module comprising traits with hydraulic, economic and photosynthetic functions (wood density, LMA, N_area_, δ^13^C, figure 3D). This trade-off defines a major axis of ecological strategy in *Eucalyptus*, separating species across a continuum between productivity and persistence in a whole-plant ‘fast-slow’ economic spectrum.

Moisture, temperature, and soil P each displayed (non-zero) partial correlations with a range of traits, indicating that adaptation to these environmental factors involves direct responses in hydraulic, economic and photosynthetic function (figure 3D). Moisture was strongly negatively associated to traits within the ‘slow’ trait module (wood density, LMA, N_area_, and δ^13^C), where-as evidence for positive associations with the ‘fast’ trait module (max height, leaf size) varied depending on the taxonomic scale at which phylogenetic effects were modelled (species- vs series-level topology, figure S2D). Pairwise partial correlations generally supported a direct functional link between moisture and max height (table S4, figure S2D), but suggest a relative decoupling between moisture and leaf size. We find particularly strong partial correlations between temperature and each of wood density, leaf area, N_area_, and δ^13^C, suggesting tight coordination between these traits and the temperature niche of eucalypt lineages. Partial correlations between soil bulk density and both max height and wood density highlight adjustments in hydraulic properties as a primary mechanism of adaptation to reduced soil water holding capacity (figure 3D; figure S2D). Taken together, these results emphasise a coordinated strategy favouring hydraulic safety, conservative growth economics, and high photosynthetic water-use-efficiency (WUE) as a consistent feature of adaptation to hot, dry, and poor soil environments in *Eucalyptus*.

### Ancestral state reconstructions

Coordination between plant traits and environment can arise by filtering on standing variation in trait values, i.e., during colonisation or rapid environmental change (environmental filtering)^49^, or by directional selection on trait values, i.e., adaptation *in situ* (local adaptation)^50^. To distinguish the relative importance of these processes, we performed ancestral state reconstructions that account for historical shifts in the Australian paleoclimate to evaluate whether changes in trait values are likely to represent a cause (environmental filtering) or consequence (local adaptation) of transitions to more arid environments (see methods). To investigate whether similar processes have acted on functional trait variation at different points along the aridity continuum, we compared the inferred sequence of trait changes during transitions between mesic (M), semi-arid (S) and arid (A) environments. Transitions from M (moisture index > 0.5) to S (moisture index < 0.5), and from S to A (moisture index < 0.2), environments have occurred many times independently across different clades within our broad phylogenetic sample of *Eucalyptus* (figure 4A; table S5). This analysis of replicated evolution reveals that selection has repeatedly produced the same solution during transitions to more arid environments. Node categories representing transition types between different aridity states were strongly associated with variation in reconstructed trait values (figure S3), reflecting a progressive shift toward slower leaf and wood economic strategies, and higher photosynthetic WUE, with increasing aridity (figure 4B).

**Figure 4.**
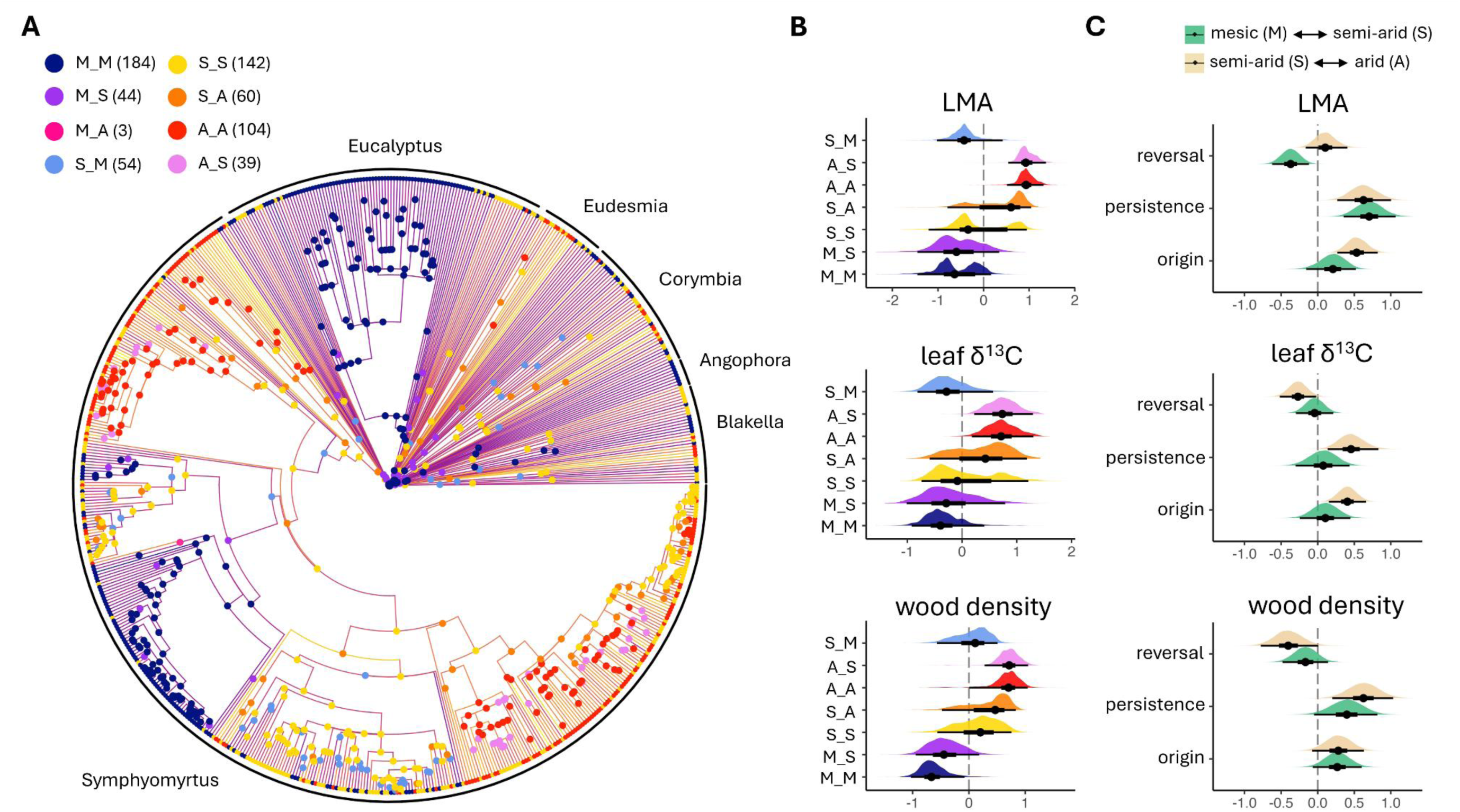
Ancestral state reconstructions suggest that adaptation to moisture gradients in *Eucalyptus* has involved both environmental filtering and directional selection on functional trait values, with more stringent filtering during the colonisation of arid environments. **A** Species level phylogeny (N_species_ = 631) showing major Eucalypt clades. Branch lengths have been transformed^51^ by δ = 10 for plotting. Branch colour maps an ancestral state reconstruction of species mean moisture index (MI) according to a shift model that accounts for strong directional change toward drier climates in Australia since the Miocene (23 Mya). Internal nodes are classified according to their own reconstructed values of MI (mesic = MI > 0.5, semi-arid = 0.2 < MI < 0.5, arid = MI < 0.2) and those of their descendants (see methods). Posterior mean counts for each node category are indicated in parentheses within the legend. Tip labels indicate whether the corresponding species MI value classifies it as mesic (purple), semi-arid (yellow) or arid (red). **B** Posterior distributions of trait values at internal nodes for each transition category from ancestral state reconstruction. **C** Contrasts between node transition categories representing different stages in the evolution of aridity tolerance (see methods). Contrasts are computed and reported for transitions between mesic and semi-arid environments (green), as well as between semi-arid and arid environments (cream). The mean (point), 50% (heavy whisker) and 95% (light whisker) CI of each posterior distribution are shown in black.

To test for differences in reconstructed trait values between node categories while accounting for phylogenetic relationships, we computed contrasts from posterior samples of an MR-PMM specifying node category as a fixed effect when estimating ancestral states. Figure 4C presents results for wood density, LMA, and δ^13^C, traits that each show partial correlations with moisture (figure 3D), and relate broadly to hydraulic, economic and photosynthetic functions (results for other traits shown in figure S4). Evidence for environmental filtering during transitions from M to S environments (M_S versus M_M) was limited to a subset of traits; M ancestors giving rise to S descendants (M_S) had lower max height, as well as marginally higher wood density and LMA, than M ancestors giving rise only to M descendants (M_M) (figure 4C; figure S4). For most traits, evidence was stronger for directional changes during persistence in S environments (S_S versus M_M), consistent with local adaptation to water limitation (figure 4C; figure S4). However, results differed when considering transitions from S to A environments, representing tolerance of more extreme water limitation. We find stronger evidence for environmental filtering, particularly on LMA, leaf N_area_ and δ^13^C, during transitions from S to A environments (S_A versus S_S). We also find strong evidence for directional changes in trait values during persistence in A environments (A_A versus S_S). Finally, we investigated whether reversals to less arid environments were associated with opposing changes in the same network of traits. Consistent with this, we observed coordinated reversion of trait values toward a faster, riskier, and more acquisitive strategy during reversals from A to S, and from S to M, environments (figure 4C, figure S4).

## Discussion

Drawing on data for 85% of *Eucalyptus* species, we find strong and consistent functional trait responses during adaptation to environmental gradients. Species respond to site-level deviations from mean environmental niche values with finely tuned adjustments in a range of functional traits (Figure 3A). However, these effects are generally small and insufficient to explain broad trait-environment relationships across species (Figure 3B). Instead, strong phylogenetic signal reveals that these relationships are driven primarily by conserved components of trait covariance, reflecting a replicated pattern of trait changes during the climatic divergence of eucalypt clades. Traits associated with hydraulic, economic and photosynthetic functions showed strong relationships with species moisture niche, consistent with an integrated whole-plant response to aridity gradients (figure 1; figure 3C).

Pairwise trait relationships were broadly consistent with predictions from leaf economic theory^17^, least-cost theory^52,53^, pace of life syndromes^26^ and the allometry of hydraulic traits^54,55^ (figure 3A). These include positive correlations between LMA and wood density (linking leaf and wood economic spectra), N_area_ and δ^13^C (linking carboxylation capacity and stomatal limitation on photosynthesis), and leaf size and max height (linking variation in the size of plant organs), as well as negative correlations consistent with trait trade-offs, including between wood density and max height (stress-tolerance versus growth rate). A strong positive relationship between LMA and N_area_ emphasises coordination between leaf economics and the manner in which nitrogen and water use in photosynthesis is optimised in relation to water limitation^53,56^, but also contributions of nitrogenous compounds other than RUBISCO, especially cell walls, which also increase LMA^57,58^.

More broadly, the functional trait network we identify indicates coordination within and between plant tissues; opposing modules of traits (figure 3D) define a spectrum of whole-plant strategy from productivity to persistence that integrates tissue economics, hydraulic vulnerability, and photosynthetic WUE^16,26^. These results are consistent with detailed comparative ecophysiology studies of *Eucalyptus*^59-62^, which report extensive coordination between economic, hydraulic and photosynthetic functions. Indeed, the conserved trait network we identify may have been key to the ecological success of *Eucalyptus*, by aligning major axes of functional variation^63^ with the harsh aridity gradients that have come to characterise the Australian continent.

The trait-environment relationships we find for *Eucalyptus* accord well with those reported for other woody taxa, indicating species from arid environments tend to be short^55^, produce dense wood^16^, and have small, robust leaves with more active stomatal regulation^23^. Of the traits examined here, wood density and max height have been linked most directly to the capacity of trees to tolerate acute water-stress^64-68^. Previous studies considering a smaller number of *Eucalyptus* species have linked wood density to moisture availability^69^ (N_species_ = 28), as well as directly to measures of hydraulic vulnerability^70^ (N_species_ = 20). We confirm a strong association between wood density and aridity tolerance across a much larger sample of *Eucalyptus* in this study (N_species_ = 767), including partial correlations with both the temperature and moisture axes of species’ environmental niches. Our analyses also support strong associations between species max height and environmental niche variables (figure 1; figure 3C), consistent with previous work on *Eucalyptus*^62,71^. Evidence for partial correlations between max height and climate varied depending on the scale at which phylogenetic effects were modelled, with stronger evidence for these effects when using the species-level phylogeny (figure 3D; figure S2D). This may suggest that conserved relationships between max height and climate can to some extent be explained by covariance with other functional traits and environmental factors. However, as max height is defined at the species level, a likely explanation for this model dependence is that lower order divergences in the phylogeny (i.e., species-level relationships) provide the necessary resolution to detect conserved relationships between max height and environmental factors. Indeed, when using the species-level phylogeny, max height and wood density showed equally strong partial phylogenetic correlations with moisture (figure S2D).

Across all analyses, LMA showed a strong relationship with the moisture niche of *Eucalyptus* species. This contrasts with global analyses, which indicate that LMA is only weakly associated with climatic variation at both the species and community trait level^14,72^ (though see^9^). Our analyses show that, within phylogenetically constrained species groups, LMA may be tightly coupled with climatic variation. Indeed, ancestral state reconstructions identified both environmental filtering and directional selection on LMA as features of transitions into arid environments (Figure 4C). This suggests that, along with traits that confer resilience to acute drought stress, successful evolutionary strategies involve optimising leaf economics toward slow, conservative growth^26^. Extreme phylogenetic signal in LMA (λ = 0.85) further indicates an evolutionary rigidity in leaf economic strategies, with individual clades occupying narrow regions of the economic spectrum. LMA was also strongly negatively related to soil P content (figure 3D), agreeing with global analyses that link leaf economics to soil fertility, in particular the availability and cost of acquiring essential soil nutrients^14^.

Both N_area_ and δ^13^C displayed partial correlations with moisture and temperature niche, indicating fundamental roles of WUE and stomatal regulation in the capacity of eucalypt species to persist in arid environments. Higher N_area_ implies greater investment in the carboxylating enzyme RUBISCO, meaning that, at a given level of stomatal conductance, leaf-internal CO_2_ during photosynthesis is lower – leading to more efficient water use (higher WUE) for a given photosynthetic rate per unit leaf area (A_area_) and site VPD^53^. Lower ci:ca is reflected in leaf δ^13^C^73,74^, which may explain the exceptionally strong positive correlations between N_area_ and δ^13^C observed in our analyses (figure 3C, figure 3D). While WUE is clearly crucial for plant performance in hot dry conditions, the negative partial correlation between N_area_ and temperature can be explained by reduced investments in RUBISCO with increasing efficiency of carboxylation at higher temperatures^52,56,75^.

Of all functional traits, leaf area showed the weakest relationships with both species mean environmental niche variables and site-level environmental conditions (figure 3A). Despite this, leaf area was strongly associated with max height, wood density and δ^13^C, consistent with a role of leaf size in plant allometric relationships affecting hydraulic demand and water-use strategies^54^.

Ancestral state reconstructions indicate that coordinated trait changes represent both a cause and a consequence of the colonisation of water-limiting environments in *Eucalyptus* (Figure 4B, C). Evidence for environmental filtering on trait variation was generally stronger during transitions from semi-arid to arid environments (S_A) compared with transitions from mesic to semi-arid environments (M_S), particularly for leaf traits (figure 4C, figure S4). This supports the notion that extreme environments impose greater constraints on viable trait combinations, leading to more stringent trait filtering^76^. A notable exception was max height, which showed stronger effects during transitions from mesic to semi-arid environments (figure S4), consistent with the drastic reduction in tree height observed between wet- and dry-sclerophyll eucalypt forest (figure 2). Our analyses further suggest that, following transitions to more water-limiting environments, ancestral eucalypts persisting in those environments underwent strong directional evolutionary change, becoming markedly shorter, with higher wood density, LMA, N_area_, and δ^13^C (figure 4; S2). Finally, during reversals to less water-limiting environment (S_M and A_S), we observed an opposing pattern of trait changes consistent with environmental filtering on the same coordinated suite of traits. The astonishing repeatability of trait changes across multiple independent transitions in aridity tolerance suggests a narrow optimal solution to water availability in *Eucalyptus^38^*. Indeed, similar trait spectra characterise drought tolerant species across many clades and biogeographic regions, suggesting convergence on a fundamental archetype of water stress tolerance in trees^9,77^.

Most variation in functional traits and species environmental niche can be attributed to phylogenetic effects (figure 3D), reflecting major ecological divergences between eucalypt clades (figure 2). This emphasizes phylogenetic niche conservatism^47,78^ as a major feature of climatic adaptation in *Eucalyptus*, mirroring findings for woody species at broader phylogenetic scales^79,80^. Indeed, the progressive emergence of aridity gradients in Australia since the Miocene (23 Mya)^36,37^ is consistent with a biogeographic history of species filtering processes followed by rapid adaptation of aridity tolerant lineages, invoking coordinated changes in traits that jointly maximised fitness in these novel environments; a scenario well supported by our ancestral state reconstructions (figure 4C). Niche conservatism demonstrates that organisms tend to establish and persist in environments to which they are already well adapted^81^. Examples of arid niche conservatism in *Eucalyptus* include the speciose sections of subgenus *Symphyomyrtus*; *Bisectae*, *Dumaria* and *Glandulosae* (figure S1). These clades radiated very recently (5-10 Mya), together accounting for the majority (78%) of arid distributed *Eucalyptus* species and are each characterised by the same suite of traits conferring slow growth and stress tolerance (figure S1). Such clades are likely to expand in distribution as climate change progresses, with profound implications for the functional and phylogenetic diversity of Australian forests^82^, including fire regimes^83^ and ecosystem services^84^. Specifically, our results suggest that the initial impact of increases in semi-arid habitat with climate change^85^ is likely to be strong species filtering based on tree height, wood density and LMA, promoting a rapid transition from tall, closed-canopy forest to sparse, open woodland, with consequences for primary productivity, carbon sequestration, and the frequency and intensity of wildfires.

The remarkable strength of phylogenetic signal (figure 3D) in functional traits and environmental niche variables we report for *Eucalyptus* confirms that phylogenetic information is severely underutilised in global predictive vegetation models^86-88^. Indeed, species hydraulic, economic, photosynthetic, and phenological traits are strongly conserved at multiple phylogenetic scales^89-91^, presenting a tremendous opportunity to improve prediction of future species distributions and abundances^87^, as well as trait-related ecosystem services such as primary productivity and carbon sequestration^92^. For *Eucalyptus*, a paleoclimatic trend of increasing aridity induced the adaptive radiation and competitive dominance of the drought tolerant *Symphyomyrtus* clade (figure 2), while reducing many humid adapted Gondwanan lineages to relictual species^93^ (figure 2j). As climate change progresses, further declines in the viability of ‘fast’, hydraulically risky, economic strategies across multiple biomes puts entire clades at risk of extirpation and extinction^94^, with major consequences for the functional and phylogenetic diversity of ecosystems^95,96^. Similar patterns of conserved hydraulic strategy within prominent woody clades such as *Quercus*^97^, *Juniper*^98^, *Magnolia*^99^ and Fabaceae^100^, as well as more broadly across both angiosperms and gymnosperms^90,101^, indicate a concentration of drought tolerance within a limited subset of current tree species diversity, raising the alarm for catastrophic, phylogenetically-biased biodiversity loss as we pass critical thresholds of global aridification^102^.

Encouragingly, strong phylogenetic correlations between functional traits and environmental variables also highlight opportunities for imputation of scarce physiological data^103,104^, vastly expanding our capacity to assess species hydraulic risk^89,105^. Using MR-PMM on a global plant dataset, Sanchez-Martinez et al.^103^ demonstrate improved prediction of leaf hydraulic traits by leveraging phylogenetic correlations with leaf economic traits. As trait-trait and trait-environment relationships are often tighter within phylogenetically constrained species groups^19,27,60^, such correlations may be particularly valuable for predicting hydraulic traits within speciose, phylogenetically structured clades, such as *Eucalyptus*. This approach has enormous potential for parameterizing species-specific hydraulic models of drought-induced tree mortality^106^, aiding the assessment of species vulnerability and ecosystem resilience under climate change scenarios. A clear direction for future work is therefore to clarify the strength of relationships between hard (physiological) and soft (morpho-anatomical) traits both within and between prominent woody clades, so that phylogenetic information can be leveraged most effectively at a global scale. Discovery and refinement of soft traits that correspond strongly to physiological thresholds of water stress tolerance (i.e., cavitation resistance and hydraulic safety margins), while being cost effective to observe and scalable across diverse taxa (e.g., xylem area per unit leaf area (XLA)^70^), is a particularly valuable avenue for future research.

Adaptation to aridity in *Eucalyptus* has proceeded via multivariate shifts along a whole-plant ‘fast-slow’ economic spectrum, prioritizing hydraulic safety and photosynthetic WUE, over resource acquisition and rate of return on tissue investment. These findings support a general trade-off between productivity and persistence in plant strategy^26^, and link variation along this trait spectrum to multiple independent transitions in aridity tolerance in a major tree genus. Strong conservatism in functional traits and environmental niche variables of woody lineages worldwide suggests that species responses to climate change will be phylogenetically biased, putting entire clades at risk of rapid decline and threatening to disrupt the structure and function of ecosystems. Out of this adversity comes a valuable opportunity to leverage the phylogenetic revolution to better predict species responses to environmental change.

## Methods

### Trait Data

We compiled all available observations of leaf mass per area (LMA), leaf area, leaf nitrogen per area (N_area_), leaf δ^13^C, maximum potential height (max height), and wood density for *Eucalyptus* species in the AusTraits^46^ database (figure S5). We augmented this dataset by 1) adding records from observations made by authors in the field (N = 1321) and at Currency Creek Arboretum, South Australia (N = 2107) between 2022-2024; 2) adding records from recent publications^60,70,107^ yet to be incorporated into AusTraits; and 3) deriving trait values when missing observations could be calculated from available trait data, e.g., calculating N_area_ from N_mass_ and LMA, or leaf area from leaf length and width measurements (see below). The combined dataset contained 22,650 observations on our 6 focal traits across 767 eucalypt species. Total sample sizes for each trait, and the proportion of eucalypt species for which data were available for each trait are shown in table 1.

#### Data cleaning and taxonomic assignment

Details of data cleaning and exclusion are summarised in figure S5. We excluded records derived from experimental studies and those reporting observations on juvenile or immature specimens. We included records reporting raw trait values (i.e., observations on an individual specimen) as well as species mean trait values estimated from multiple observations within- or between-sites. We restricted data on height to records of maximum species height, taking the maximum value if multiple values were reported for a given species.

In AusTraits, leaf dimension measurements (length and width) were available for many more *Eucalyptus* species than leaf area measurements (96% vs 68% of species). To leverage these data in our study, we used leaf length and width measurements for each species to calculate leaf area as,

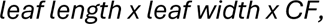

where CF = 0.68 is the correction factor (CF) applicable to *Myrtaceae* based on Shrader et al. (2021). To confirm the accuracy of this method, we regressed observed leaf area values against those calculated from length and width measurements (where both were available) and confirmed a tight linear relationship (Pearsons’s correlation coefficient = 0.97).

### Climate and Soil Data

To separate the effects of local environmental conditions, phylogenetic history, and study effects on functional trait variation, we integrated climate (mean annual temperature, mean moisture index) and soil (phosphorus content, bulk density) data (table 1) observed at two different spatial scales (species mean values and site-level values) into a single model.

For species mean values, climate and soil characteristics of each species range were extracted by intersecting point locations associated with digitised herbarium collections (latitude and longitude) with long-term gridded datasets of climate and modelled soil variables and extracting values. Climate data was sourced from the CHELSA dataset^109^ where current climate is represented by average conditions between 1979-2013, and soil data from the National Soil and Landscape Grid of Australia (https://esoil.io/TERNLandscapes/Public/Pages/SLGA/). This provided a distribution of values across raster cells for each environmental variable for each species, from which we calculated species mean values, representing the average climatic and soil conditions experienced by each species across their geographic range. These species mean environmental variables (values constant within-species) were then fit alongside functional trait variables (values variable within-species) in a multi-response (MR) phylogenetic mixed model (PMM), allowing us to decompose trait-environment correlations across different levels in the model hierarchy.

For site-level environmental data, we used the procedure described above to extract point-location climate and soil values for each trait observation reporting latitude and longitude of the collection site (N = 4120). For each observation, we then subtracted the relevant species mean environmental niche values from the site-level environment values, yielding a set of variables expressing the differences between site-level environmental conditions and species mean environmental niche values, i.e., environmental anomalies. Records for which site-level data were unavailable (N = 6872) were given a value of 0 for these anomaly variables and were thus assumed to have been collected at sites representing species mean environmental conditions. By fitting species-specific environmental anomalies as fixed effects in the MR-PMM, this approach allowed us to separate and test the influence of site-level environmental conditions on functional trait variation within-species, when estimating correlations between niche characteristics and functional traits between-species.

### Phylogeny

Hybridisation and introgression among closely related species complicate the phylogenetic inference of *Eucalyptus*^110,111^, leading to difficulties in confidently assigning sister species relationships. To account for the potential impacts of this phylogenetic uncertainty, we integrated the latest phylogenetic evidence^44^ and taxonomic revisions^45,112^ to augment the largest time-calibrated phylogeny currently available for *Eucalyptus*^35^. Then, we used this augmented tree to model phylogenetic structure at two different taxonomic scales (species-level and series-level). This approach allowed us to maximise the number of species and trait observations included in analyses, while explicitly incorporating phylogenetic uncertainty into parameter estimates. For all analyses, we follow the recent taxonomic revision proposed by Nicolle et al. (2024a; 2024b), in which all previously recognised eucalypt genera (i.e., *Eucalyptus*, *Corymbia*, *Angophora* and *Blakella*) are treated as subgenera of a unified genus *Eucalyptus*.

First, we used the recent phylogeny of Crisp et al. (2024) to infer a strict consensus tree of *Eucalyptus* at the taxonomic section level. We began by generating 1000 candidate section-level topologies from each of the two maximum likelihood partition trees presented in Crisp et al. (2024) by randomly sampling one species from each taxonomic section and pruning the respective tree to those taxa. We combined these 2000 candidate section-level topologies into a single set to infer a strict consensus tree. We then augmented the “ML1” phylogeny presented in Thornhill et al. (2019) to match this backbone topology by 1) removing taxa that were discordant with this consensus tree, 2) re-locating subgenus *Eudesmia* to be sister to subgenera *Eucalyptus* and *Symphyomyrtus*, in line with Crisp et al. (2024); and 3) excluding taxa identified as inter-sectional or inter-series hybrids according to Nicolle et al. (2024).

Ancestral state reconstructions require species-level topology to leverage information on trait differences between closely related species that have nonetheless diverged in their climatic preferences. However, for inferring and decomposing correlations between traits, it may be preferable to collapse the phylogeny to a higher taxonomic rank to include as much data as possible in analyses (see below). We therefore used our augmented species-level tree to further generate candidate series-level topologies, by sampling one species from each taxonomic series and pruning the tree to those taxa. The resulting trees represent hypotheses of phylogenetic relationships between *Eucalyptus* taxonomic series, rather than species, providing several advantages for downstream analyses. In particular, by treating species as replicates of a taxonomic series we: 1) avoid strict assumptions about sister-species relationships that may be complicated by hybridisation and introgression; 2) include data for all species with resolved series-level taxonomy, including those that do not feature in the species-level phylogeny; 3) are able to directly propagate phylogenetic uncertainty into our analyses by fitting models over a sample of candidate topologies and combining posterior estimates for inference.

### Statistical Analyses

#### Multi-Response Phylogenetic Mixed Models (MR-PMM)

We used Bayesian MR-PMM to decompose correlations between species functional traits and environmental niche variables into phylogenetic and non-phylogenetic components^48^. Specifically, we fit all functional traits and environmental niche characteristics jointly as response variables, removing the global intercept to estimate separate intercepts for each response variable, and using correlated random effects to specify phylogenetic and non-phylogenetic between-species covariance matrices. We modelled multi-level structure in the data by fitting random intercepts across species (accounting for within-species replication) and studies (accounting for study effects). Retaining observation-level data allowed us to estimate within-species variances for each functional trait, allowing us to more accurately estimate phylogenetic and non-phylogenetic trait (co)variances^113^. All MR-PMM were fit using the MCMCglmm package^114^ in R.

#### Phylogenetic imputation

Traits varied in the extent of missing data, ranging from 50% missingness at the species level for wood density to 1% missingness for leaf area and max height (table 1). Missing response values are permitted in MCMCglmm, with missing values imputed conditional on the full phylogenetic covariance structure of the model. In our case, this means that both phylogenetic and non-phylogenetic between-species trait correlations inform the imputation of missing values. One benefit of this approach over multiple imputation procedures is that imputation uncertainty is naturally propagated through to the posterior distribution of parameter estimates. We used the gap-filled dataset predicted from the MR-PMM fit to conduct PCA (figure 1) and visualise the phylogenetic distribution of functional trait values (figure 2).

#### Model Settings

For all analyses based on series-level phylogeny (see above), we incorporated phylogenetic uncertainty by fitting models over a sample of 100 candidate topologies, and pooling MCMC samples across fits prior to inference. We assessed convergence by 1) visually inspecting traces of the MCMC posterior estimates; and 2) confirming potential scale reduction factors (*r̂*), a convergence diagnostic test that compares within- and between-chain variance^115^, were ≤1.01 for all parameter estimates. Estimates for all variance components converged successfully using parameter expanded priors with (variance) V = I_m_ (an identity matrix of dimension equal to the number of response traits, *k*), and (degree of belief) ν = 1*k*+1 for random effects^116^, nitts = 11000, burn-in = 1000, and thin = 10. We used the default independent normal priors with (mean) = 0 and V = 10^10^ for fixed effects. We performed model validation via predictive assessment using approximate leave-one-out (LOO) cross-validation (CV) procedures^117,118^ (figure S8) and posterior predictive checks (figure S9).

Parameter estimates from models are reported as posterior means and 95% credible intervals (table S4). For hypothesis tests of the significance of effects estimated by MR-PMM (see below), we checked whether the 95% CI of the posterior distribution of estimates crossed zero.

#### Phylogenetic signal

We calculated posterior samples of the phylogenetic signal (λ) in each trait and environmental factor from the fitted MR-PMM as

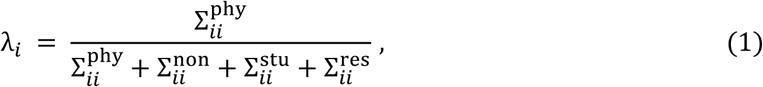

where 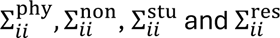 are the estimated phylogenetic (between-species), non-phylogenetic (between-species), study and residual variances for trait *i*, respectively.

#### Correlation estimates

Based on the MR-PMM decomposition, we estimated correlations between traits, and between traits and environmental factors on both the phylogenetic and non-phylogenetic level. We calculated the phylogenetic correlation between two response variables as

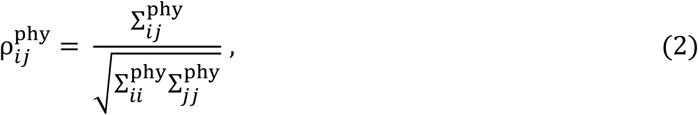

Where 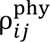 is the phylogenetic correlation between traits *i* and *j*, 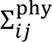 is the phylogenetic covariance between traits *i* and *j*, and 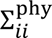 is the phylogenetic variance in trait *i*. Non-phylogenetic between-species correlations were calculated similarly by substituting elements of **Σ**^ind^ into eq. 2.

As for interpretation, phylogenetic correlations represent correlations between traits that are conserved over evolutionary time, i.e., the tendency for traits to change together (and be conserved) during the divergence of phylogenetic clades. Non-phylogenetic correlations represent correlations between traits that are independent of phylogenetic structure, i.e., the tendency for traits to covary after accounting for conserved relationships. Thus, by comparing phylogenetic and residual correlations, we can evaluate the extent to which trait (co)variation can be explained by conserved differences between clades versus differences between individual species that are independent of phylogenetic affiliation.

#### Partial Correlation Estimates

To identify candidate causal relationships between response variables, we calculated partial correlations and associated credible intervals for all pairs of response variables from the precision of the phylogenetic trait covariance matrix^48,119^. The precision matrix (inverse of the covariance matrix) estimates the undirected conditional dependency among a set of variables, from which we can derive partial correlations; pairwise relationships between variables after accounting for covariances with all other variables^120,121^. For example, if we obtain data on three covarying species traits x, y and z, the conditional dependency between x and y is the component of covariance between these traits that cannot be explained by z. If we assume no important omitted variables, significant partial correlations are consistent with direct relationships between traits, as opposed to relationships manifest from cross-correlation with other variables. We used pcor.shrink() from the corpcor^122^ R package to compute partial correlations using the default regularisation settings. By conditioning on all other response variables, (non-zero) partial correlations between species traits provide evidence for direct functional or mechanistic associations, rather than correlations arising from trait responses to the environment^48^. Similarly, partial correlations between traits and environmental factors provide evidence for direct associations between trait values and the ability to persist and compete across environmental gradients, rather than correlations due to relationships with other traits.

#### Ancestral State Reconstruction

Given strong phylogenetic signal in all traits and environmental factors (figure 3B), we used ancestral state reconstruction to evaluate whether changes in trait values represent causes or consequences of transitions to arid environments, i.e., to assess hypotheses relating to the sequence of evolutionary events. We did this using a two-step process following the approach of Cornwallis et al. (2017).

For step one, we used an MR-PMM fit to the species-level phylogeny to estimate ancestral states for moisture index (MI) at all internal nodes. These estimates represent the expected MI of ancestral species, after accounting for model multilevel structure, according to a Brownian motion model of evolution. We then adjusted these Brownian ancestral state estimates according to a shift model, which assumes that all ancestral *Eucalyptus* lineages occupied humid (MI > 0.65) niches prior to the Miocene boundary (23 Mya), before experiencing a directional trend toward more arid niches into the present day (figure S6). Specifically, we adjusted ancestral states of internal nodes prior to the Miocene boundary (node depth > 23 Mya) by an amount equal to the difference between the root state estimated by Brownian motion (MI = 0.46) and the MI defining humid environments (MI = 0.65). We adjusted ancestral states of internal nodes following the Miocene boundary (node depth < 23 Mya) by an amount equal to (23 – D)*R, where D is the node depth and R is the rate of directional change implied by the supplied root state. This procedure yielded a set of ancestral state estimates for MI that represent Brownian evolution from a humid root state (MI = 0.65) up until 23 Mya, followed by a directional trend toward lower MI values between 23 Mya-present (figure S6). This scenario more accurately reflects the paleoclimatic history of Australia, which is characterised by progressive aridification since the Miocene, due to northward drift of the Australian continent and establishment of the Antarctic Circumpolar Current^36,37^. Nonetheless, sensitivity analyses confirmed that results from contrasts (see below) were similar whether we applied the shift model or simple Brownian motion (figure S10).

For step two, we used these reconstructed MI values to compare reconstructed functional trait values between parent nodes that did and did not give rise to more arid descendants (figure S7). Specifically, for each posterior sample of ancestral MI estimates, we first classified each internal node as mesic (M) if the estimated MI > 0.5, semi-arid (S) if the estimated MI was 0.5 > X > 0.2, or arid (A) if the estimated MI < 0.2. We then re-classified all internal nodes by adding information about the state of their descendant nodes. For example, a parent node was classified as M_M if it was M and gave rise only to M descendants (child nodes); M_S if it was M and gave rise to at least one S descendant (representing an origin, or transition, from mesic to semi-arid environments); S_S if it was S and gave rise only to S descendants; and S_M if it was S and gave rise to at least one M descendant (representing a reversal, or transition, to mesic environments). We used this procedure to classify all internal nodes in the phylogeny. We then fit an MR-PMM taking only functional traits as response variables, specifying a trait-level fixed effect of node classification (possible levels: M_M, M_S, M_A, S_S, S_A, S_M, A_A, A_S, A_M), and a phylogenetic random effect linked to ancestral nodes, to estimate the influence of node class on reconstructed trait values. Importantly, the model was fit only to trait data at the tips, which treats tips as replicate observations (descendants) of parental nodes when estimating the influence of node classification on reconstructed trait values. This approach means that uncertainty about reconstructed trait values at more ancient evolutionary divergences is naturally propagated through to the fixed effect estimates of node classification.

To test whether reconstructed trait values differed for nodes representing **origins** of transitions to more arid environments (i.e., nodes directly preceding a transition to a more arid environment, i.e., M_S and S_A) consistent with environmental filtering, we constructed contrasts from the posterior samples of the fitted model by subtracting the relevant posterior distribution of effect sizes for each transition type. Specifically, by subtracting fixed effect estimates associated with no change in aridity state (e.g., M_M) from those associated with transition to a more arid state among descendants (e.g., M_S), and checking whether the 95% CI of the resulting distribution of contrast values crossed zero. These contrasts allowed us to test for differences in reconstructed trait values associated with the initial transition to more arid environments. We then used this technique to create additional contrasts that tested for changes in trait values associated with **persistence** in more arid environments (S_S – M_M and A_A – S_S) consistent with adaptation *in situ*, and filtering during **reversals** to less arid environments (S_M – S_S, and A_S – A_A). Parameter uncertainty from the MR-PMM fit was propagated into ancestral state reconstructions by classifying nodes and calculating contrasts for each posterior sample.

## Acknowledgments

We thank Brad M. Potts for helpful discussions regarding eucalypt phylogenetics. We thank Vinod Jacob and Emma E. Sumner for assistance with field data collection. This work was funded by The Australian Research Council Centre of Excellence for Plant Success in Nature and Agriculture (CE200100015).

## Data Accessibility

Data and R code sufficient to replicate all analyses are available at https://github.com/Benjamin-Halliwell/Eucalyptus_Replicated_Evolution

## Supplementary figures

**Figure S1.**
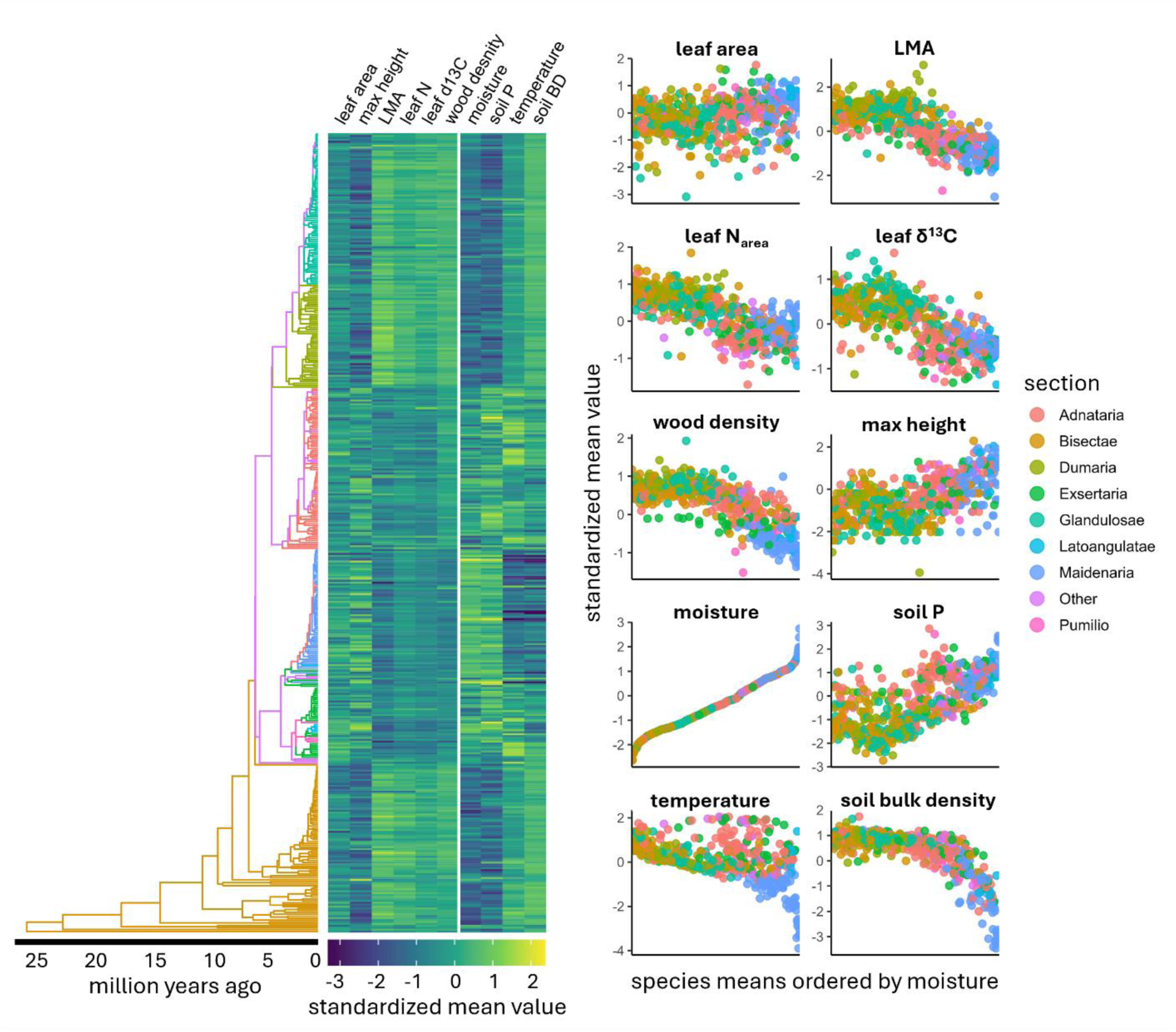
Phylogenetic distribution of functional trait values and environmental niche variables in *Eucalyptus* subgenus *Symphyomyrtus* using gap-filled data imputed by MR-PMM. Species-level topology (left) coloured by section is plotted against a matrix of species-level means (centre) for each trait and niche variable. Scatterplots (right) show species-level means for each trait and niche variable ordered by moisture niche

**Figure S2.**
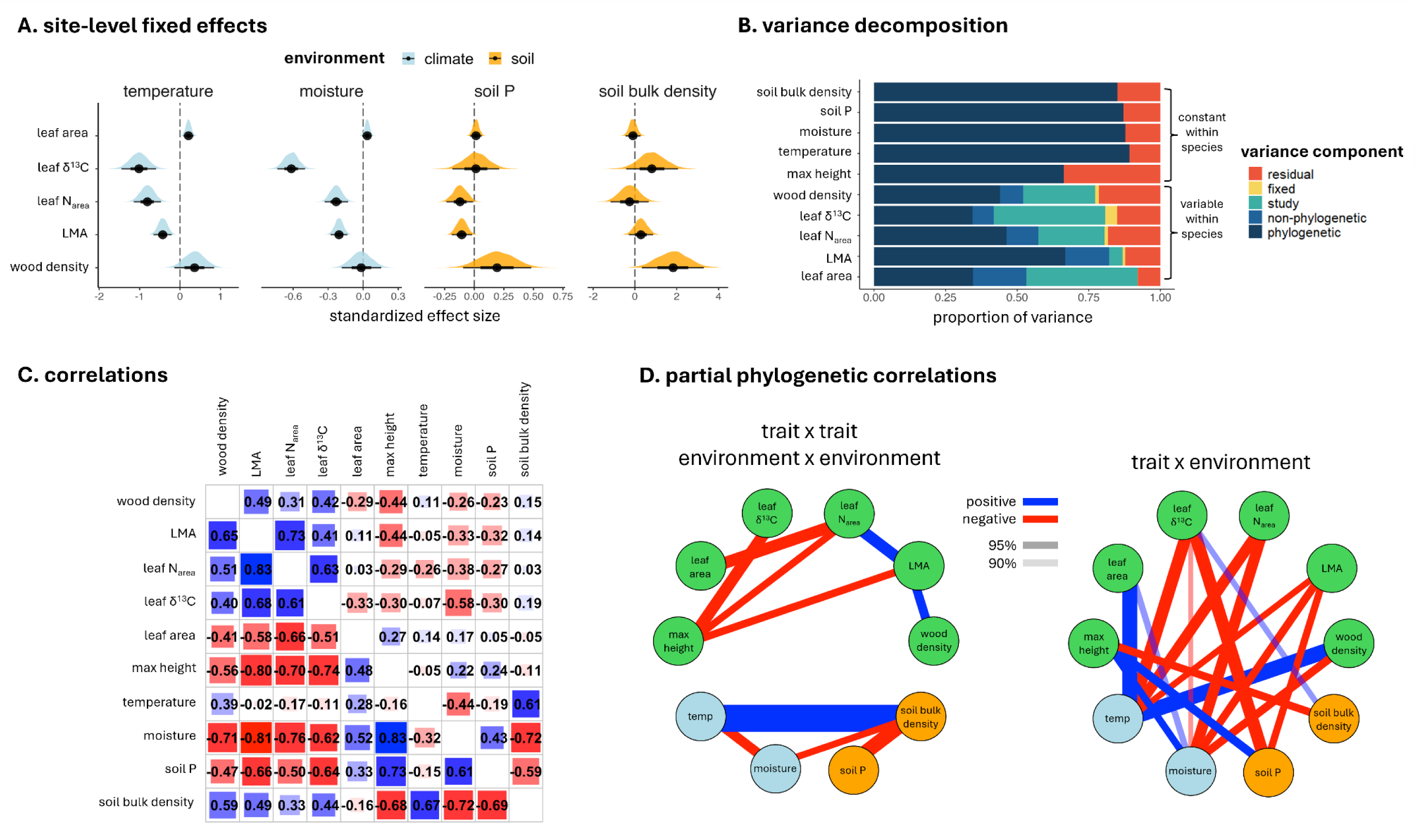
Results from MR-PMM fit to species-level phylogeny (N_species_ = 631): **A** Posterior fixed effect estimates of site-level environmental anomalies for each functional trait associated with site-level data. **B** Variance decomposition for each functional trait and environmental niche variable. Traits that are constant for a species are partitioned into phylogenetic and residual components, while traits varying within-species are partitioned into phylogenetic, non-phylogenetic, study, fixed, and residual components. **C** Phylogenetic (lower triangular) and non-phylogenetic (upper triangular) between-species correlations between response variables from MR-PMM. Values represent the posterior mean estimate for each correlation coefficient. **D** Network graphs showing significant (90% and 95% CI) partial phylogenetic correlations (positive = blue edges, negative = red edges) between functional traits (green), climate (light blue) and soil (orange) niche variables. Edge widths are proportional to partial correlation coefficients. All correlation estimates derive from the same MR-PMM model fit but are grouped and plotted in separate networks for illustrative purposes.

**Figure S3.**
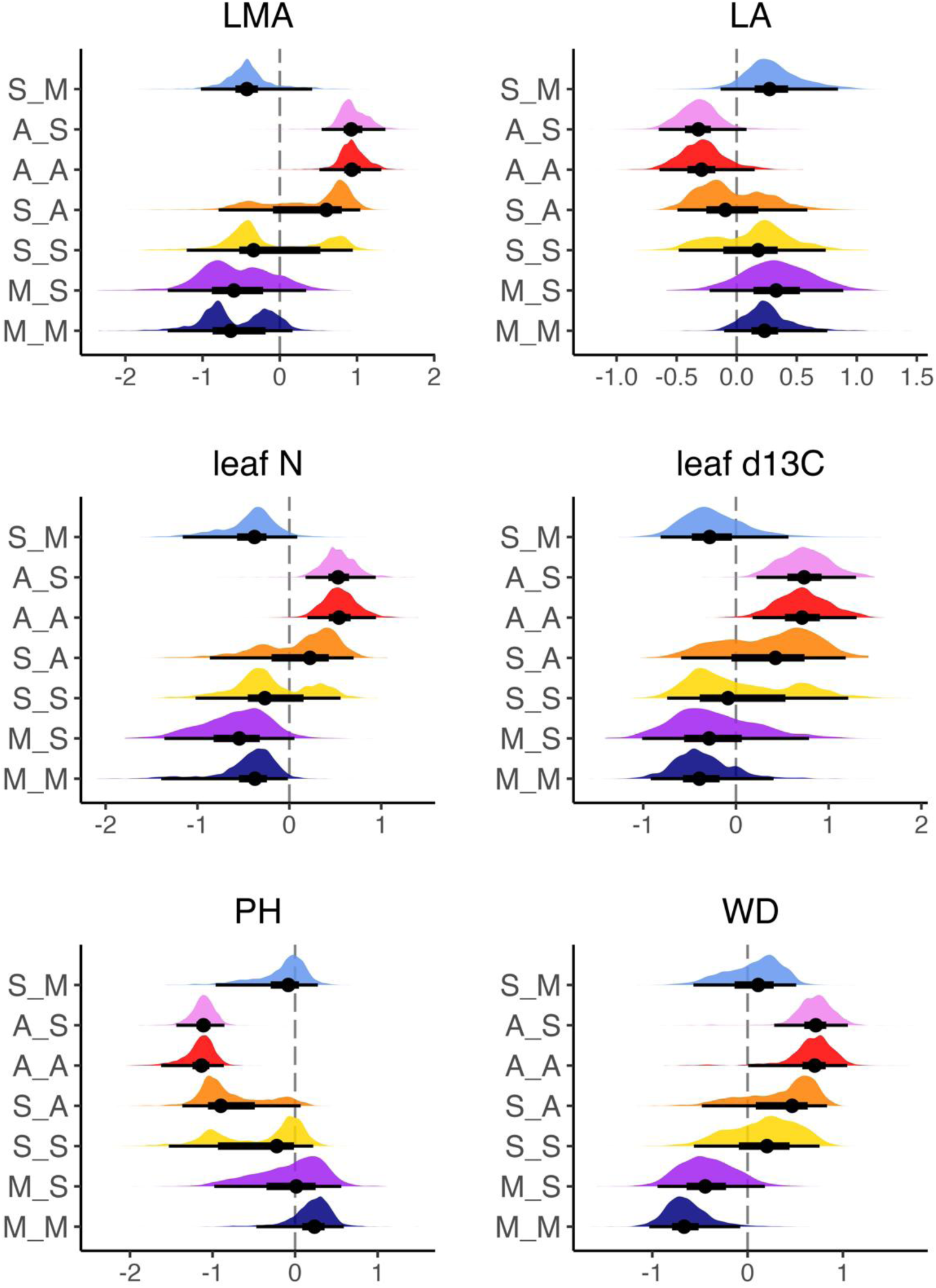
Posterior distributions of estimated trait values at internal nodes for each transition category with transition classes coloured as in figure 4. The mean (point), 50% (heavy whisker) and 95% (light whisker) CI of each posterior distribution are shown in black.

**Figure S4.**
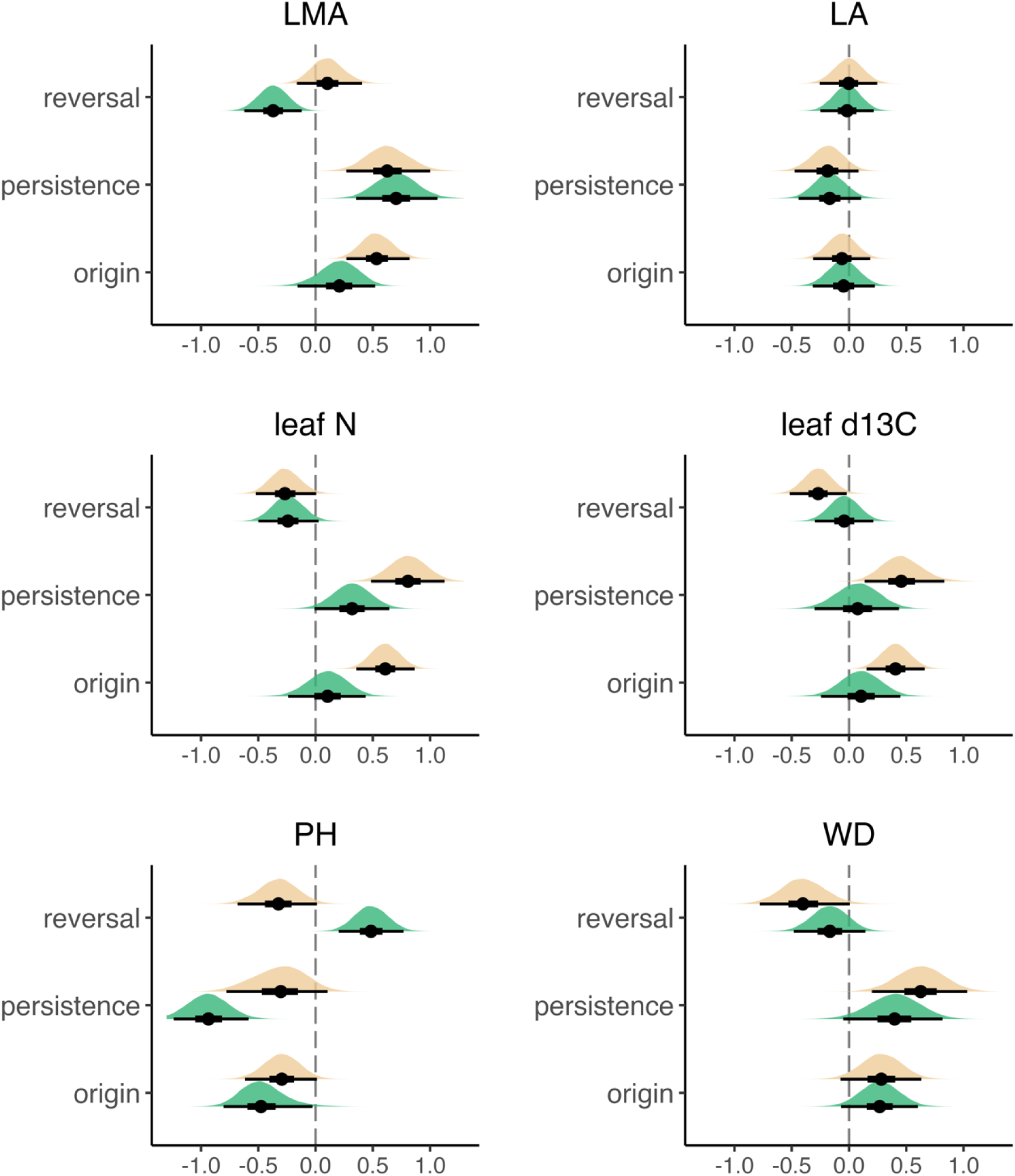
Posterior distributions of contrasts between node categories from an MR-PMM ancestral state reconstruction adjusted according to a shift model (see methods). Contrasts are computed and reported for transitions between mesic and semi-arid environments (green), as well as between semi-arid and arid environments (cream). The mean (point), 50% (heavy whisker) and 95% (light whisker) CI of each posterior distribution are shown in black.

**Figure S5.**
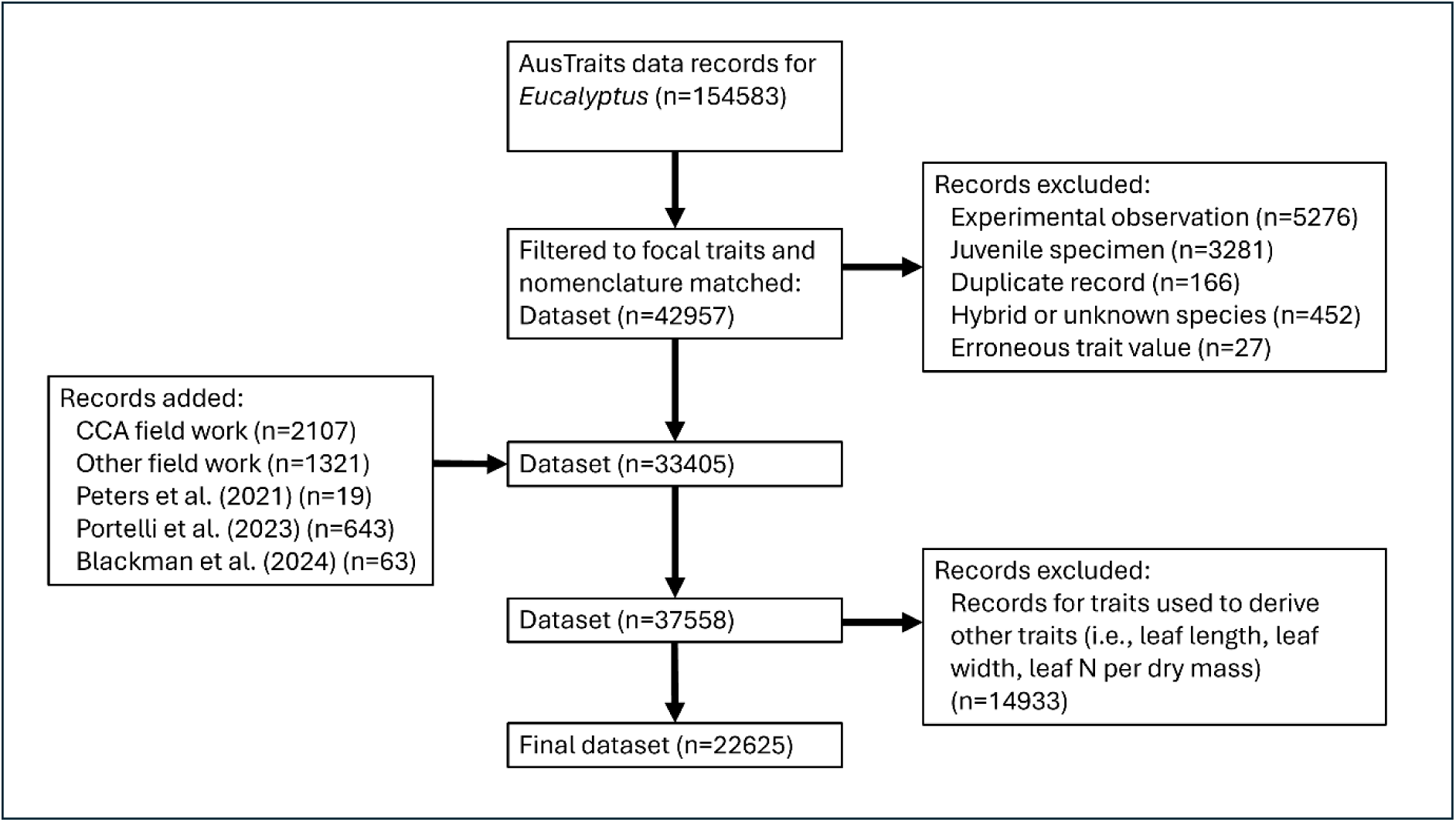
Flow diagram of data compiled for MR-PMM analyses. Data records from AusTraits^46^ were filtered to focal traits then cleaned by matching species nomenclature with the phylogeny^35^ and excluding records based on the criteria shown. Additional trait observations from targeted field work by the authors (n=3428) and a recent study^60,70,107^ not yet included in AusTraits were added. Finally, records for superfluous variables were removed.

**Figure S6.**
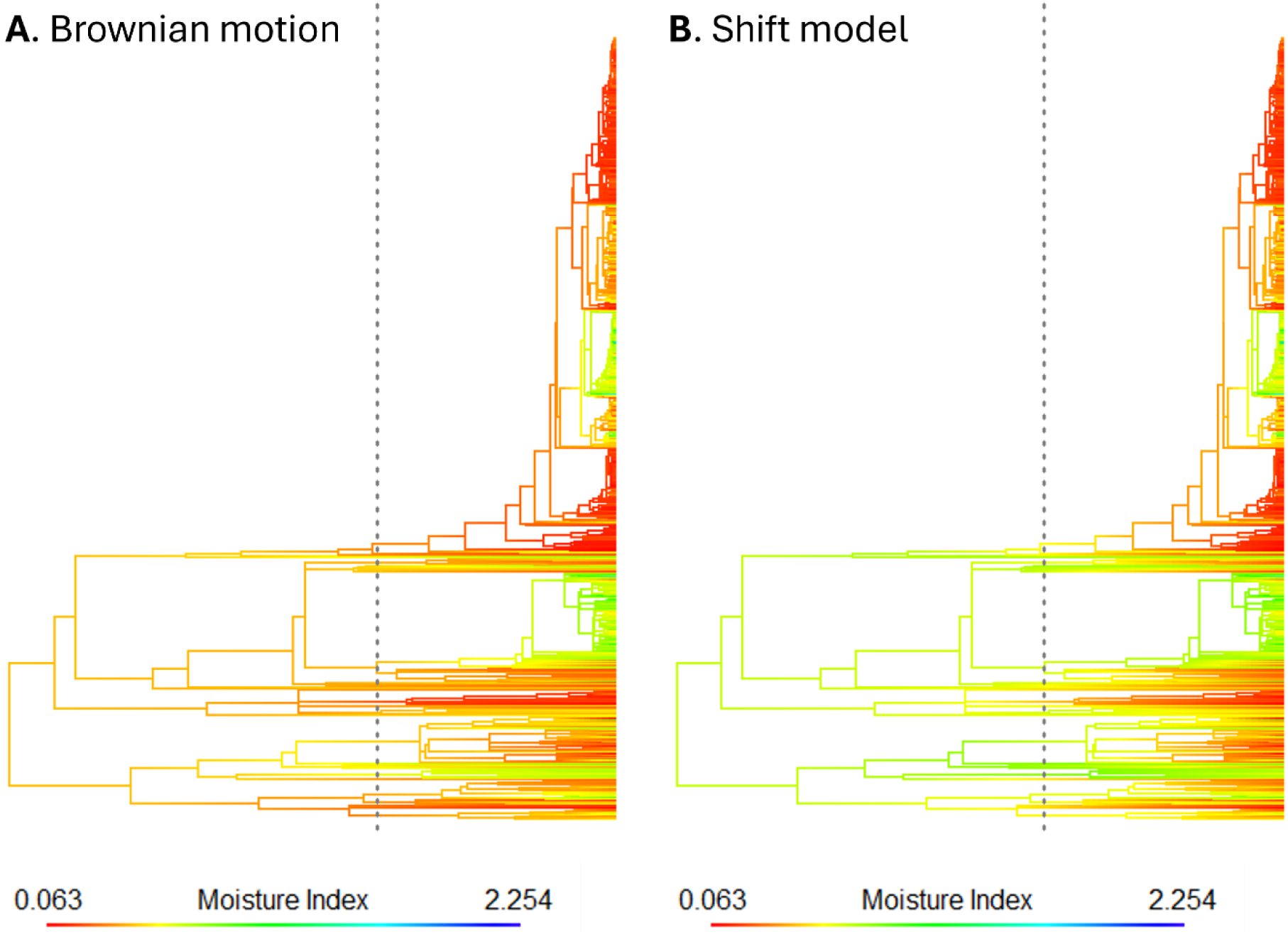
Ancestral state reconstructions of moisture index for *Eucalyptus* species according to **A** a simple Brownian motion and **B** a shift model in which the ancestral moisture niche of *Eucalyptus* is assumed to be humid (moisture index > 0.65), with a directional trend toward more arid (lower moisture index) environments from the Miocene boundary 23 Mya (dotted line).

**Figure S7.**
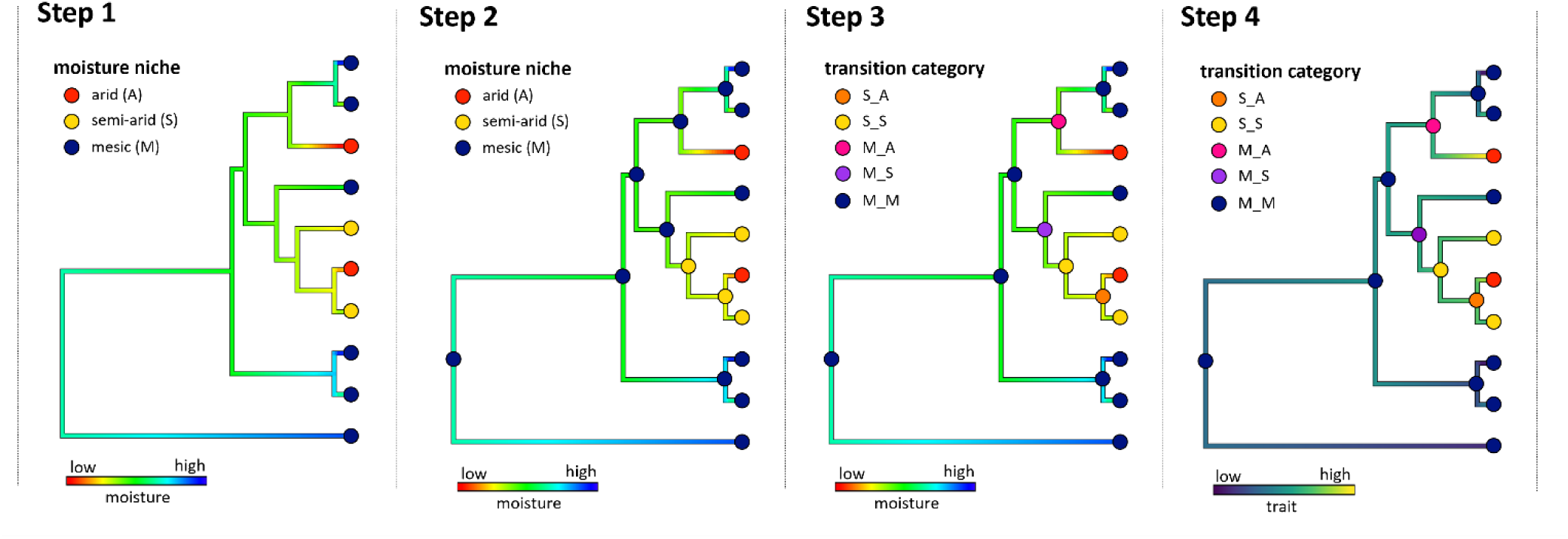
Simplified diagrammatic example of the method used to define aridity transition categories for internal nodes. In step 1, continuous values of moisture index at the tips (species mean moisture) are used to classify tips into discrete moisture niche categories and reconstruct ancestral states along internal branches. In step 2, reconstructed values are used to categorise internal nodes. In step 3, internal nodes are re-classified according to their own moisture niche state and that of their descendant (child) nodes, defining transition categories that distinguish internal nodes that did and did not give rise to more / less arid descendants. In step 4, reconstructed functional trait values are mapped onto transition categories, facilitating comparisons between transition categories in reconstructed values (figure 4B) as well as contrasts between posterior distributions of fixed effects for different transition categories (figure 4C).

**Figure S8.**
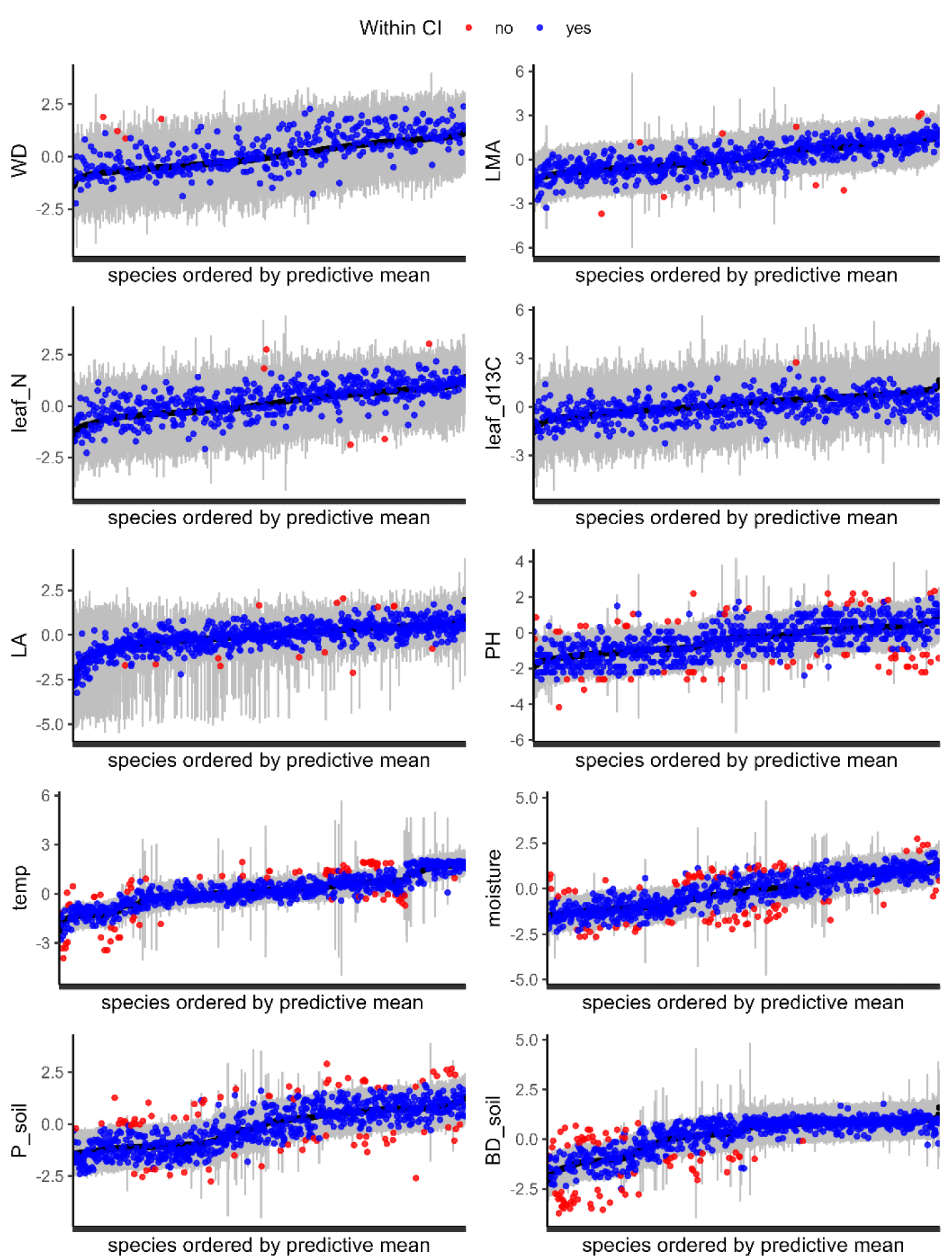
Leave-one-out (LOO) predictive checks for functional traits and environmental niche variables from MR-PMM. Three-point summaries show the mean (black points) and 0.95 CI (grey bars) of model predictions for left-out species. Predictions are made conditional on all random effects, with observations ordered by the predictive mean. Observed species mean values for each trait and niche variable are superimposed and coloured based on whether the observed value falls within the predictive interval. Species for which the model had greater predictive uncertainty (wide grey bars) are generally those with few close relatives in the phylogeny, e.g. monotypic subgenera, highlighting the importance of phylogeny for predictive performance. Species-level traits (environmental niche variables and max height) tend to show poorer predictive performance, with more observations falling outside the predictive interval. For some variables, predictive performance varies across the range of data values, e.g., poorer predictive performance at lower values of soil BD.

**Figure S9.**
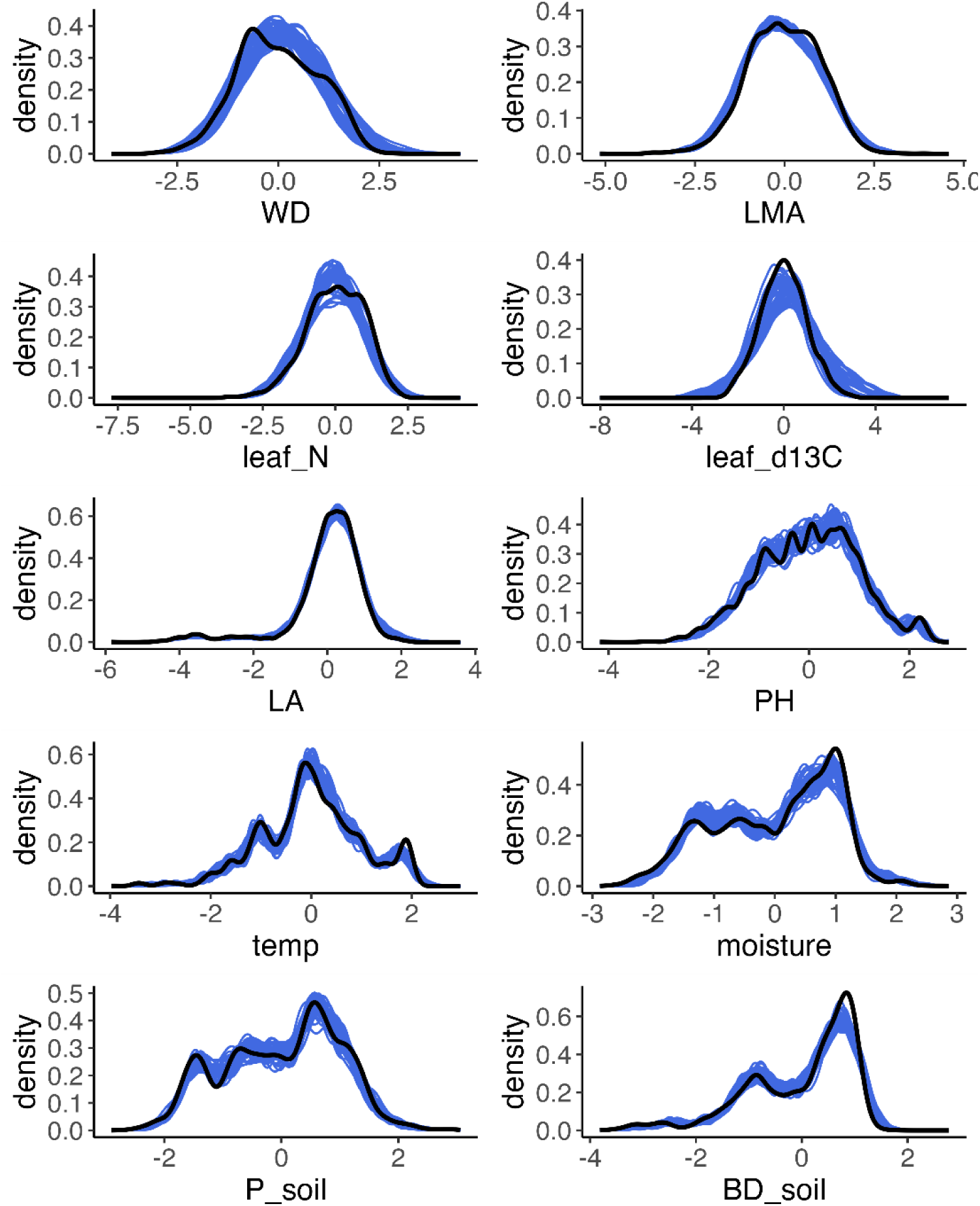
Posterior predictive checks for functional traits and environmental niche variables from MR-PMM. Posterior predictive checks compare simulated data from the fitted model (blue lines) to the observed data distribution (black line) to test both the adequacy of the fit to the data and the plausibility of model predictions.

**Figure S10.**
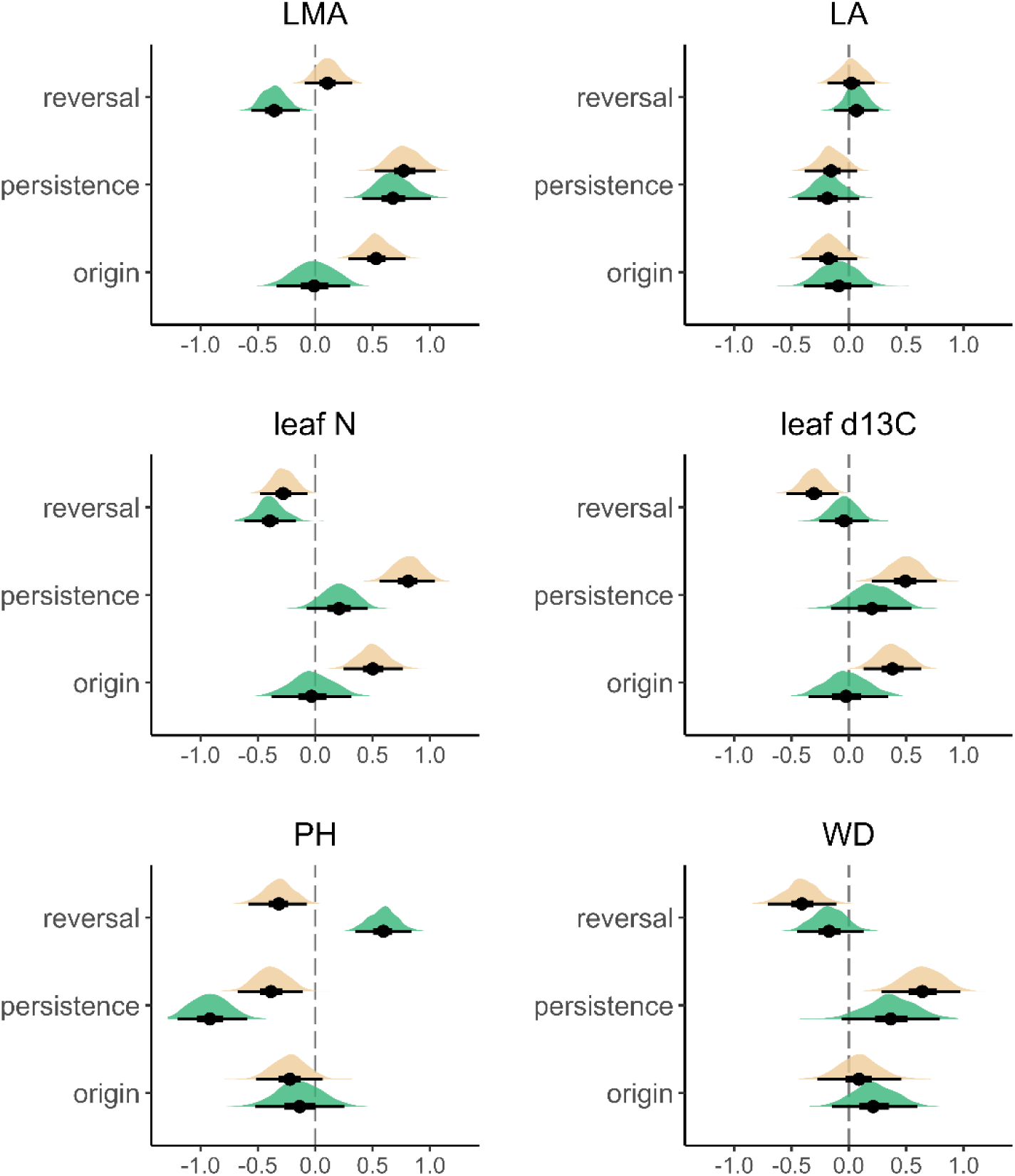
Posterior distributions of contrasts between node categories from an MR-PMM ancestral state reconstruction based on Brownian Motion. Contrasts are computed and reported for transitions between mesic and semi-arid environments (green), as well as between semi-arid and arid environments (cream). The mean (point), 50% (heavy whisker) and 95% (light whisker) CI of each posterior distribution are shown in black.

