## Supplementary material for "Conserved strategies underpin the replicated evolution of aridity tolerance in trees": data and code: ERE_ANALYSES.R

```
#------------------------------------------------------------------------#
#                                                                        #
# Data Analysis: Conserved strategies underpin the replicated evolution  #
# of aridity tolerance in trees                                          #
#                                                                        #
# Ben Halliwell                                                          #
# 2025                                                                   #
#                                                                        #
#------------------------------------------------------------------------#

## SETUP

# load packages
library(doParallel)
library(purrr)
library(MCMCglmm)
library(MASS)
library(tidyverse)
library(tidybayes)
library(ape)
library(phytools)
library(geiger)
library(igraph)
library(viridis)
library(ggpubr)
library(ggtree)
library(ggnewscale)
library(bayesplot)
library(plotrix)

# read in functions script
source("ERE_FUNCTIONS.R")

# assign select function
select <- dplyr::select

# set plot theme
theme_set(theme_classic())

#------------------------------------------------------------------------#

# TREES
phy_sp <- readRDS("euc_phy_sp.rds") # species-level topology
phy_ser_list <- readRDS("euc_phy_ser_list.rds") # list of candidate series-level topologies

# DATA
dat <- read.csv("euc_data.csv")

# subset data for different analyses (species- and series-level phylogeny). scale variables after subsetting to retain centering.
dat_sp <- dat %>% # species level data
  filter(taxon_sp %in% phy_sp$tip.label) %>%
  mutate(LA = scale(log(leaf_area))[,1],
         LMA = scale(log(leaf_mass_per_area))[,1],
         leaf_d13C = scale(log(abs(leaf_delta13C))*-1)[,1],
         leaf_N = scale(log(leaf_N))[,1],
         WD = scale(log(wood_density))[,1],
         PH = scale(log(plant_height_taxon_sp))[,1],
         moisture = scale(log(moisture_mean_taxon_sp))[,1],
         temp = scale(log(temp_mean_taxon_sp))[,1],
         BD_soil = scale(log(BD_mean_taxon_sp))[,1],
         P_soil = scale(log(P_mean_taxon_sp))[,1],
         phylo = taxon_sp)
dat_ser <- dat %>% # series level data
  filter(series %in% phy_ser_list[[1]]$tip.label) %>% 
  mutate(LA = scale(log(leaf_area))[,1],
         LMA = scale(log(leaf_mass_per_area))[,1],
         leaf_d13C = scale(log(abs(leaf_delta13C))*-1)[,1],
         leaf_N = scale(log(leaf_N))[,1],
         WD = scale(log(wood_density))[,1],
         PH = scale(log(plant_height_taxon_sp))[,1],
         moisture = scale(log(moisture_mean_taxon_sp))[,1],
         temp = scale(log(temp_mean_taxon_sp))[,1],
         BD_soil = scale(log(BD_mean_taxon_sp))[,1],
         P_soil = scale(log(P_mean_taxon_sp))[,1],
         phylo = series)

#------------------------------------------------------------------------#

## FIT MODELS

# run parameters
n_resp = 10 # number of response variables in model
n_tree = 100 # number of series-level trees to sample
n_chain = 4 # number of chains for model using species-level tree
n_cores = 4 # number of cores for parallel processing
nitts = 11000
burns = 1000
thins = 10
shrink = 1 # mild shrinkage applied to make prior uniform (uninformative) on correlation coefficients

# parameter expanded prior
p_exp <- list(G = list(G1=list(V=diag(n_resp), nu=n_resp+shrink, alpha.mu = rep(0,n_resp), alpha.V = diag(n_resp)*1000),
                       G2=list(V=diag(n_resp), nu=n_resp+shrink, alpha.mu = rep(0,n_resp), alpha.V = diag(n_resp)*1000),
                       G3=list(V = 1, nu = 0.002, alpha.mu = 0, alpha.V = 1000),
                       G4=list(V = 1, nu = 0.002, alpha.mu = 0, alpha.V = 1000),
                       G5=list(V = 1, nu = 0.002, alpha.mu = 0, alpha.V = 1000),
                       G6=list(V = 1, nu = 0.002, alpha.mu = 0, alpha.V = 1000),
                       G7=list(V = 1, nu = 0.002, alpha.mu = 0, alpha.V = 1000)),
              R = list(V=diag(n_resp), nu=n_resp, fix = 6)) # fix residual variances of species-level traits

# set up cluster to run models in parallel
result_list_ser <- list() # list for mod results
result_list_sp <- list()# list for mod results
cl <- makeCluster(n_cores)
clusterEvalQ(cl,  library(tidyverse))
clusterEvalQ(cl,  library(MCMCglmm))
registerDoParallel(cl)

#------------------------------------------------------------------------#

## SERIES LEVEL TREE

# fit dummy model
C_ser <- MCMCglmm::inverseA(phy_ser_list[[1]])$Ainv
fit_ser <- MCMCglmm(cbind(WD, LMA, leaf_N, leaf_d13C, LA, PH, temp, moisture, P_soil, BD_soil) ~ 
                                trait-1 + # fit separate intercept for each trait
                                at.level(trait,"LA"):temp_diff_taxon_sp + at.level(trait,"LA"):moisture_diff_taxon_sp + at.level(trait,"LA"):P_diff_taxon_sp + at.level(trait,"LA"):BD_diff_taxon_sp + # taxon_sp anomalies
                                at.level(trait,"LMA"):temp_diff_taxon_sp + at.level(trait,"LMA"):moisture_diff_taxon_sp + at.level(trait,"LMA"):P_diff_taxon_sp + at.level(trait,"LMA"):BD_diff_taxon_sp +
                                at.level(trait,"leaf_N"):temp_diff_taxon_sp + at.level(trait,"leaf_N"):moisture_diff_taxon_sp + at.level(trait,"leaf_N"):P_diff_taxon_sp + at.level(trait,"leaf_N"):BD_diff_taxon_sp +
                                at.level(trait,"leaf_d13C"):temp_diff_taxon_sp + at.level(trait,"leaf_d13C"):moisture_diff_taxon_sp + at.level(trait,"leaf_d13C"):P_diff_taxon_sp + at.level(trait,"leaf_d13C"):BD_diff_taxon_sp +
                                at.level(trait,"WD"):temp_diff_taxon_sp + at.level(trait,"WD"):moisture_diff_taxon_sp + at.level(trait,"WD"):P_diff_taxon_sp + at.level(trait,"WD"):BD_diff_taxon_sp,
                    random   = ~us(trait):phylo + # phylogenetic between-series covariance
                                us(trait):taxon_sp + # non-phylogenetic between-species covariance
                                idh(at.level(trait,"LA")):dataset_id + # between-study variances for relevant response variables
                                idh(at.level(trait,"LMA")):dataset_id +
                                idh(at.level(trait,"leaf_N")):dataset_id +
                                idh(at.level(trait,"leaf_d13C")):dataset_id +
                                idh(at.level(trait,"WD")):dataset_id,
                    rcov     = ~idh(trait):units, # (within-species) residual variance
                    ginv     = list(phylo = C_ser),
                    family   = rep("gaussian", n_resp),
                    nitt     = 11,
                    burnin   = 1,
                    thin     = 1,
                    data     = as.data.frame(dat_ser),
                    prior    = p_exp,
                    pr = T, pl = T, saveX = T, saveZ = T, verbose = F)
saveRDS(fit_ser, "fit_ser.rds")
# run (takes >12 hours for n_tree=100. Reduce nitts/burns/thins to reduce compute time)
{
  result_list_ser <- foreach(i=1:n_tree) %dopar% {
    C_ser <- MCMCglmm::inverseA(phy_ser_list[[i]])$Ainv
    fit_ser <- MCMCglmm(cbind(WD, LMA, leaf_N, leaf_d13C, LA, PH, temp, moisture, P_soil, BD_soil) ~ 
                                    trait-1 + # fit separate intercept for each trait
                                    at.level(trait,"LA"):temp_diff_taxon_sp + at.level(trait,"LA"):moisture_diff_taxon_sp + at.level(trait,"LA"):P_diff_taxon_sp + at.level(trait,"LA"):BD_diff_taxon_sp + # taxon_sp anomalies
                                    at.level(trait,"LMA"):temp_diff_taxon_sp + at.level(trait,"LMA"):moisture_diff_taxon_sp + at.level(trait,"LMA"):P_diff_taxon_sp + at.level(trait,"LMA"):BD_diff_taxon_sp +
                                    at.level(trait,"leaf_N"):temp_diff_taxon_sp + at.level(trait,"leaf_N"):moisture_diff_taxon_sp + at.level(trait,"leaf_N"):P_diff_taxon_sp + at.level(trait,"leaf_N"):BD_diff_taxon_sp +
                                    at.level(trait,"leaf_d13C"):temp_diff_taxon_sp + at.level(trait,"leaf_d13C"):moisture_diff_taxon_sp + at.level(trait,"leaf_d13C"):P_diff_taxon_sp + at.level(trait,"leaf_d13C"):BD_diff_taxon_sp +
                                    at.level(trait,"WD"):temp_diff_taxon_sp + at.level(trait,"WD"):moisture_diff_taxon_sp + at.level(trait,"WD"):P_diff_taxon_sp + at.level(trait,"WD"):BD_diff_taxon_sp,
                        random   = ~us(trait):phylo + # phylogenetic between-series covariance
                                    us(trait):taxon_sp + # non-phylogenetic between-species covariance
                                    idh(at.level(trait,"LA")):dataset_id + # between-study variances for relevant response variables
                                    idh(at.level(trait,"LMA")):dataset_id +
                                    idh(at.level(trait,"leaf_N")):dataset_id +
                                    idh(at.level(trait,"leaf_d13C")):dataset_id +
                                    idh(at.level(trait,"WD")):dataset_id,
                        rcov     = ~idh(trait):units, # (within-species) residual variance
                        ginv     = list(phylo = C_ser),
                        family   = rep("gaussian", n_resp),
                        nitt     = nitts,
                        burnin   = burns,
                        thin     = thins,
                        data     = as.data.frame(dat_ser),
                        prior    = p_exp,
                        pr = T, pl = T, saveX = T, saveZ = T, verbose = F)
    list(fit = list(Sol = fit_ser$Sol %>% as_tibble %>% mutate(tree = i, iter = 1:((nitts-burns)/thins)),
                    VCV = fit_ser$VCV %>% as_tibble %>% mutate(tree = i, iter = 1:((nitts-burns)/thins))),
         pars = list(n_resp = n_resp, n_tree = n_tree, nitts = nitts, burns = burns, thins = thins))
  }
}
setwd(getwd());saveRDS(result_list_ser, "result_list_ser.rds");gc()


#------------------------------------------------------------------------#

## SPECIES LEVEL TREE

# fit dummy model
C_sp <- MCMCglmm::inverseA(phy_sp)$Ainv
fit_sp <- MCMCglmm::MCMCglmm(cbind(WD, LMA, leaf_N, leaf_d13C, LA, PH, temp, moisture, P_soil, BD_soil) ~
                                         trait-1 + # fit separate intercept for each trait
                                         at.level(trait,"LA"):temp_diff_taxon_sp + at.level(trait,"LA"):moisture_diff_taxon_sp + at.level(trait,"LA"):P_diff_taxon_sp + at.level(trait,"LA"):BD_diff_taxon_sp +
                                         at.level(trait,"LMA"):temp_diff_taxon_sp + at.level(trait,"LMA"):moisture_diff_taxon_sp + at.level(trait,"LMA"):P_diff_taxon_sp + at.level(trait,"LMA"):BD_diff_taxon_sp +
                                         at.level(trait,"leaf_N"):temp_diff_taxon_sp + at.level(trait,"leaf_N"):moisture_diff_taxon_sp + at.level(trait,"leaf_N"):P_diff_taxon_sp + at.level(trait,"leaf_N"):BD_diff_taxon_sp +
                                         at.level(trait,"leaf_d13C"):temp_diff_taxon_sp + at.level(trait,"leaf_d13C"):moisture_diff_taxon_sp + at.level(trait,"leaf_d13C"):P_diff_taxon_sp + at.level(trait,"leaf_d13C"):BD_diff_taxon_sp +
                                         at.level(trait,"WD"):temp_diff_taxon_sp + at.level(trait,"WD"):moisture_diff_taxon_sp + at.level(trait,"WD"):P_diff_taxon_sp + at.level(trait,"WD"):BD_diff_taxon_sp,
                             random   = ~us(trait):phylo + 
                                         us(trait):taxon_sp +
                                         idh(at.level(trait,"LA")):dataset_id + # between-study variances for relevant response variables
                                         idh(at.level(trait,"LMA")):dataset_id +
                                         idh(at.level(trait,"leaf_N")):dataset_id +
                                         idh(at.level(trait,"leaf_d13C")):dataset_id +
                                         idh(at.level(trait,"WD")):dataset_id,
                             rcov     = ~idh(trait):units,
                             ginv     = list(phylo = C_sp),
                             family   = rep("gaussian", n_resp),
                             nitt     = 11,
                             burnin   = 1,
                             thin     = 1,
                             data     = as.data.frame(dat_sp),
                             prior    = p_exp,
                             pr = T, pl = T, saveX = T, saveZ = T, verbose = F)
saveRDS(fit_sp, "fit_sp.rds")
# run
{
  result_list_sp <- foreach(i=1:n_chain) %dopar% {
    C_sp <- MCMCglmm::inverseA(phy_sp)$Ainv
    fit_sp <- MCMCglmm::MCMCglmm(cbind(WD, LMA, leaf_N, leaf_d13C, LA, PH, temp, moisture, P_soil, BD_soil) ~
                                   trait-1 + # fit separate intercept for each trait
                                   at.level(trait,"LA"):temp_diff_taxon_sp + at.level(trait,"LA"):moisture_diff_taxon_sp + at.level(trait,"LA"):P_diff_taxon_sp + at.level(trait,"LA"):BD_diff_taxon_sp +
                                   at.level(trait,"LMA"):temp_diff_taxon_sp + at.level(trait,"LMA"):moisture_diff_taxon_sp + at.level(trait,"LMA"):P_diff_taxon_sp + at.level(trait,"LMA"):BD_diff_taxon_sp +
                                   at.level(trait,"leaf_N"):temp_diff_taxon_sp + at.level(trait,"leaf_N"):moisture_diff_taxon_sp + at.level(trait,"leaf_N"):P_diff_taxon_sp + at.level(trait,"leaf_N"):BD_diff_taxon_sp +
                                   at.level(trait,"leaf_d13C"):temp_diff_taxon_sp + at.level(trait,"leaf_d13C"):moisture_diff_taxon_sp + at.level(trait,"leaf_d13C"):P_diff_taxon_sp + at.level(trait,"leaf_d13C"):BD_diff_taxon_sp +
                                   at.level(trait,"WD"):temp_diff_taxon_sp + at.level(trait,"WD"):moisture_diff_taxon_sp + at.level(trait,"WD"):P_diff_taxon_sp + at.level(trait,"WD"):BD_diff_taxon_sp,
                                 random   = ~us(trait):phylo + 
                                   us(trait):taxon_sp +
                                   idh(at.level(trait,"LA")):dataset_id + # between-study variances for relevant response variables
                                   idh(at.level(trait,"LMA")):dataset_id +
                                   idh(at.level(trait,"leaf_N")):dataset_id +
                                   idh(at.level(trait,"leaf_d13C")):dataset_id +
                                   idh(at.level(trait,"WD")):dataset_id,
                                 rcov     = ~idh(trait):units,
                                 ginv     = list(phylo = C_sp),
                                 family   = rep("gaussian", n_resp),
                                 nitt     = nitts,
                                 burnin   = burns,
                                 thin     = thins,
                                 data     = as.data.frame(dat_sp),
                                 prior    = p_exp,
                                 pr = T, pl = T, saveX = T, saveZ = T, verbose = F)
    list(fit = list(Sol = fit_sp$Sol %>% as_tibble %>% mutate(chain = i, iter = 1:((nitts-burns)/thins)),
                    VCV = fit_sp$VCV %>% as_tibble %>% mutate(chain = i, iter = 1:((nitts-burns)/thins))),
         pars = list(n_resp = n_resp, n_chain = n_chain, nitts = nitts, burns = burns, thins = thins))
  }
}
setwd(getwd());saveRDS(result_list_sp, "result_list_sp.rds");gc()


#---------------------------LOAD MODEL------------------------------#

# Models above were fit specifying phylogenetic effects at two different taxonomic levels (either the 
# series level [fit_ser] or species level [fit_sp]) to assess the influence of phylogenetic structure 
# on parameter estimates. The series level analyses are preferable because they account for phylogenetic 
# uncertainty and allow data on additional species to be included (see methods), though species-level
# analyses may provide better resolution for species-level traits (i.e., max height).
# To generate results from the workflow below, choose which model to assess by running the relevant chunk below:

# load save fits
fit_ser <- readRDS("fit_ser.rds")
result_list_ser <- readRDS("result_list_ser.rds")
fit_sp <- readRDS("fit_sp.rds")
result_list_sp <- readRDS("result_list_sp.rds")

## SERIES LEVEL
# combine samples across tree fits
result_list <- result_list_ser
res_Sol <- result_list %>% purrr::map("fit") %>% purrr::map_dfr(~ .x[['Sol']])
res_VCV <- result_list %>% purrr::map("fit") %>% purrr::map_dfr(~ .x[['VCV']])
res <- dplyr::bind_cols(dplyr::select(res_Sol, -tree, -iter), res_VCV)
# fill dummy model with samples
fit_ser$Sol <- res_Sol %>% dplyr::select(-c(tree,iter)) %>% slice_sample(n=10000) %>% as.mcmc # subsample posterior for model object to reduce compute (DOES NOT CHANGE RESULT)
fit_ser$VCV <- res_VCV %>% dplyr::select(-c(tree,iter)) %>% slice_sample(n=10000) %>% as.mcmc # subsample posterior for model object to reduce compute (DOES NOT CHANGE RESULT)
# assign fit_ser as focal model
mod <- fit_ser
mod_dat <- dat_ser

# ## SPECIES LEVEL
# # combine samples across chains
# result_list <- result_list_sp
# res_Sol <- result_list %>% purrr::map("fit") %>% purrr::map_dfr(~ .x[['Sol']])
# res_VCV <- result_list %>% purrr::map("fit") %>% purrr::map_dfr(~ .x[['VCV']])
# res <- dplyr::bind_cols(dplyr::select(res_Sol, -chain, -iter), res_VCV)
# # fill dummy model with samples
# fit_sp$Sol <- res_Sol %>% select(-c(chain,iter)) %>% as.mcmc
# fit_sp$VCV <- res_VCV %>% select(-c(chain,iter)) %>% as.mcmc
# # assign fit_sp as focal model
# mod <- fit_sp
# mod_dat <- dat_sp

## load in run parameters
res_pars <- result_list[[1]]$pars
n_tree = res_pars$n_tree
n_resp = res_pars$n_resp
n_chain = res_pars$n_chain
nitts = res_pars$nitts
burns = res_pars$burns
thins = res_pars$thins

#------------------------------------------------------------------------#

## CONVERGENCE DIAGNOSTICS

# CHECK 1 - compute scale-reduction factors (R hat) to assess convergence of combined samples
res_VCV %>% 
  dplyr::select(!any_of(c("tree", "chain", "iter"))) %>%
  as.data.frame() %>% 
  map_dbl(rstan::Rhat) %>% .[!is.na(.)] %>% {.[.>1.01]}

# CHECK 2 - assess whether certain trees/chains give convergence issues using R hat (takes a few minutes)
bad_trees <- res %>%
  group_by(tree) %>% 
  # group_by(chain) %>% # species-level tree
  summarise_all(~posterior::rhat(.)) %>% 
  mutate(row_mean = rowMeans(across(setdiff(everything(), any_of(c("tree","chain","iter")))))) %>%
  filter(row_mean > 1.005) %>% # filter to tree fits with poor mean R hat across all parameters
  pull(tree)
bad_trees # none detected

# CHECK 3 - compare parameter estimates across trees (fit_ser) or chains (fit_sp) by plotting samples 
# from each tree/chain as a density overlay
var_comp = "traitPH:traitmoisture.phylo" # choose variance component you wish to assess
y <- res %>% pull(var_comp) %>% sample(., ((nitts-burns)/thins)) # draw samples from combined posterior to make y
yrep <- res %>% dplyr::select(any_of(c(var_comp,"tree", "chain", "iter"))) %>%
  pivot_wider(names_from = iter, values_from = any_of(var_comp)) %>%
  dplyr::select(!any_of(c("tree", "chain"))) %>% as.matrix
ppc_dens_overlay(y = y, yrep = yrep, size = 0.75) # plot param dist from each tree/chain fit as separate density

## NOTE
# 2 out of 100 trees (38 and 58) produce markedly stronger estimates for trait correlations
yrep %>% rowMeans() %>% hist(breaks=30)
# however excluding samples from models fit to these trees (code below) DOES NOT CHANGE RESULTS

# # exclude outlier trees
# res_Sol <- res_Sol %>% filter(!tree%in%c(38,58));gc()
# res_VCV <- res_VCV %>% filter(!tree%in%c(38,58));gc()
# res <- res %>% filter(!tree%in%c(38,58));gc()

#------------------------------------------------------------------------#

## MODEL VALIDATION

## POSTERIOR PREDICTIVE CHECKS
# perform pp checks to assess whether the model simulates data with a marginal distribution
# similar to the real data (i.e., assess whether model predictions are plausible)
resp_names <- c("WD","LMA","leaf_N","leaf_d13C","LA","PH","temp","moisture","P_soil","BD_soil") # must match order of model fit
result <- ppc(mod, mod_dat, n_resp, resp_names, n_samples = 30)
# create and display plots
plot_list <- create_ppc_plots(result, mod_dat)
xlab = resp_names
plot_list <- lapply(seq_along(plot_list), function(i) {
  plot_list[[i]]$labels$x <- xlab[i]
  plot_list[[i]]
})
plot_list <- lapply(plot_list, function(plot) {
  plot + theme(
    text = element_text(size = 14),
    axis.text = element_text(size = 12), 
    axis.title = element_text(size = 14), 
    legend.text = element_text(size = 12), 
    legend.title = element_text(size = 14)
  )
})
ggpubr::ggarrange(
  plot_list[[1]],plot_list[[2]],
  plot_list[[3]],plot_list[[4]],
  plot_list[[5]],plot_list[[6]],
  plot_list[[7]],plot_list[[8]],
  plot_list[[9]],plot_list[[10]],
  nrow = 5, ncol = 2)

#--------------------------------------------------------#

## LOO-CV
# perform leave-one-out (LOO) cross validation (CV) to assess model predictive performance.
# We preform LOO-CV by leaving out all observations for a single species with each iteration.

# # load saved runs
# loo_pred_fit_ser <- readRDS("loo_pred_fit_ser.rds")

# run parameters
trees = phy_ser_list
# trees = list(phy_sp) # for species level analyses [fit_sp]
resp_names = c("WD","LMA","leaf_N","leaf_d13C","LA","PH","temp","moisture","P_soil","BD_soil")
n_species = length(unique(mod_dat$taxon_sp)) # perform LOO over all species
nitt = 3300
burnin = 300
thin = 3
n_cores = 4

## RUN LOO (takes >24 hours. Reduce nitt/burnin/thin to reduce compute time)
loo_pred_fit_ser <- loo_cv_species(fit = mod, data = mod_dat, trees = trees, prior = p_exp, n_species = n_species,
                                   part.1 = "phylo", part.2 = "taxon_sp", calc_cor = TRUE, res_cor = FALSE, 
                                   family = rep("gaussian", n_resp), n_resp = n_resp, resp_names = resp_names,
                                   cores = n_cores, nitt = nitt, burnin = burnin, thin = thin)
saveRDS(loo_pred_fit_ser, "loo_pred_fit_ser.rds")


## VALIDATE CORRELATIONS

# CHECK 1 - combining posterior samples across LOO fits when inferring partial phylogenetic  
# correlations returns equivalent results with only slightly greater uncertainty
loo_cor <- purrr::map(loo_pred_fit_ser, "par_phy_cor") %>% bind_rows() %>% as.matrix() %>% as_tibble() 
ordered_indices <- order(apply(loo_cor, 2, mean));ordered_phy_cor <- loo_cor[, ordered_indices] # reorder by posterior mean
ordered_phy_cor %>% mcmc_intervals(prob_outer = 0.95) # plot

# CHECK 2 - comparing parameter estimates across LOO fits shows inferences are robust to cross validation
cor_est = "moisture:ph" # choose partial correlation coefficient to assess
loo_cor <- loo_cor %>% mutate(loo_id = rep(1:n_species, each = (nitt-burnin)/thin), iter = rep(1:((nitt-burnin)/thin), n_species))
y <- loo_cor %>% pull(cor_est) %>% sample(., ((nitt-burnin)/thin)) # draw samples from combined posterior to make y
yrep <- loo_cor %>% dplyr::select(any_of(cor_est), loo_id, iter) %>%
  pivot_wider(names_from = iter, values_from = any_of(cor_est)) %>% 
  dplyr::select(-loo_id) %>% as.matrix
ppc_dens_overlay(y = y, yrep = yrep, size = 0.75) # plot param dist from each tree/chain fit as separate density


## PLOT DATA OVER LOO-CV PREDICTION INTERVALS
pred_plot_1 <- purrr::map(loo_pred_fit_ser, "predictive_interval") %>% bind_rows() %>% filter(trait %in% resp_names[1:5])
pred_plot_2 <- purrr::map(loo_pred_fit_ser, "confidence_interval") %>% bind_rows() %>% filter(trait %in% resp_names[6:10]) # use confidence interval for species level traits because no residual variance
pred_plot <- bind_rows(pred_plot_1, pred_plot_2) %>% mutate(taxon_sp = species)
dat_order <- mod_dat %>% 
  dplyr::select(c(taxon_sp, all_of(resp_names))) %>%
  filter(taxon_sp %in% pred_plot$taxon_sp) %>% 
  group_by(taxon_sp) %>%
  summarise_if(is.numeric, mean, na.rm = TRUE)
pred_plot <- pred_plot %>% left_join(dat_order) %>% 
  left_join(mod_dat %>% dplyr::select(taxon_sp, series, subgenus) %>% distinct) %>% 
  mutate(pred_range = upr - lwr)
# plot
plot_list <- list()
for (i in seq_along(resp_names)) {
  plot_list[[i]] <- pred_plot %>%
    filter(trait == resp_names[i]) %>%
    mutate(in_pred = as.factor(ifelse(.data[[resp_names[i]]] > lwr & .data[[resp_names[i]]] < upr, "yes", "no"))) %>%
    drop_na(all_of(resp_names[i])) %>%
    arrange(all_of(resp_names[i])) %>%
    arrange(mean) %>%
    ggplot(aes(x = factor(species, levels = unique(species)), y = mean)) +
    geom_errorbar(aes(ymin = lwr, ymax = upr), width = 0, col = "gray75") +
    geom_point(size = 1) +
    geom_point(aes(y = .data[[resp_names[i]]], col = in_pred), size = 1, alpha = 0.8) +
    scale_color_manual(values = c("red", "blue")) +
    ylab(resp_names[i]) +
    xlab("species ordered by predictive mean") +
    theme(axis.text.x = element_blank()) +
    labs(colour = "Within CI")
}
ggpubr::ggarrange(
  plot_list[[1]],plot_list[[2]],
  plot_list[[3]],plot_list[[4]],
  plot_list[[5]],plot_list[[6]],
  plot_list[[7]],plot_list[[8]],
  plot_list[[9]],plot_list[[10]],
  nrow = 5, ncol = 2,
  common.legend = T)

#-------------------------GAP FILL DATA-------------------------------#

## GAP FILL DATA TO PERFORM PCA AND PLOT TRAITS AGAINST TREE

# predict from model (takes a few minutes)
pred <- predict(fit_ser, marginal = NULL)
pred <- split(pred, ceiling(seq_along(pred)/(nrow(pred)/n_resp))) %>% as_tibble
names(pred) <- c("WD", "LMA", "leaf_N", "leaf_d13C", "LA", "PH", "temp", "moisture", "P_soil", "BD_soil")

# create gap filled data set by replacing missing values (NA's in data) with model predictions
dat_gf <- dat_ser %>% 
  mutate(LA = ifelse(is.na(LA), pred$LA, LA),
         LMA = ifelse(is.na(LMA), pred$LMA, LMA),
         leaf_N = ifelse(is.na(leaf_N), pred$leaf_N, leaf_N),
         leaf_d13C = ifelse(is.na(leaf_d13C), pred$leaf_d13C, leaf_d13C),
         PH = ifelse(is.na(PH), pred$PH, PH),
         WD = ifelse(is.na(WD), pred$WD, WD),
         temp = ifelse(is.na(temp), pred$temp, temp),
         moisture = ifelse(is.na(moisture), pred$moisture, moisture),
         P_soil = ifelse(is.na(P_soil), pred$P_soil, P_soil),
         BD_soil = ifelse(is.na(BD_soil), pred$BD_soil, BD_soil))

#-----------PCA AND TREE PLOT WITH GAP-FILLED DATA---------------------#

## PCA
vars <- c("taxon_sp","subgenus","WD", "LMA", "leaf_N", "leaf_d13C", "LA", "PH", "temp", "moisture", "P_soil", "BD_soil")
d.pca <- dat_gf %>% select(all_of(vars))
names(d.pca) <- c("taxon_sp","subgenus","wood density", "LMA", "leaf N per area", "leaf d13C", "leaf area", "max height","temperature", "moisture", "soil P", "soil bulk density")
pca.d<-d.pca[,-c(1,2)]
pca.1 <- prcomp(pca.d, center = TRUE, scale = TRUE)
summary(pca.1); pca.1$rotation[,1:4]
# plot
p <- factoextra::fviz_pca_var(pca.1, 
                              arrowsize = 1.5,
                              col.var = "cos2", col.circle = "transparent",
                              gradient.cols = c("#00AFBB", "#E7B800", "#FC4E07"), 
                              repel = T) +
  labs(x="PC1 (49.3%)", y="PC2 (17.4%)") +
  theme_classic() +
  theme(legend.position = "top",
        legend.title = element_text(vjust = 0.85, size = 16),
        legend.text = element_text(size = 12),
        axis.title = element_text(size = 16),
        axis.text = element_text(size = 16),
        plot.margin = margin(0.5,2,0.5,0.5, "cm"),
        plot.title = element_blank())
p$labels$colour<-'contribution to variance'
# re-arrange plot layers to get arrows on top of points
p$layers[4:5] <- NULL
p$layers[3:4] <- NULL
p1 <- p + geom_point(data = data.frame(subgenus = d.pca$subgenus, pca.1$x), 
                     aes(x = PC1*0.15, y = PC2*0.15), col = "grey50", size = 2, alpha = 0.2)
layers <- p1$layers
p1$layers[[1]] <- layers[[3]]
p1$layers[[3]] <- layers[[1]]
p1


## PLOT DATA AGINST TREE
# make colour scale
cols = gg_color_hue(7)
# order of traits for matrix plot
mod_traits <- c("LA", "PH", "LMA", "leaf_N","leaf_d13C", "WD", "moisture", "P_soil","temp", "BD_soil")
# summarise data by series
series_mean <- dat_gf %>% as_tibble %>% 
  group_by(series) %>% 
  summarise(across(all_of(mod_traits), ~ .x %>% mean(na.rm = TRUE))) %>% 
  left_join(., dat_gf %>% select(series, section, subgenus) %>% distinct)
# prep trait data frame
dat_cc_ser <- series_mean
X <- dat_cc_ser %>% ungroup() %>% select(all_of(mod_traits)) %>% as.matrix()
rownames(X) <- dat_cc_ser %>% pull(series)
phy_cc <- phy_ser_list[[1]]
# group by operational taxonomic unit (OTU)
grp <- list(Eucalyptus = dat_cc_ser[which(dat_cc_ser$subgenus == 'Eucalyptus'), ]$series,
            Eudesmia = dat_cc_ser[which(dat_cc_ser$subgenus == 'Eudesmia'), ]$series,
            Symphyomyrtus = dat_cc_ser[which(dat_cc_ser$subgenus == 'Symphyomyrtus'), ]$series,
            Corymbia = dat_cc_ser[which(dat_cc_ser$subgenus == 'Corymbia'), ]$series,
            Angophora = dat_cc_ser[which(dat_cc_ser$subgenus == 'Angophora'), ]$series,
            Blakella = dat_cc_ser[which(dat_cc_ser$subgenus == 'Blakella'), ]$series,
            Other = dat_cc_ser[which(dat_cc_ser$subgenus %in% c('Idiogenes','Acerosae','Cruciformes',
                                                                'Alveolata','Cuboidea')), ]$series)
# build plot
p <- ggtree(phy_cc, layout = 'rectangular', size = 0.85) + 
  scale_color_manual(values=c("#53B400",cols))
p <- groupOTU(p, grp, 'subgenus') + aes(color=subgenus)
# add heatmap
p1 <- gheatmap(p, X, offset=0, width=0.75, color=NA,
               colnames=T, colnames_position = "top", 
               colnames_offset_x = 0, colnames_offset_y = 0.5, colnames_angle = 60, 
               custom_column_labels = c("leaf area", "max height", "LMA", "leaf N", "leaf d13C", "wood desnity", "moisture", "soil P","temperature", "soil BD"),
               font.size = 4, hjust = 0) +
  scale_fill_viridis_c(option="D", name="", na.value="grey90") +
  guides(color="none", fill="none") +
  ylim(0, 130)
# add scatterplots of series mean trait values ordered my moisture index
dat_plot <- series_mean %>% pivot_longer(2:11, names_to = "trait", values_to = "value")
order <- dat_plot %>% filter(trait == "moisture") %>% arrange(value) %>% pull(series)
p2 <- dat_plot %>%
  mutate(series = factor(series, levels = order),
         subgenus = ifelse(subgenus %in% c("Idiogenes","Acerosae","Cruciformes","Alveolata","Cuboidea"), "Other", subgenus),
         trait = factor(trait, levels = c("LA","LMA","leaf_N","leaf_d13C", "WD", "PH", "moisture", "P_soil","temp", "BD_soil"))) %>% 
  ggplot(., aes(x = series,
                y = value,
                col = subgenus)) +
  geom_point(size = 2.5, alpha = 1) +
  scale_color_manual(values=cols) +
  facet_wrap(~trait, ncol = 2, scales= "free") +
  theme(axis.text.x = element_blank(),
    axis.ticks.x = element_blank(),
    axis.title.x = element_text(size=12),
    legend.text = element_text(size=10),
    legend.title = element_text(size=12),
    strip.background = element_blank(),
    strip.text = element_text(size = 12),
    plot.margin = margin(0.2,0.2,0.2,0.2, "cm")) +
  ylab("standardized mean value") + xlab("taxonomic series")
# plot
ggarrange(p1,p2)

#-------------------------INFERENCE-------------------------------#

## FIXED EFFECTS
# plot posterior estimates of fixed effects for site-level environmental anomalies (deviations from 
# species-mean environmental conditions) for each response variable

# extract estimates
res_fe <- res %>% dplyr::select(contains('at.level(trait, "WD")') |
                           contains('at.level(trait, "LMA")') |
                           contains('at.level(trait, "leaf_N")') |
                           contains('at.level(trait, "leaf_d13C")') |
                           contains('at.level(trait, "LA")')) %>% 
  dplyr::select(!(contains("dataset_id")))

# build plot
res_fe %>% as_tibble %>% 
  mutate(.chain = 1, .iteration = 1:nrow(res)) %>% 
  pivot_longer(1:20, names_to = "parameter", values_to = "estimate") %>%
  mutate(parameter = factor(rep(c("temperature", "moisture","soil P", "soil bulk density"),nrow(res)*5),
                            levels = c("temperature", "moisture","soil P", "soil bulk density")),
         env = rep(c("climate", "climate","soil", "soil"),nrow(res)*5),
         trait = rep(rep(c("WD","LMA","leaf_N","leaf_d13C","LA"),each=4),nrow(res))) %>% 
  ggplot(aes(x = estimate, 
             y = factor(trait, levels=c("WD","LMA","leaf_N","leaf_d13C","LA")))) +
  geom_vline(xintercept=0, linetype="longdash", col="gray30") +
  stat_halfeye(aes(fill=env),
               normalize = "groups", scale = 0.5, alpha = 0.8, fatten_point = 2.25) +
  theme(axis.text = element_text(size=10),
        axis.title.x = element_text(size=20),
        legend.text = element_text(size=18),
        legend.title = element_text(size=20),
        strip.background = element_blank(),
        strip.text = element_text(size = 16),
        plot.margin = margin(0.2,0.2,0.2,0.2, "cm"),
        panel.spacing = unit(1.5, "lines")) +
  scale_fill_manual(values=c("lightblue","orange")) +
  ylab("") + xlab("") + 
  facet_grid(~parameter, scale = "free_x")

#------------------------------------------------------#

## VARIANCE PARTITIONING
# Decompose the variance in each response variable across levels in the model hierarchy. These include
# phylogenetic and non-phylogenetic between-species effects, study effects, and residual errors.
# Variance explained by fixed effects is calculated using partial R2 (Nakagawa and Schielzeth 2013).

# calculate variance decomposition
fit <- mod
sample <- sample(1:nrow(fit$VCV), 1000) # subsample posteriors to reduce compute time (DOES NOT CHANGE RESULT)
fit$VCV <- fit$VCV[sample,] %>% as.mcmc
fit$Sol <- fit$Sol[sample,] %>% as.mcmc
resp_names <- c("WD","LMA","leaf_N","leaf_d13C","LA","PH","temp","moisture","P_soil","BD_soil") # must match order of model fit
levels <- c("phylo","taxon_sp","dataset_id","units") # change to taxon_sp for fit_ser_taxon_sp
res_var_list <- var_decomp(fit, mod_dat, resp_names, levels, fixed_effects = TRUE, species_cor = TRUE, res_cor = FALSE, NA_fixed = FALSE)
res_var <- res_var_list[[1]] # extract posterior means for each component

# plot
plot_var <- res_var %>% 
  filter(!component %in% c("full","random", "units")) %>% 
  mutate(component = ifelse(trait %in% c("PH", "temp", "moisture", "P_soil", "BD_soil") & 
                              component == "taxon", "residual", 
                            ifelse(trait %in% c("PH", "temp", "moisture", "P_soil", "BD_soil") & 
                                     component == "taxon_sp", "residual", component)),
         component = gsub("phylo","phylogenetic",component),
         component = gsub("taxon_sp","non-phylogenetic",component),
         component = gsub("taxon","non-phylogenetic",component),
         component = gsub("dataset_id","study",component)) 
ggplot(plot_var, 
       aes(x = factor(trait, levels=c("LA", "LMA", "leaf_N", "leaf_d13C", "WD", "PH",
                                      "temp", "moisture", "P_soil", "BD_soil")),
           y = post.mean, 
           fill = factor(component, 
                         levels=rev(c("phylogenetic","non-phylogenetic","study","fixed","residual"))))) +
  geom_bar(position="fill", stat="identity") +
  scale_fill_manual(values=rev(c("#173f5f","#20639b","#3caea3","#f6d55c","#ed553b"))) +
  guides(fill=guide_legend("")) +
  theme(axis.text = element_text(size=14),
        axis.title.x = element_text(size=18),
        legend.text= element_text(size=16)) +
  ylab("proportion of variance") + xlab("") + 
  coord_flip()

#------------------------------------------------------------------------#

## CORRELATIONS
# calculate phylogenetic (phy) and non-phylogenetic (ind) correlations between response variables. The
# function level_cor_vcv() calculates both standard and partial (par) correlations from the fitted VCV. 
# Choose which component you would like to plot results for by selecting a named element of 'cor_list'.

# calculate
cor_list <- level_cor_vcv(mod$VCV, n_resp, part.1 = "phylo", part.2 = "taxon_sp")

# choose component to plot
component = "phy"
component = "par_phy"
component = "ind"
component = "par_ind"

# PLOT INTERVALS
cor_list$posteriors[[paste0(component, "_cor")]] %>% mcmc_intervals(prob_outer = 0.95) + xlim(-1,1)

# PLOT NETWORK
cor <- paste0(component, "_cor")
mat <- paste0(component, "_mat")
sig.95 <- cor_list$posteriors[[cor]] %>% summarise_all(~ quantile(., prob=c(0.025,0.975)) %>% {. > 0} %>% table %>% length == 1) %>% t() %>% as_tibble %>% mutate(sig.95 = V1, edge = 1:((n_resp^2-n_resp)/2)) %>% dplyr::select(-V1)
sig.90 <- cor_list$posteriors[[cor]] %>% summarise_all(~ quantile(., prob=c(0.05,0.95)) %>% {. > 0} %>% table %>% length == 1) %>% t() %>% as_tibble %>% mutate(sig.90 = V1, edge = 1:((n_resp^2-n_resp)/2)) %>% dplyr::select(-V1)
sig <- left_join(sig.95, sig.90, by = "edge") %>% mutate(mean = cor_list$posteriors[[cor]] %>% summarise_all(~ mean(.)) %>% t %>% as.vector)
sig <- sig %>% mutate(sig = ifelse(sig.90 == T | sig.95==T, T, F),
                      col = ifelse(sig.90 == T & sig.95 == T & mean > 0, "blue", "red"),
                      lty = ifelse(sig.90 == T & sig.95 != T & mean > 0, "solid",
                                   ifelse(sig.90 == T & sig.95 == T & mean > 0, "solid",
                                          ifelse(sig.90 == T & sig.95 != T & mean < 0, "solid", "solid"))),
                      alpha = ifelse(sig.90 == T & sig.95 != T & mean > 0, 0.5,
                                     ifelse(sig.90 == T & sig.95 == T & mean > 0, 1,
                                            ifelse(sig.90 == T & sig.95 != T & mean < 0, 0.5, 1))))
cor_mat <- cor_list$matrices[[mat]]
dimnames(cor_mat)[[1]] <- c("wood\ndensity","LMA","leaf N\n per area","leaf\nd13C","leaf\narea","max\nheight","temperature","moisture","soil P","soil bulk\ndensity")
dimnames(cor_mat)[[2]] <- c("wood\ndensity","LMA","leaf N\n per area","leaf\nd13C","leaf\narea","max\nheight","temperature","moisture","soil P","soil bulk\ndensity")
diag(cor_mat) <- 0
g <- igraph::graph_from_adjacency_matrix(cor_mat, weighted=TRUE, mode="upper")
E(g)$names <- sig$edge

# filter edges to display
g <- delete_edges(g, E(g)[sig %>% filter(sig.90 == F) %>% pull(edge)]) # delete no-sig edges
# trait-trait and environment-environment
l1 <- str_detect(attr(E(g),"vnames") ,"leaf\narea|LMA|leaf N\n per area|leaf\nd13C|max\nheight|wood\ndensity", negate = T) # env-env edges
l2 <- str_detect(attr(E(g),"vnames") ,"temperature|moisture|soil P|soil bulk\ndensity", negate = T) # trait-trait edges
g1 <- delete_edges(g, E(g)[!(l1|l2)]) # delete trait-trait edges
# trait-environment
g2 <- delete_edges(g, E(g)[str_detect(attr(E(g),"vnames") ,"temperature|moisture|soil P|soil bulk\ndensity", negate = T)]) # delete trait-trait edges
g2 <- delete_edges(g2, E(g2)[str_detect(attr(E(g2),"vnames") ,"leaf\narea|LMA|leaf N\n per area|leaf\nd13C|max\nheight|wood\ndensity", negate = T)]) # delete env-env edges
# update data for plotting
sig1 <- sig %>% filter(edge %in% E(g1)$names)
sig2 <- sig %>% filter(edge %in% E(g2)$names)

# TRAIT-TRAIT AND ENVIRONMENT-ENVIRONMENT
plot(g1,
     layout = layout.circle,
     vertex.size = 20,
     vertex.label.cex=0.5,
     vertex.label.color="black",
     vertex.frame.width=2,
     vertex.frame.color="black",
     vertex.color= c("#00ba38","#00ba38","#00ba38","#00ba38","#00ba38",
                     "#00ba38","lightblue","lightblue","orange","orange"),
     edge.width = (abs(E(g1)$weight)^1)*75,
     edge.color = edge_col(sig1 %>% filter(sig == T) %>% pull(mean),
                           sig1 %>% filter(sig == T) %>% pull(alpha)),
     edge.lty = sig1 %>% filter(sig == T) %>% pull(lty)
)

# TRAIT-ENVIRONMENT
plot(g2,
     layout = layout.circle,
     vertex.size = 20,
     vertex.label.cex=0.5,
     vertex.label.color="black",
     vertex.frame.width=2,
     vertex.frame.color="black",
     vertex.color= c("#00ba38","#00ba38","#00ba38","#00ba38","#00ba38",
                     "#00ba38","lightblue","lightblue","orange","orange"),
     edge.width = (abs(E(g2)$weight)^1)*75,
     edge.color = edge_col(sig2 %>% filter(sig == T) %>% pull(mean),
                           sig2 %>% filter(sig == T) %>% pull(alpha)),
     edge.lty = sig2 %>% filter(sig == T) %>% pull(lty)
)

# PLOT PHYLOGENETIC (LOWER) AND NON-PHYLOGENETIC (UPPER) CORRELATIONS AS CORRELATION MATRIX
comb <- matrix(0,10,10)
comb[lower.tri(comb)] <- cor_list$matrices$phy_mat[lower.tri(cor_list$matrices$phy_mat)]
comb[upper.tri(comb)] <- cor_list$matrices$ind_mat[upper.tri(cor_list$matrices$ind_mat)]
dimnames(comb)[[1]] <- c("wood density","LMA","leaf N per area","leaf d13C","leaf area","max height","temperature","moisture","soil P","soil bulk density")
dimnames(comb)[[2]] <- c("wood density","LMA","leaf N per area","leaf d13C","leaf area","max height","temperature","moisture","soil P","soil bulk density")
corrplot::corrplot(comb, method = "square", cl.pos = "n", tl.col = "black", tl.cex = 1.25, addCoef.col ='black', number.cex = 1, diag = F, col = colorRampPalette(c("red","white", "blue"))(10))


#-------------------ANCESTRAL STATE RECONSTRUCTION----------------------#

# Perform ancestral state reconstructions that assess whether changes in trait values
# represent a cause or a consequence of transitions in aridity tolerance. Do so assuming
# 1) pure Brownian Motion and 2) a shift model including a directional trend toward more arid
# environments from the Miocene boundary (23 Mya), consistent with known trends in Australia's
# paleoclimatic history. These analyses require the species-level tree and fit.

# # load dummy model and parallel run
# fit_sp <- readRDS("fit_sp.rds")
# result_list_sp <- readRDS("result_list_sp.rds")

# fill dummy model with concatenated chains from parallel run
result_list <- result_list_sp
fit_sp$Sol <- result_list %>% purrr::map("fit") %>% lapply(function (x) x[c('Sol')]) %>% purrr::map(bind_rows) %>% bind_rows() %>% as.mcmc
fit_sp$VCV <- result_list %>% purrr::map("fit") %>% lapply(function (x) x[c('VCV')]) %>% purrr::map(bind_rows) %>% bind_rows() %>% as.mcmc

# prepare data for ASR
dat_sum <- dat_sp %>% group_by(taxon_sp) %>% dplyr::select(moisture_mean_taxon_sp) %>% distinct
trait <- dat_sum$moisture_mean_taxon_sp
names(trait) <- dat_sum$taxon_sp
tree <- phy_sp %>% keep.tip(dat_sum$taxon_sp)
tree$node.label <- (Ntip(tree)+1):(Ntip(tree)+Nnode(tree))

# METHOD 1 - reconstruct moisture index (MI) according to Brownian Motion (BM)
BM_states <- fastAnc(tree, trait)
root_BM <- BM_states[1]

# METHOD 2 - Reconstruct MI according to a shift model, in which: 
# 1. the root state for Eucalyptus is assumed to be humid (MI = 0.65)
#    reflecting warm, wet conditions during the Paleogene (66-23 Mya)
# 2. the evolution of MI proceeded by BM from the root (~58 Mya) of 
#    Eucalyptus until a shift point coinciding with significant climatic
#    changes in Australia during the Miocene (23-5 Mya)
# 3. the evolution of MI followed a trend toward drier (lower MI) values
#    from the Miocene boundary (23.03 Mya) to the present day
#
root_humid = 0.65 # humid classification (MI > 0.65)
shift_depth = 23.03 # shift to trend model at the Miocene boundary (23.03 Mya)
BMT_states <- adjust_ancestral_states(tree, trait, root_humid, shift_depth)

## VALIDATE SHIFT MODEL
# all bmt states are larger (wetter) than bm states due to offset 
table(BMT_states > BM_states)#; plot(BM_states, BMT_states); abline(0,1)
# model recovers supplied root state
BMT_states[1] == root_humid
# nodes deeper than shift_depth are all adjusted the same amount (i.e., the offset)
head(BMT_states,3) - head(BM_states,3)
# nodes shallower than shift_depth are adjusted according to their depth (offset + trend)
# with negligible adjustment for nodes close to the tips
tail(BMT_states,3) - tail(BM_states,3)

## CONTMAP TO VISUALISE THE EFFECT OF SHIFT MODEL VS. BM
par(mfrow=c(1,2))
# plot contmap using BM ancestral states
fit.BM<-contMap(tree,trait,plot=FALSE,method="user",anc.states=BM_states)
plot(fit.BM, fsize=c(0,1), ftype = "off", lwd = 2.5, outline = F, leg.txt="Moisture Index")
abline(v = max(node.depth.edgelength(tree))-shift_depth, col = "gray50", lty = 3, lwd = 3)
# plot contmap using BMT ancestral states
fit.BMT<-contMap(tree,trait,plot=FALSE,method="user",anc.states=BMT_states)
plot(fit.BMT, fsize=c(0,1), ftype = "off", lwd = 2.5, outline = F, leg.txt="Moisture Index")
abline(v = max(node.depth.edgelength(tree))-shift_depth, col = "gray50", lty = 3, lwd = 3)

#------------------------------------------------------------------------#

## FIT MODELS TO COMPUTE CONTRASTS BETWEEN NODE TRANSITION CATEGORIES
# Based on reconstructed moisture index (MI) values according to BM and BMT models, classify internal nodes
# into transition categories describing changes in MI between parent and child nodes. Next, fit MR-PMM
# specifying node category as a fixed effect when estimating ancestral states for functional traits.  
# Finally, compute contrasts between posterior distributions of fixed effect estimates from fitted models to  
# evaluate trait changes associated with origins, persistence and reversals in aridity tolerance.

# model and data
model = fit_sp
d.asr = dat_sp
phy.asr = phy_sp # ASR performed on species-level tree
n_samp = 100 # number of posterior draws to sample
root_humid = 0.65 # humid classification (MI > 0.65)
shift_depth = 23.03 # shift to trend model at the Miocene boundary (23.03 Mya)
S_cut = 0.5 # moisture index cut-off for defining semi-arid environments
A_cut = 0.2 # moisture index cut-off for defining arid environments
mod_traits = c("WD", "LMA", "leaf_N", "leaf_d13C", "LA", "PH","temp","moisture", "P_soil", "BD_soil")

# run parameters
nitts = 3300
burns = 300
thins = 3

# result data frames
d.trans.comb.BMT <- tibble()
trans.tab.BMT <- tibble(iter=NULL,A_A=NULL,A_S=NULL,M_M=NULL,M_S=NULL,S_A=NULL,S_M=NULL,S_S=NULL)
res.obs.MR.BMT <- c()
d.trans.comb.BM <- tibble()
trans.tab.BM <- tibble(iter=NULL,A_A=NULL,A_S=NULL,M_M=NULL,M_S=NULL,S_A=NULL,S_M=NULL,S_S=NULL)
res.obs.MR.BM <- c()

# run (takes several hours for n_samp=100. Reduce nitts/burns/thins to reduce compute time)
for (i in sample(1:nrow(model$VCV),n_samp)){
  
  # extract ancestral state estimates per posterior draw (n_samp draws)
  d.asr.pred <- pred_states_mcmcglmm(mod = model, mod_traits = mod_traits, slice = i, phy = phy.asr, data = d.asr, use.obs = T)
  
  # back transform MR-PMM node estimates to moisture index scale (MR-PMM different to BM, probably due to residual variance)
  d.asr.pred2 <-  d.asr.pred %>%
    filter(des %in% (Ntip(phy.asr)+1):(Nnode(phy.asr)+Ntip(phy.asr))) %>% 
    mutate(anc.moisture.back = exp(anc.moisture * sd(log(d.asr$moisture_mean_taxon_sp)) + mean(log(d.asr$moisture_mean_taxon_sp))))
  
  # re-order estimates from trans data frame to match phylogeny and back transform (keep on transformed scale to use distinct())
  anc.order <- d.asr.pred2 %>% dplyr::select(node=anc, anc.moisture=anc.moisture) %>% bind_rows(., d.asr.pred2 %>% dplyr::select(node=des, anc.moisture = des.moisture)) %>% distinct() %>% arrange(node) %>% 
    mutate(anc.moisture.back = exp(anc.moisture * sd(log(d.asr$moisture_mean_taxon_sp)) + mean(log(d.asr$moisture_mean_taxon_sp))))
  
  # adjust ancestral states according to shift model
  BMT_states_model <- adjust_ancestral_states(phy.asr, trait2, root_humid, shift_depth, estimates = anc.order$anc.moisture.back)
  BMT_states_model <- tibble(anc = (Ntip(phy.asr)+1):(Nnode(phy.asr)+Ntip(phy.asr)), anc.moisture = anc.order$anc.moisture, anc.moisture.back = anc.order$anc.moisture.back, anc.moisture.BMT.back = BMT_states_model)
  
  # join to trans data frame
  asr.pred <- d.asr.pred2 %>% 
    bind_rows(d.asr.pred %>% filter(!des %in% (Ntip(phy.asr)+1):(Nnode(phy.asr)+Ntip(phy.asr)))) %>% 
    left_join(., BMT_states_model, by = "anc") %>%
    mutate(obs.moisture.back = exp(obs.moisture * sd(log(d.asr$moisture_mean_taxon_sp)) + mean(log(d.asr$moisture_mean_taxon_sp))),
           anc.moisture.back = anc.moisture.back.y,
           anc.moisture = anc.moisture.y) %>% 
    dplyr::select(-anc.moisture.back.x,-anc.moisture.back.y,
                  -anc.moisture.x,-anc.moisture.y)
  
  # match in descendant states
  asr.pred$des.moisture.back <- c(BMT_states_model[match(d.asr.pred2$des, BMT_states_model$anc),]$anc.moisture.back, asr.pred$obs.moisture.back[!is.na(asr.pred$obs.moisture)])
  asr.pred$des.moisture.BMT.back <- c(BMT_states_model[match(d.asr.pred2$des, BMT_states_model$anc),]$anc.moisture.BMT.back, asr.pred$obs.moisture.back[!is.na(asr.pred$obs.moisture)])

  #------------------------------------------------------------------------------------#
  
  ## BMT MODEL
  d.trans.BMT <- asr.pred %>% mutate(anc.moisture.z = anc.moisture,
                                       des.moisture.z = des.moisture,
                                       anc.moisture = anc.moisture.BMT.back,
                                       des.moisture = des.moisture.BMT.back)
  
  # create trans data for a single posterior draw using observed data for tips and model estimates for internal nodes
  d.trans.BMT <- d.trans.BMT %>% mutate(trans = ifelse(anc.moisture > S_cut & des.moisture > S_cut,"M_M",
                                                    ifelse(anc.moisture > S_cut & des.moisture <= S_cut & des.moisture > A_cut,"M_S",
                                                    ifelse(anc.moisture > S_cut & des.moisture <= A_cut,"M_A",
                                                    ifelse(anc.moisture <= S_cut & anc.moisture > A_cut & des.moisture <= S_cut & des.moisture > A_cut,"S_S",
                                                    ifelse(anc.moisture <= S_cut & anc.moisture > A_cut & des.moisture <= A_cut,"S_A",
                                                    ifelse(anc.moisture <= A_cut & des.moisture <= A_cut,"A_A",
                                                    ifelse(anc.moisture <= A_cut & des.moisture > A_cut & des.moisture <= S_cut,"A_S",
                                                    ifelse(anc.moisture <= A_cut & des.moisture > S_cut,"A_M",
                                                    ifelse(anc.moisture <= S_cut & anc.moisture > A_cut & des.moisture > S_cut,"S_M","OTHER"))))))))))
  
  # re-classify nodes according to whether at least one descendant underwent transition
  d.trans.BMT <-  d.trans.BMT %>% group_by(anc) %>% 
    mutate(trans = case_when("M_S" %in% trans ~ "M_S", TRUE ~ trans),
           trans = case_when("M_A" %in% trans ~ "M_A", TRUE ~ trans),
           trans = case_when("S_M" %in% trans ~ "S_M", TRUE ~ trans),
           trans = case_when("S_A" %in% trans ~ "S_A", TRUE ~ trans),
           trans = case_when("A_S" %in% trans ~ "A_S", TRUE ~ trans),
           trans = case_when("A_M" %in% trans ~ "A_M", TRUE ~ trans))
  
  # slice one row per ancestral node to tabulate and plot
  trans.tab.BMT <- bind_rows(d.trans.BMT %>% slice_head(n=1) %>% pull(trans) %>% table() %>% as.matrix() %>% t %>% as_tibble() %>%  mutate(iter=i) %>% relocate(iter), trans.tab.BMT)
  
  # generate combined data set
  d.trans.comb.BMT <- d.trans.comb.BMT %>% bind_rows(d.trans.BMT %>% mutate(iter = i)) %>% mutate(animal = paste0("node",(anc-Ntip(phy.asr))))
  
  
  ## FIT MULTIVARIATE ASR MODEL
  
  # data
  d.trans.BMT <- d.trans.BMT %>% mutate(animal = paste0("node",(anc-Ntip(phy.asr))))
  d.trans.BMT <- d.trans.BMT %>% filter(animal != "node1") %>% as.data.frame # remove root node (intercept in mod) - must be DF for MCMCglmm
  
  # parameter expanded prior
  n_resp = 6
  p_exp.ASR <- list(R = list(V=diag(n_resp),nu=n_resp+1),
                G = list(G1=list(V=diag(n_resp), nu=n_resp+1, alpha.mu = rep(0,n_resp), alpha.V = diag(n_resp)*1000)))
  # fit model
  mod.obs.BMT <- suppressWarnings( # suppress warning of non-estimable fixed effects (some node classes don't occur, i.e., A_M)
    MCMCglmm(cbind(obs.LA, obs.LMA, obs.leaf_N, obs.leaf_d13C, obs.PH, obs.WD) ~ trait:trans - 1,
                      random   = ~us(trait):animal,
                      rcov     = ~us(trait):units,
                      pedigree = phy.asr,
                      family   = rep("gaussian",n_resp),
                      nodes    = "ALL", 
                      data     = d.trans.BMT,
                      prior    = p_exp.ASR,
                      nitt     = nitts, 
                      burnin   = burns,
                      thin     = thins,
                      pr=TRUE, verbose = FALSE)
    )
  
  # response variables
  resp <- c("LA", "LMA", "leaf_N", "leaf_d13C", "PH", "WD")
  
  # trans - observed data only
  for (k in 1:length(resp)){
    mod.BMT <- mod.obs.BMT
    check <- paste0("trans",c("M_M","M_S","S_S","S_A","A_A","A_S","S_M"))
    if (length(table(check %in% colnames(mod.BMT$Sol)))==2) {NULL} else {
      
      # difference between node categories
      origin <- mod.BMT$Sol[,paste0('traitobs.',resp[k],':transM_S')]-mod.BMT$Sol[,paste0('traitobs.',resp[k],':transM_M')]
      following <- mod.BMT$Sol[,paste0('traitobs.',resp[k],':transS_S')]-mod.BMT$Sol[,paste0('traitobs.',resp[k],':transM_S')]
      persistence <- mod.BMT$Sol[,paste0('traitobs.',resp[k],':transS_S')]-mod.BMT$Sol[,paste0('traitobs.',resp[k],':transM_M')]
      reversal <- mod.BMT$Sol[,paste0('traitobs.',resp[k],':transS_M')]-mod.BMT$Sol[,paste0('traitobs.',resp[k],':transS_S')]
      
      origin2 <- mod.BMT$Sol[,paste0('traitobs.',resp[k],':transS_A')]-mod.BMT$Sol[,paste0('traitobs.',resp[k],':transS_S')]
      following2 <- mod.BMT$Sol[,paste0('traitobs.',resp[k],':transA_A')]-mod.BMT$Sol[,paste0('traitobs.',resp[k],':transS_A')]
      persistence2 <- mod.BMT$Sol[,paste0('traitobs.',resp[k],':transA_A')]-mod.BMT$Sol[,paste0('traitobs.',resp[k],':transS_S')]
      reversal2 <- mod.BMT$Sol[,paste0('traitobs.',resp[k],':transA_S')]-mod.BMT$Sol[,paste0('traitobs.',resp[k],':transA_A')]
      
      res <- tibble(origin,following,persistence,reversal,origin2,following2,persistence2,reversal2, iter = i, trait = resp[k])
      res.obs.MR.BMT <- bind_rows(res,res.obs.MR.BMT)
    }
  }
  
  
  #------------------------------------------------------------------------------------#
  
  ## BM MODEL
  d.trans.BM <- asr.pred %>% mutate(anc.moisture.z = anc.moisture,
                                    des.moisture.z = des.moisture,
                                    anc.moisture = anc.moisture.back,
                                    des.moisture = des.moisture.back)
  
  # create trans data for a single posterior draw using observed data for tips and model estimates for internal nodes
  d.trans.BM <- d.trans.BM %>% mutate(trans = ifelse(anc.moisture > S_cut & des.moisture > S_cut,"M_M",
                                              ifelse(anc.moisture > S_cut & des.moisture <= S_cut & des.moisture > A_cut,"M_S",
                                              ifelse(anc.moisture > S_cut & des.moisture <= A_cut,"M_A",
                                              ifelse(anc.moisture <= S_cut & anc.moisture > A_cut & des.moisture <= S_cut & des.moisture > A_cut,"S_S",
                                              ifelse(anc.moisture <= S_cut & anc.moisture > A_cut & des.moisture <= A_cut,"S_A",
                                              ifelse(anc.moisture <= A_cut & des.moisture <= A_cut,"A_A",
                                              ifelse(anc.moisture <= A_cut & des.moisture > A_cut & des.moisture <= S_cut,"A_S",
                                              ifelse(anc.moisture <= A_cut & des.moisture > S_cut,"A_M",
                                              ifelse(anc.moisture <= S_cut & anc.moisture > A_cut & des.moisture > S_cut,"S_M","OTHER"))))))))))
  
  # re-classify nodes according to whether at least one descendant underwent transition
  d.trans.BM <-  d.trans.BM %>% group_by(anc) %>%
    mutate(trans = case_when("M_S" %in% trans ~ "M_S", TRUE ~ trans),
           trans = case_when("M_A" %in% trans ~ "M_A", TRUE ~ trans),
           trans = case_when("S_M" %in% trans ~ "S_M", TRUE ~ trans),
           trans = case_when("S_A" %in% trans ~ "S_A", TRUE ~ trans),
           trans = case_when("A_S" %in% trans ~ "A_S", TRUE ~ trans))
  
  # slice one row per ancestral node to tabulate and plot
  trans.tab.BM <- bind_rows(d.trans.BM %>% slice_head(n=1) %>% pull(trans) %>% table() %>% as.matrix() %>% t %>% as_tibble() %>%  mutate(iter=i) %>% relocate(iter), trans.tab.BM)
  
  # generate combined data set
  d.trans.comb.BM <- d.trans.comb.BM %>% bind_rows(d.trans.BM %>% mutate(iter = i)) %>% mutate(animal = paste0("node",(anc-Ntip(phy.asr))))
  
  ## FIT MULTIVARIATE ASR MODEL
  
  # data
  d.trans.BM <- d.trans.BM %>% mutate(animal = paste0("node",(anc-Ntip(phy.asr))))
  d.trans.BM <- d.trans.BM %>% filter(animal != "node1") %>% as.data.frame # remove root node (intercept in mod) - must be DF for MCMCglmm
  
  # fit model
  mod.obs.BM <- suppressWarnings( # suppress warning of non-estimable fixed effects (some node classes don't occur, i.e., A_M)
    MCMCglmm(cbind(obs.LA, obs.LMA, obs.leaf_N, obs.leaf_d13C, obs.PH, obs.WD) ~ trait:trans - 1,
                         random   = ~us(trait):animal,
                         rcov     = ~us(trait):units,
                         pedigree = phy.asr,
                         family   = rep("gaussian",n_resp),
                         nodes    = "ALL",
                         data     = d.trans.BM,
                         prior    = p_exp.ASR,
                         nitt     = nitts,
                         burnin   = burns,
                         thin     = thins,
                         pr=TRUE, verbose = FALSE)
    )
  
  # trans - observed data only
  for (k in 1:length(resp)){
    mod.BM <- mod.obs.BM
    check <- paste0("trans",c("M_M","M_S","S_S","S_A","A_A","A_S","S_M"))
    if (length(table(check %in% colnames(mod.BM$Sol)))==2) {NULL} else {
      
      # difference between node categories
      origin <- mod.BM$Sol[,paste0('traitobs.',resp[k],':transM_S')]-mod.BM$Sol[,paste0('traitobs.',resp[k],':transM_M')]
      following <- mod.BM$Sol[,paste0('traitobs.',resp[k],':transS_S')]-mod.BM$Sol[,paste0('traitobs.',resp[k],':transM_S')]
      persistence <- mod.BM$Sol[,paste0('traitobs.',resp[k],':transS_S')]-mod.BM$Sol[,paste0('traitobs.',resp[k],':transM_M')]
      reversal <- mod.BM$Sol[,paste0('traitobs.',resp[k],':transS_M')]-mod.BM$Sol[,paste0('traitobs.',resp[k],':transS_S')]
      
      origin2 <- mod.BM$Sol[,paste0('traitobs.',resp[k],':transS_A')]-mod.BM$Sol[,paste0('traitobs.',resp[k],':transS_S')]
      following2 <- mod.BM$Sol[,paste0('traitobs.',resp[k],':transA_A')]-mod.BM$Sol[,paste0('traitobs.',resp[k],':transS_A')]
      persistence2 <- mod.BM$Sol[,paste0('traitobs.',resp[k],':transA_A')]-mod.BM$Sol[,paste0('traitobs.',resp[k],':transS_S')]
      reversal2 <- mod.BM$Sol[,paste0('traitobs.',resp[k],':transA_S')]-mod.BM$Sol[,paste0('traitobs.',resp[k],':transA_A')]
      
      res2 <- tibble(origin,following,persistence,reversal,origin2,following2,persistence2,reversal2, iter = i, trait = resp[k])
      res.obs.MR.BM <- bind_rows(res2,res.obs.MR.BM)
    }
  }
  
  print(i) # n_samp iter
  
}

## save outputs
# BMT root state
saveRDS(d.trans.comb.BMT, "d.trans.comb.BMT.rds")
saveRDS(trans.tab.BMT, "trans.tab.BMT.rds")
saveRDS(res.obs.MR.BMT, "res.obs.MR.BMT.rds")
# BM root state
saveRDS(d.trans.comb.BM, "d.trans.comb.BM.rds")
saveRDS(trans.tab.BM, "trans.tab.BM.rds")
saveRDS(res.obs.MR.BM, "res.obs.MR.BM.rds")

# # load in saved runs
# d.trans.comb.BMT <- readRDS("d.trans.comb.BMT.rds")
# trans.tab.BMT <- readRDS("trans.tab.BMT.rds")
# res.obs.MR.BMT <- readRDS("res.obs.MR.BMT.rds")
# d.trans.comb.BM <- readRDS("d.trans.comb.BM.rds")
# trans.tab.BM <- readRDS("trans.tab.BM.rds")
# res.obs.MR.BM <- readRDS("res.obs.MR.BM.rds")

# CHOOSE model to assess (BM vs BMT)
d.trans.comb = d.trans.comb.BMT
trans.tab = trans.tab.BMT

# -------------------------------------------------------------#

# summarise trans category counts across posterior samples
trans.tab %>%
  mutate(M_A = ifelse(is.na(M_A), 0, M_A)) %>%  # occurs occasionally
  { if ("A_M" %in% names(.)) mutate(., A_M = ifelse(is.na(A_M), 0, A_M)) else . } %>% # occurs very occasionally
  dplyr::select(-iter) %>% 
  pivot_longer(everything(), names_to = "trans") %>% 
  group_by(trans) %>% 
  summarise_all(list(mean=mean, 
                     lower=~quantile(., probs = 0.025),
                     upper=~quantile(., probs = 0.975)))

# compute posterior mean estimates across all samples combined
d.trans.post.mean <- d.trans.comb %>% 
  group_by(anc, des, des.phy.label) %>% 
  summarise_if(is.numeric, ~ mean(.)) %>% 
  ungroup() %>%
  mutate(anc.moisture = anc.moisture.BMT.back,
         des.moisture = des.moisture.BMT.back) %>%
  mutate(trans = ifelse(anc.moisture > S_cut & des.moisture > S_cut,"M_M",
                 ifelse(anc.moisture > S_cut & des.moisture <= S_cut & des.moisture > A_cut,"M_S",
                 ifelse(anc.moisture > S_cut & des.moisture <= A_cut,"M_A",
                 ifelse(anc.moisture <= S_cut & anc.moisture > A_cut & des.moisture <= S_cut & des.moisture > A_cut,"S_S",
                 ifelse(anc.moisture <= S_cut & anc.moisture > A_cut & des.moisture <= A_cut,"S_A",
                 ifelse(anc.moisture <= A_cut & des.moisture <= A_cut,"A_A",
                 ifelse(anc.moisture <= A_cut & des.moisture > A_cut & des.moisture <= S_cut,"A_S",
                 ifelse(anc.moisture <= A_cut & des.moisture > S_cut,"A_M",
                 ifelse(anc.moisture <= S_cut & anc.moisture > A_cut & des.moisture > S_cut,"S_M","OTHER")))))))))) %>% 
  group_by(anc) %>%
  mutate(trans = case_when("M_S" %in% trans ~ "M_S", TRUE ~ trans),
         trans = case_when("M_A" %in% trans ~ "M_A", TRUE ~ trans),
         trans = case_when("S_M" %in% trans ~ "S_M", TRUE ~ trans),
         trans = case_when("S_A" %in% trans ~ "S_A", TRUE ~ trans),
         trans = case_when("A_S" %in% trans ~ "A_S", TRUE ~ trans)) %>% 
  ungroup()
# posterior mean estimates
d.trans.post.mean

# --------------------------- PLOTTING -----------------------------------#

## PLOT ESTIMATED NODE VALUES BY TRANSITION CATEGORY
order <- c("M_M","M_S","S_S","S_A","A_A","A_S","S_M")
d.plot <- d.trans.comb %>%
  filter(trans %in% order) %>%
  filter(str_detect(des.phy.label, "node", negate = T)) %>% # negate = T filter to nodes with tip daughters (data only at tips)
  group_by(trans)
# create plot
p1 <- d.plot %>% 
  ggplot(aes(x = factor(trans, levels = order), y = anc.LMA)) + 
  geom_hline(yintercept = 0, linetype = "longdash", col="grey50") +
  tidybayes::stat_slab(aes(fill=trans),
                       .width = c(.99),
                       position = position_nudge(x=0),
                       normalize = "groups", scale = 0.75, alpha = 0.85) +
  stat_pointinterval(.width = c(.5,.95),position = position_nudge(x=0),fatten_point = 1.25) +
  scale_colour_manual(values = c("red", "violet","navy","purple","darkorange1","cornflowerblue","gold")) +
  scale_fill_manual(values = c("red", "violet","navy","purple","darkorange1","cornflowerblue","gold")) +
  theme_classic() + labs(title="LMA", x="", y="") + 
  theme(plot.title = element_text(size = 14, hjust=0.5),
        axis.text.x = element_text(size = 10),  
        axis.text.y = element_text(size = 12), 
        legend.title = element_blank(), 
        legend.text = element_text(size=12)) + 
  guides(fill="none") +
  coord_flip()
p2 <- p1 + aes(y = anc.LA) + labs(title="LA", x="", y="")# + theme(axis.text.y = element_blank())
p3 <- p1 + aes(y = anc.leaf_N) + labs(title="leaf N", x="", y="")
p4 <- p1 + aes(y = anc.leaf_d13C) + labs(title="leaf d13C", x="", y="")# + theme(axis.text.y = element_blank())
p5 <- p1 + aes(y = anc.PH) + labs(title="PH", x="", y="")
p6 <- p1 + aes(y = anc.WD) + labs(title="WD", x="", y="")# + theme(axis.text.y = element_blank())
# plot
ggarrange(p1,p4,p6, nrow=3, ncol=1, common.legend = F)
ggarrange(p1,p2,p3,p4,p5,p6, nrow=3, ncol=2, common.legend = F)

## PLOT CONTRASTS
level_order <- c('origin', 'persistence',"reversal")
# data
res.list.comb <- res.obs.MR.BMT %>% 
  pivot_longer(1:8, values_to = "value", names_to = "type") %>% 
  mutate(trans = ifelse(str_detect(type, "2"), "S_A", "M_S"),
         type = str_remove(type, "2")) %>% 
  filter(type %in% level_order)
# plot
p1 <- res.list.comb %>% filter(trait == "LMA") %>% 
  ggplot(aes(y=factor(type, level = level_order), x = value)) + 
  geom_vline(xintercept = 0, linetype = "longdash", col="grey50") +
  stat_slab(aes(fill=trans),.width = c(.99),position = position_dodge(width=0.625),
            normalize = "groups", scale = 0.575, alpha = 0.8) +
  stat_pointinterval(aes(group=trans),.width = c(.5,.95),position = position_dodge(width=0.625),
                     fatten_point = 1.25) +
  scale_fill_manual(values=c("#37b578ff","#Efcc98")) +
  theme_classic() + labs(title="LMA", x="", y="") + 
  theme(plot.title = element_text(size = 14, hjust=0.5),
        axis.text.x = element_text(size = 10),  
        axis.text.y = element_text(size = 12), 
        legend.title = element_blank(), 
        legend.text = element_text(size=12)) +
  xlim(-1.3, 1.3)
p2 <- p1;p2$data <- res.list.comb %>% filter(trait == "LA");p2 <- p2 + labs(title="LA", x="", y="")# + theme(axis.text.y = element_blank())
p3 <- p1;p3$data <- res.list.comb %>% filter(trait == "leaf_N");p3 <- p3 + labs(title="leaf N", x="", y="")
p4 <- p1;p4$data <- res.list.comb %>% filter(trait == "leaf_d13C");p4 <- p4 + labs(title="leaf d13C", x="", y="")# + theme(axis.text.y = element_blank())
p5 <- p1;p5$data <- res.list.comb %>% filter(trait == "PH");p5 <- p5 + labs(title="PH", x="", y="")
p6 <- p1;p6$data <- res.list.comb %>% filter(trait == "WD");p6 <- p6 + labs(title="WD", x="", y="")# + theme(axis.text.y = element_blank())
# plot
ggarrange(p1,p4,p6, nrow=3, ncol=1,  common.legend = T, legend = "none")
ggarrange(p1,p2,p3,p4,p5,p6, nrow=3, ncol=2,  common.legend = T, legend = "none")

#-----------------------------------------------------------------#

## PLOT ASR ON TREE

# transform branch lengths for plotting
phy <- motmot::transformPhylo(phy_sp, model = "delta", delta = 10)

# use node classification based on posterior mean for plotting
post_mean_nodes <- d.trans.post.mean[order(match(d.trans.post.mean$des,phy_sp$edge[,2])),] # ensure order matches tree

# node count in each category based on posterior mean
post_mean_nodes %>% select(anc, trans) %>% distinct() %>% pull(trans) %>% table()

# identify nodes to category
M_M <- post_mean_nodes %>% select(anc, trans) %>% distinct() %>% filter(trans == "M_M") %>% pull(anc)
M_S <- post_mean_nodes %>% select(anc, trans) %>% distinct() %>% filter(trans == "M_S") %>% pull(anc)
M_A <- post_mean_nodes %>% select(anc, trans) %>% distinct() %>% filter(trans == "M_A") %>% pull(anc)
S_S <- post_mean_nodes %>% select(anc, trans) %>% distinct() %>% filter(trans == "S_S") %>% pull(anc)
S_A <- post_mean_nodes %>% select(anc, trans) %>% distinct() %>% filter(trans == "S_A") %>% pull(anc)
S_M <- post_mean_nodes %>% select(anc, trans) %>% distinct() %>% filter(trans == "S_M") %>% pull(anc)
A_A <- post_mean_nodes %>% select(anc, trans) %>% distinct() %>% filter(trans == "A_A") %>% pull(anc)
A_S <- post_mean_nodes %>% select(anc, trans) %>% distinct() %>% filter(trans == "A_S") %>% pull(anc)
A_M <- post_mean_nodes %>% select(anc, trans) %>% distinct() %>% filter(trans == "A_M") %>% pull(anc)

# plot
par(mfrow=c(1,1))
cols<-setNames(colorRampPalette(rev(viridis::plasma(6)))(510),1:510)
edge.vals<-d.trans.post.mean %>% arrange(factor(des.phy.label, levels = d.asr.pred$des.phy.label)) %>% pull(des.moisture.BMT.back) # arrange to match tree edges
edge.vals<-log(edge.vals)-mean(log(edge.vals[Ntip(phy.asr):length(edge.vals)]), na.rm=T) / sd(log(edge.vals[Ntip(phy.asr):length(edge.vals)]), na.rm=T) # back transform by tip data values
max.val<-max(edge.vals)+1e-8
min.val<-min(edge.vals)-1e-8
intervals<-seq(min.val,max.val,length.out=511)
intervals<-cbind(intervals[1:510],intervals[2:511])
edge.ind<-sapply(edge.vals, function(x,intervals) 
  intersect(which(x>intervals[,1]),which(x<=intervals[,2])),
  intervals=intervals)
painted<-paintBranches(phy,phy$edge[1,2],state=edge.ind[1])
for(i in 2:length(edge.ind)) painted<-paintBranches(painted,phy$edge[i,2],state=edge.ind[i])
plot(painted,colors=cols,lwd=1,split.vertical=TRUE,outline=F,fsize=0.025, offset = 0.75, type = "fan", ftype="off")

# tip labels
d.points <- d.asr %>% select(taxon_sp, moisture_mean_taxon_sp) %>% distinct() %>% 
  mutate(arid_cat = ifelse(moisture_mean_taxon_sp > S_cut, "M", 
                           ifelse(moisture_mean_taxon_sp > A_cut & moisture_mean_taxon_sp <= S_cut, 
                                  "S", "A")))
d.points <- d.points[order(match(d.points$taxon_sp,phy_sp$tip.label)),]
state<-setNames(as.factor(d.points %>% pull(arid_cat)), d.points %>% pull(taxon_sp))
cols2 <- c("navy", "gold","red");names(cols2) <- c("M","S","A")
tiplabels(cex = 0.75, pch = 20,
          col = cols2[as.character(state[d.points %>% pull(taxon_sp)])])

# arc labels
arc.cladelabels(text="Eudesmia",node=findMRCA(painted, dat_sp %>% filter(subgenus=="Eudesmia") %>% pull(taxon_sp)),ln.offset=1.03,lab.offset=1.075,cex=1.25,lwd=2.5, orientation = "horizontal")
arc.cladelabels(text="Symphyomyrtus",node=findMRCA(painted, dat_sp %>% filter(subgenus=="Symphyomyrtus") %>% pull(taxon_sp)),ln.offset=1.03,lab.offset=1.075,cex=1.25,lwd=2.5, orientation = "horizontal")
arc.cladelabels(text="Eucalyptus",node=findMRCA(painted, dat_sp %>% filter(subgenus=="Eucalyptus") %>% pull(taxon_sp)),ln.offset=1.03,lab.offset=1.075,cex=1.25,lwd=2.5, orientation = "horizontal")
arc.cladelabels(text="Corymbia",node=findMRCA(painted, dat_sp %>% filter(subgenus=="Corymbia") %>% pull(taxon_sp)),ln.offset=1.03,lab.offset=1.075,cex=1.25,lwd=2.5, orientation = "horizontal")
arc.cladelabels(text="Blakella",node=findMRCA(painted, dat_sp %>% filter(subgenus=="Blakella") %>% pull(taxon_sp)),ln.offset=1.03,lab.offset=1.075,cex=1.25,lwd=2.5, orientation = "horizontal")
arc.cladelabels(text="Angophora",node=findMRCA(painted, dat_sp %>% filter(subgenus=="Angophora") %>% pull(taxon_sp)),ln.offset=1.03,lab.offset=1.075,cex=1.25,lwd=2.5, orientation = "horizontal")

# node labels
nodelabels(pch = 20, col = "cornflowerblue", node=S_M, cex = 1.25)
nodelabels(pch = 20, col = "violet", node=A_S, cex = 1.25)
nodelabels(pch = 20, col = "red", node=A_A, cex = 1.25)
nodelabels(pch = 20, col = "darkorange1", node=S_A, cex = 1.25)
nodelabels(pch = 20, col = "gold", node=S_S, cex = 1.25)
nodelabels(pch = 20, col = "deeppink1", node=M_A, cex = 1.25)
nodelabels(pch = 20, col = "purple", node=M_S, cex = 1.25)
nodelabels(pch = 20, col = "navy", node=M_M, cex = 1.25)

#---------------------------------------------------------------------#
```
