## Supplementary material for "Conserved strategies underpin the replicated evolution of aridity tolerance in trees": data and code: ERE_DATA_PREP.R

```
#------------------------------------------------------------------------#
#                                                                        #
# Data Prep: Conserved strategies underpin the replicated evolution      #
# of aridity tolerance in trees                                          #
#                                                                        #
# Ben Halliwell                                                          #
# 2025                                                                   #
#                                                                        #
#------------------------------------------------------------------------#

## SETUP

# source functions
source("ERE_FUNCTIONS.R")

# # install ggtree from Bioconductor
# install.packages("BiocManager"); BiocManager::install("ggtree")

# load packages
library(tidyverse)
library(purrr)
library(ape)
library(phytools)
library(phangorn)
library(fuzzyjoin)
library(igraph)
library(ggpubr)
library(bayesplot)
library(ggtree)

# assign select function
select <- dplyr::select

# set ggplot theme
theme_set(theme_classic())

#------------------------------------------------------------------------#

## PHYLOGENY
# Read in Thornhill (2019) species-level phylogeny and harmonize with up-to-date taxonomy (Nicolle 2024).

# read in Thornhill 2019 eucalypt phylogeny
phy <- read.tree(file = 'Eucalypts_ML1_dated_r8s.tre')

# read in sheet matching Thornhill phy tip labels to species names
dat_thorn <- read.csv("Thornhill_tip_matching.csv")

# read in up-to-date Eucalyptus taxonomy (Nicolle 2024) and match in
taxonomy <- read.csv("euc_taxonomy_Nicolle_2024.csv")
dat_thorn <- dat_thorn %>% 
  mutate(series = taxonomy[match(.data$taxon_sp, taxonomy$taxon_sp),]$series,
         section = taxonomy[match(.data$taxon_sp, taxonomy$taxon_sp),]$section,
         subgenus = taxonomy[match(.data$taxon_sp, taxonomy$taxon_sp),]$subgenus)

# prune tree to taxa with known taxonomy
phy <- keep.tip(phy, dat_thorn %>% pull(thornhill_tip_label))
phy <- multi2di(phy) # make binary
phy <- force.ultrametric(phy, method = "extend", message=F)  # make ultrametric
phy$tip.label <- dat_thorn[match(phy$tip.label,dat_thorn$thornhill_tip_label),]$taxon
phy$node.label <- paste0("node",1:phy$Nnode)
phy$tip.label <- gsub('^([^_]+_[^_]+).*', "\\1", phy$tip.label) # drop subsp epithets off tip labels to help match

#------------------------------------------------------------------------#

## Infer strict consensus section-level tree from Crisp et al. (2024).

# read in partition trees presented in Crisp et al. (2024)
phy_crisp_1 <- read.tree(file="101concat-29May.nex.run1.treefile.tre")
phy_crisp_2 <- read.tree(file="101concat-29May.phy.gz.run2.treefile.tre")

# update taxonomy and filter to taxa with known series taxonomy
dat_crisp <- read.csv("Crisp_tip_matching.csv") %>% as_tibble
dat_crisp <- dat_crisp %>% 
  mutate(series = taxonomy[match(dat_crisp$taxon_sp, taxonomy$taxon_sp),]$series,
         section = taxonomy[match(dat_crisp$taxon_sp, taxonomy$taxon_sp),]$section) %>% 
  mutate(phylo = section) %>% 
  filter(str_detect(series, "hybrid", negate = T)) # remove taxonomic hybrids

# sample section-level topologies from crisp
outgroup <- c(taxonomy %>% filter(!str_detect(taxon_sp, "Eucalyptus_")) %>% pull(section) %>% unique) %>% as_tibble # define clade comprising subgenera Corymbia, Angophora, and Blakella as 'outgroup' to Eucalyptus
phy_crisp_list <- list(phy_crisp_1, phy_crisp_2)
n_tree = 1000
tree_samp_crisp <- list()
C_samp_crisp <- list()
# generate 1000 candidate topologies per partition tree
for(j in 1:length(phy_crisp_list)){
  phy_crisp <- phy_crisp_list[[j]]
  for (i in 1:n_tree) {
    samp <- dat_crisp %>% 
      group_by(phylo) %>% 
      slice_sample(n=1) # sample one species from each section and prune to generate section level topologies
    phy.sec <- keep.tip(phy_crisp, samp$taxon_crisp)
    samp <- samp %>% arrange(factor(taxon_crisp, levels = phy.sec$tip.label))
    phy.sec$tip.label <- samp$phylo %>% as.vector()
    phy.sec <- root(phy.sec, outgroup %>% filter(value %in% phy.sec$tip.label) %>% pull, resolve.root = T)
    phy.sec$node.label <- paste0("node",1:phy.sec$Nnode)
    tree_samp_crisp[[((j-1)*n_tree)+i]] <- phy.sec
    C_samp_crisp[[((j-1)*n_tree)+i]] <- vcv.phylo(phy.sec, cor = T)
  }
}

# infer section-level strict consensus tree from sampled topologies
t_con <- consensus(tree_samp_crisp, rooted = T, p = 1) # p = 1 computes strict consensus tree
t_con %>% plot(cex=0.75)

#------------------------------------------------------------------------#

## Augment Thornhill tree
# Identify species in the Thornhill tree that are incongruent with the Crisp section-level consensus tree (i.e., species with phylogenetic placement outside their respective taxonomic section).

# PLOT TREE TO IDENTIFY TAXONOMIC OUTLIERS
dat_thorn <- dat_thorn %>% mutate(section = taxonomy[match(dat_thorn$taxon_sp, taxonomy$taxon_sp),]$section,
                                  section_label = paste0(section,"_",taxon_sp))
phy_out <- phy
phy_out$tip.label <- dat_thorn[match(phy_out$tip.label, dat_thorn$taxon_sp),]$section_label

# create OTUs
grp <- list(
  Glandulosae = dat_thorn %>% filter(section == "Glandulosae") %>% pull(section_label) %>% unique,
  Adnataria = dat_thorn %>% filter(section == "Adnataria") %>% pull(section_label) %>% unique,
  Maidenaria = dat_thorn %>% filter(section == "Maidenaria") %>% pull(section_label) %>% unique,
  Frutices = dat_thorn %>% filter(section == "Frutices") %>% pull(section_label) %>% unique,
  Amentum = dat_thorn %>% filter(section == "Amentum") %>% pull(section_label) %>% unique,
  Bisectae = dat_thorn %>% filter(section == "Bisectae") %>% pull(section_label) %>% unique,
  Dumaria = dat_thorn %>% filter(section == "Dumaria") %>% pull(section_label) %>% unique,
  Eucalyptus = dat_thorn %>% filter(section == "Eucalyptus") %>% pull(section_label) %>% unique,
  Exsertaria = dat_thorn %>% filter(section == "Exsertaria") %>% pull(section_label) %>% unique,
  Reticulatae = dat_thorn %>% filter(section == "Reticulatae") %>% pull(section_label) %>% unique,
  Pumilio = dat_thorn %>% filter(section == "Pumilio") %>% pull(section_label) %>% unique,
  Latoangulatae = dat_thorn %>% filter(section == "Latoangulatae") %>% pull(section_label) %>% unique,
  Equatoria = dat_thorn %>% filter(section == "Equatoria") %>% pull(section_label) %>% unique,
  Platysperma = dat_thorn %>% filter(section == "Platysperma") %>% pull(section_label) %>% unique,
  Longistylus = dat_thorn %>% filter(section == "Longistylus") %>% pull(section_label) %>% unique,
  Aurantistamineae = dat_thorn %>% filter(section == "Aurantistamineae") %>% pull(section_label) %>% unique,
  Sejunctae = dat_thorn %>% filter(section == "Sejunctae") %>% pull(section_label) %>% unique,
  Subgen_Idiogenes = dat_thorn %>% filter(section == "Subgen_Idiogenes") %>% pull(section_label) %>% unique,
  Limbatae = dat_thorn %>% filter(section == "Limbatae") %>% pull(section_label) %>% unique,
  Incognitae = dat_thorn %>% filter(section == "Incognitae") %>% pull(section_label) %>% unique,
  Subgen_Acerosae = dat_thorn %>% filter(section == "Subgen_Acerosae") %>% pull(section_label) %>% unique,
  Bolites = dat_thorn %>% filter(section == "Bolites") %>% pull(section_label) %>% unique,
  Subgen_Cruciformes = dat_thorn %>% filter(section == "Subgen_Cruciformes") %>% pull(section_label) %>% unique,
  Domesticae = dat_thorn %>% filter(section == "Domesticae") %>% pull(section_label) %>% unique,
  Racemus = dat_thorn %>% filter(section == "Racemus") %>% pull(section_label) %>% unique,
  Subgen_Alveolata = dat_thorn %>% filter(section == "Subgen_Alveolata") %>% pull(section_label) %>% unique,
  Subgen_Cuboidea = dat_thorn %>% filter(section == "Subgen_Cuboidea") %>% pull(section_label) %>% unique,
  Notiales = dat_thorn %>% filter(section == "Notiales") %>% pull(section_label) %>% unique,
  Abbreviatae = dat_thorn %>% filter(section == "Abbreviatae") %>% pull(section_label) %>% unique,
  Naviculares = dat_thorn %>% filter(section == "Naviculares") %>% pull(section_label) %>% unique,
  Calophyllae = dat_thorn %>% filter(section == "Calophyllae") %>% pull(section_label) %>% unique,
  Maculatae = dat_thorn %>% filter(section == "Maculatae") %>% pull(section_label) %>% unique,
  Subgen_Angophora = dat_thorn %>% filter(section == "Subgen_Angophora") %>% pull(section_label) %>% unique
)

# study plot to identify taxa incongruent with the Crisp section-level consensus tree
p <- ggtree(phy_out, layout = 'rectangular') +
  geom_tiplab(size = 0.4, offset = 0)
groupOTU(p, grp, 'section') + aes(color=section) +
  theme(legend.position="right") + guides(col="none")

#------------------------------------------------------------------------#

## Exclude taxonomic outliers identified in previous step, and those flagged as uncertain in Thornhill (2019), from tree.

# exclude taxonomic outliers
drop <- read.csv("euc_taxa_to_drop.csv") # read in sheet of species to drop
keep <- phy$tip.label[!phy$tip.label %in% drop$taxon_sp]
phy <- keep.tip(phy, phy$tip.label[!phy$tip.label %in% drop$taxon_sp])

#------------------------------------------------------------------------#

## Relocate subgenus Eudesmia to be sister to subgenera Eucalyptus and Symphyomyrtus sensu Crisp et al. (2024).

# current topology
ggtree(phy, layout = 'rectangular') + geom_tiplab(size = 0.4, offset = 0)

# move clade
clade_sp <- dat_thorn %>% filter(subgenus %in% c("Eudesmia","Acerosae","Cuboidea")) %>% select(taxon_sp) %>% unique %>% filter(taxon_sp !="Eucalyptus_megasepala") %>% pull
clade <- keep.tip(phy, clade_sp)
adjust <- nodeheight(phy, 774) - nodeheight(phy, 773)
position = 2
phy_2 <- drop.tip(phy, clade_sp %>% as_tibble %>% filter(value != "Eucalyptus_tenuipes") %>% pull)
phy_2 <- bind.tree(phy_2, clade, where = 751, position = position)
phy_2$node.label <- paste0("node",1:phy_2$Nnode)
phy_2 <- drop.tip(phy_2, 681)
phy_2$edge.length[1316] <- phy_2$edge.length[1316] + adjust + position
phy_2$edge.length[phy_2$edge.length == 0] <- 0.001 # fix zero length branches
phy_2 <- force.ultrametric(phy_2, method = "extend") # make ultrametric
phy_2 <- ape::rotate(phy_2, node = 773)
phy_2 <- untangle(phy_2, method = "read.tree")

# augmented topology
ggtree(phy_2, layout = 'rectangular') + geom_tiplab(size = 0.4, offset = 0)

#------------------------------------------------------------------------#

## Generate series level topologies from augmented Thornhill tree.

# clean and group data
dat_series <- dat_thorn %>% 
  filter(taxon_sp %in% phy_2$tip.label,
         str_detect(series, "hybrid", negate = T), # exclude taxonomic hybrids
         !is.na(series)) %>% # exclude taxa with no recognised taxonomy
  mutate(phylo=series) %>% 
  group_by(series)

# run parameters
n_tree = 100
tree_samp <- list()
C_samp <- list()

# loop to sample
for (i in 1:n_tree) {
  samp <- dat_series %>% slice_sample(n=1) # sample one species per series and prune to generate series-level topologies
  phy.ser <- keep.tip(phy_2, samp$taxon_sp) # prune species-level tree to sampled taxa
  phy.ser <- multi2di(phy.ser) # make binary
  phy.ser <- force.ultrametric(phy.ser, method = "extend", message=F)  # make ultrametric
  samp <- samp %>% arrange(factor(taxon_sp, levels = phy.ser$tip.label))
  phy.ser$tip.label <- samp$phylo %>% na.omit() %>% as.vector()
  phy.ser$node.label <- paste0("node",1:phy.ser$Nnode)
  tree_samp[[i]] <- phy.ser
  C_samp[[i]] <- vcv.phylo(phy.ser, cor = T)
}
# result
tree_samp

# plot candidate topology
ggtree(tree_samp[[1]], layout = 'rectangular') + geom_tiplab(size = 2, offset = 0) + xlim(-10, 70)

#------------------------------------------------------------------------#

# LOAD IN TRAIT DATA
dat <- read.csv("euc_data.csv") %>% as_tibble

# prune trees to taxa in data
phy_sp <- keep.tip(phy_2, dat %>% filter(taxon_sp %in% phy_2$tip.label) %>% pull(taxon_sp) %>% unique) # prune to species in data
phy_sp$node.label <- paste0("node",1:phy_sp$Nnode) # update node labels
phy_ser_list <- lapply(tree_samp, keep.tip, dat$series %>% unique) # prune to series in data

## SAVE TREES
saveRDS(phy_sp, "euc_phy_sp.rds")
saveRDS(phy_ser_list, "euc_phy_ser_list.rds")

#------------------------------------------------------------------------#

## SUMMARISE EUCALYPTUS TRAIT DATA

# ASSESS DATA COVERAGE
mod_traits <- c("leaf_area", "leaf_mass_per_area", "leaf_N","leaf_delta13C", "wood_density", "plant_height_taxon_sp")

# number of observations for each trait across all taxa
dat %>% select(all_of(mod_traits), -plant_height_taxon_sp) %>% summarise_all(~ sum((!is.na(.))))

# ~40% of obs are associated with site level environmental data
dat %>% select(moisture_diff_taxon_sp) %>% filter(moisture_diff_taxon_sp!=0) %>% nrow

# number of taxa compared to recognized species 767/901 = 85%
dat %>% pull(taxon_sp) %>% unique %>% length
taxonomy %>% select(taxon_sp) %>% filter(str_detect(taxon_sp,"subsp", negate=T)) %>% filter(str_detect(taxon_sp,"_sp\\.", negate=T)) %>% unique %>% nrow()

# number and proportion of species with observations for each trait
data_coverage <- dat %>% group_by(taxon_sp) %>% summarise(across(all_of(mod_traits),~ sum(!is.na(.))))
# proportion
data_coverage %>% mutate(across(all_of(mod_traits), ~ ifelse(.>0,1,0))) %>% select(-1) %>% colSums() %>% `/`(length(unique(data_coverage$taxon_sp))) %>% round(2)
# number
data_coverage %>% mutate(across(all_of(mod_traits), ~ ifelse(.>0,1,0))) %>% select(-1) %>% colSums()

# average number of obs for each trait at the taxon_sp (species) level
data_coverage_sp <- dat %>% group_by(taxon_sp) %>% summarise(across(all_of(mod_traits),~ sum(!is.na(.))))
data_coverage_sp %>% select(-taxon_sp) %>% mutate(across(everything(), ~ na_if(., 0))) %>% colMeans(na.rm = T)
# at the series level
data_coverage_sp <- dat %>% group_by(series) %>% summarise(across(all_of(mod_traits),~ sum(!is.na(.))))
data_coverage_sp %>% select(-series) %>% mutate(across(everything(), ~ na_if(., 0))) %>% colMeans(na.rm = T)
```
