## Supplementary material for "Conserved strategies underpin the replicated evolution of aridity tolerance in trees": data and code: ERE_FUNCTIONS.R

```
#------------------------------------------------------------------------#
#                                                                        #
# Functions: Conserved strategies underpin the replicated evolution      #
# of aridity tolerance in trees                                          #
#                                                                        #
# Ben Halliwell                                                          #
# 2025                                                                   #
#                                                                        #
#------------------------------------------------------------------------#


# function to collapse rows from a common observation id into a single row
coalesce_by_column <- function(df) {
  return(dplyr::coalesce(!!! as.list(df)))
}

# function to colour edges in network plot
edge_col <- function(mean, alpha) {
  ifelse(mean > 0, rgb(0,0,1, alpha = alpha),
         rgb(1,0,0, alpha = alpha))
}

# function to create gg colour hue
gg_color_hue <- function(n) {
  hues = seq(15, 375, length = n + 1)
  hcl(h = hues, l = 65, c = 100)[1:n]
}

# function to compute variance decomposition from fitted model
var_decomp <- function(fit, data, resp_names, levels, fixed_effects = TRUE, species_cor = TRUE, res_cor = FALSE, NA_fixed = TRUE) {
  
  # index rows with complete site-level environmental data 
  rows <- data %>% rownames_to_column("row") %>% filter(!is.na(temp_site),!is.na(moisture_site),!is.na(P_site),!is.na(BD_site)) %>% pull(row) %>% as.numeric
  
  var_fe <- c() # posterior means
  var_fe_list <- list() # per draw
  var_re <- c()
  var_re_list <- list()
  var_tot <- c()
  var_tot_list <- list()
  res_var <- data.frame(trait=NULL, component=NULL, post.mean=NULL)
  for (i in 1:length(resp_names)){
    
    fe <- fit$Sol %>% as_tibble() %>% dplyr::select(contains(resp_names[i], ignore.case = F) & 
                                               !contains(paste0(resp_names[i],"_"), ignore.case = F) &
                                               # contains(paste0("trait",resp_names[i]), ignore.case = F) & # restrict to intercept to check var is zero
                                               !contains(c("node","phylo",".taxon","dataset_id")))
    
    X <- fit$X[,colnames(fe)]
    
    if (fixed_effects == FALSE) {
      X <- X[(((i-1)*(nrow(data)))+1):(i*(nrow(data)))]
    } else if (resp_names[i] %in% c("WD","LMA","leaf_N","leaf_d13C","LA")) {
      X <- X[(((i-1)*(nrow(data)))+1):(i*(nrow(data))),]
      if (NA_fixed == FALSE) {
        X <- X[rows,]
      }
    } else {
      X <- X[(((i-1)*(nrow(data)))+1):(i*(nrow(data)))]
      if (NA_fixed == FALSE) {
        X <- X[rows]
      }
    }
    
    var_fe[i] <- fe %>% colMeans %*% t(X) %>% as.vector %>% .[!is.na(.)] %>% var
    
    # calc var of fixed effects predictions per draw
    vmVarF<- numeric(length(fit$Sol[,1]))
    for(j in 1:length(fit$Sol[,1])){
      Var<-var(as.vector(as.numeric(fe[j,]) %*% t(X))) # Sol[i, 1:k]
      vmVarF[j]<-Var}
    var_fe_list[[i]] <- vmVarF
    
    # summarise var components  
    var <- fit$VCV %>% as_tibble() %>% 
      dplyr::select(contains(paste0("trait",resp_names[i]), ignore.case = F) |
               contains(paste0("trait, \""), ignore.case = F)) %>%
      dplyr::select(!contains(c(resp_names[-i]), ignore.case = F)) 
    var_re_list[[i]] <- var %>% rowSums
    var <- var %>% summarise_all(~ mean(.))
    var_re[i] <- var %>% rowSums
    
    # total var in trait
    var_tot_list[[i]] <- var_fe_list[[i]] + var_re_list[[i]]
    var_tot[i] <- var_fe[i] + var_re[i]
    
    # calculate marginal and conditional R^2
    R2m <- var_fe[i]/var_tot[i] # variance explained by fixed effects
    R2c <- (var_fe[i]+rowSums(var[,colnames(var[str_detect(colnames(var), "units", negate = T)])]))/var_tot[i]# var explained by fixed + random (excluding residual)
    
    if (resp_names[i] %in% c("WD","LMA","leaf_N","leaf_d13C","LA")) {
    R2r <- var[,colnames(var[str_detect(colnames(var), "units", negate = F)])]/var_tot[i] # var unexplained (residual)
    } else {
    R2r = 0 # remove fixed residual variance for species-level traits (fixed at 1 by default in MCMCglmm)
    }
    
    test <- mod$VCV[,str_detect(colnames(mod$VCV), "units", negate = F)]
    
    if (resp_names[i] %in% c("WD","LMA","leaf_N","leaf_d13C","LA")) {
      
      # variance partitioning
      res_v <- data.frame(trait = resp_names[i],
                          component = c("full", "fixed", "random", "residual",
                                        levels),
                          round(cbind(rbind(R2c,
                                            R2m,
                                            R2c-R2m,
                                            R2r,
                                            fit$VCV[,paste0("trait",resp_names[i],":trait",resp_names[i],".",levels[1])] %>% mean %>% {./var_tot[i]},
                                            if (species_cor == FALSE) {
                                              fit$VCV[,paste0("trait",resp_names[i],".",levels[2])] %>% mean %>% {./var_tot[i]}
                                            } else {
                                              fit$VCV[,paste0("trait",resp_names[i],":trait",resp_names[i],".",levels[2])] %>% mean %>% {./var_tot[i]}
                                            },
                                            fit$VCV[,paste0("at.level(trait, \"",resp_names[i],"\").",levels[3])] %>% mean %>% {./var_tot[i]},
                                            if (res_cor == FALSE) {
                                              fit$VCV[,paste0("trait",resp_names[i],".units")] %>% mean %>% {./var_tot[i]}
                                            } else {
                                              fit$VCV[,paste0("trait",resp_names[i],":trait",resp_names[i],".units")] %>% mean %>% {./var_tot[i]}
                                            }
                          )
                          ),3))
      names(res_v)[3] <- "post.mean";row.names(res_v) <- NULL
      
    } else {
      
      # variance partitioning
      res_v <- data.frame(trait = resp_names[i],
                          component = c("full", "fixed", "random", "residual",
                                        levels[-3]),
                          round(cbind(rbind(R2c,
                                            R2m,
                                            R2c-R2m,
                                            R2r,
                                            fit$VCV[,paste0("trait",resp_names[i],":trait",resp_names[i],".",levels[1])] %>% mean %>% {./var_tot[i]},
                                            if (species_cor == FALSE) {
                                              fit$VCV[,paste0("trait",resp_names[i],".",levels[2])] %>% mean %>% {./var_tot[i]}
                                            } else {
                                              fit$VCV[,paste0("trait",resp_names[i],":trait",resp_names[i],".",levels[2])] %>% mean %>% {./var_tot[i]}
                                            },
                                            if (res_cor == FALSE) {
                                              fit$VCV[,paste0("trait",resp_names[i],".units")] %>% mean %>% {./var_tot[i]}
                                            } else {
                                              fit$VCV[,paste0("trait",resp_names[i],":trait",resp_names[i],".units")] %>% mean %>% {./var_tot[i]}
                                            }
                          )),3))
      names(res_v)[3] <- "post.mean";row.names(res_v) <- NULL
      
      
    }
    
    res_var <- rbind(res_var, res_v)
    
  }
  
  # rename result
  names(var_fe_list) <- paste0(resp_names, "_fe_var")
  names(var_fe) <- paste0(resp_names, "_fe_var")
  names(var_re_list) <- paste0(resp_names, "_re_var")
  names(var_re) <- paste0(resp_names, "_re_var")
  names(var_tot_list) <- paste0(resp_names, "_tot_var")
  names(var_tot) <- paste0(resp_names, "_tot_var")
  
  return(list(res_var = res_var, 
              var_fe_list = var_fe_list, 
              var_fe = var_fe, 
              var_re_list = var_re_list, 
              var_re = var_re, 
              var_tot_list = var_tot_list, 
              var_tot = var_tot))
  
}

# function to calculate correlations from fitted MR-PMM
level_cor_vcv <- function(vcv, n_resp, part.1 = "phylo", part.2 =  "taxon", lambda = NULL) {
  
  # index corrmat elements for output
  len_cor <- choose(n_resp,2)
  m <- matrix(1:(n_resp^2),n_resp,n_resp)
  for (i in 1:nrow(m)) {
    for (j in i:(ncol(m))) {
      m[i,j] <- 0
    }
  }
  cols <- unique(as.vector(m));cols <- cols[cols!=0]
  
  # part 1
  draws <- vcv %>% as_tibble() %>% rename_with(tolower) %>% rename_with(~str_remove_all(., 'trait'))
  cor_draws <- draws %>% dplyr::select(all_of(contains(part.1))) %>% rename_with(~str_remove_all(., paste0('.',part.1)))
  phy_cor <- matrix(nrow = nrow(cor_draws), ncol = ncol(cor_draws))
  par_phy_cor <- matrix(nrow = nrow(cor_draws), ncol = ncol(cor_draws))
  for (i in 1:nrow(cor_draws)){
    cor <- unlist(cor_draws[i,])
    phy_res <- matrix(0, n_resp, n_resp)
    phy_res[] <- cor
    phy_res <- cov2cor(phy_res) # compute correlation
    par_phy_res <- if(is.null(lambda)) corpcor::pcor.shrink(phy_res, verbose = F) # compute partial correlation
    else(corpcor::pcor.shrink(phy_res, lambda = lambda, verbose = F))
    phy_cor[i,] <- as.vector(phy_res)
    par_phy_cor[i,] <- as.vector(par_phy_res)
  }
  phy_cor <- suppressMessages({phy_cor %>% as_tibble(.name_repair = "unique") %>% dplyr::select(all_of(cols)) %>% setNames(names(cor_draws %>% dplyr::select(all_of(cols))))})
  par_phy_cor <- suppressMessages({par_phy_cor %>% as_tibble(.name_repair = "unique") %>% dplyr::select(all_of(cols)) %>% setNames(names(cor_draws %>% dplyr::select(all_of(cols))))})
  phy_res[lower.tri(phy_res)] <- phy_cor %>% summarise(across(everything(), ~ mean(.))) %>% slice(1) %>% as.numeric(); phy_res[upper.tri(phy_res)] <- t(phy_res)[upper.tri(phy_res)]; dimnames(phy_res)[[1]] <- gsub(":","",gsub("trait","",gsub("[^:]+$","",colnames(vcv)[1:n_resp])));dimnames(phy_res)[[2]] <- gsub(":","",gsub("trait","",gsub("[^:]+$","",colnames(vcv)[1:n_resp])))
  par_phy_res[lower.tri(par_phy_res)] <- par_phy_cor %>% summarise(across(everything(), ~ mean(.))) %>% slice(1) %>% as.numeric(); par_phy_res[upper.tri(par_phy_res)] <- t(par_phy_res)[upper.tri(par_phy_res)]; dimnames(par_phy_res)[[1]] <- gsub(":","",gsub("trait","",gsub("[^:]+$","",colnames(vcv)[1:n_resp])));dimnames(par_phy_res)[[2]] <- gsub(":","",gsub("trait","",gsub("[^:]+$","",colnames(vcv)[1:n_resp])))
  
  # part 2
  draws <- vcv %>% as_tibble() %>% rename_with(tolower) %>% rename_with(~str_remove_all(., 'trait'))
  cor_draws <- draws %>% dplyr::select(all_of(contains(paste0('.',part.2)))) %>% rename_with(~str_remove_all(., paste0('.',part.2)))
  ind_cor <- matrix(nrow = nrow(cor_draws), ncol = ncol(cor_draws))
  par_ind_cor <- matrix(nrow = nrow(cor_draws), ncol = ncol(cor_draws))
  for (i in 1:nrow(cor_draws)){
    cor <- unlist(cor_draws[i,])
    ind_res <- matrix(0, n_resp, n_resp)
    ind_res[] <- cor
    ind_res <- cov2cor(ind_res)
    par_ind_res <- if(is.null(lambda)) corpcor::pcor.shrink(ind_res, verbose = F)
    else(corpcor::pcor.shrink(ind_res, lambda = lambda, verbose = F))
    ind_cor[i,] <- as.vector(ind_res)
    par_ind_cor[i,] <- as.vector(par_ind_res)
  }
  ind_cor <- suppressMessages({ind_cor %>% as_tibble(.name_repair = "unique") %>% dplyr::select(all_of(cols)) %>% setNames(names(cor_draws %>% dplyr::select(all_of(cols))))})
  par_ind_cor <- suppressMessages({par_ind_cor %>% as_tibble(.name_repair = "unique") %>% dplyr::select(all_of(cols)) %>% setNames(names(cor_draws %>% dplyr::select(all_of(cols))))})
  ind_res[lower.tri(ind_res)] <- ind_cor %>% summarise(across(everything(), ~ mean(.))) %>% slice(1) %>% as.numeric(); ind_res[upper.tri(ind_res)] <- t(ind_res)[upper.tri(ind_res)]; dimnames(ind_res)[[1]] <- gsub(":","",gsub("trait","",gsub("[^:]+$","",colnames(vcv)[1:n_resp])));dimnames(ind_res)[[2]] <- gsub(":","",gsub("trait","",gsub("[^:]+$","",colnames(vcv)[1:n_resp])))
  par_ind_res[lower.tri(par_ind_res)] <- par_ind_cor %>% summarise(across(everything(), ~ mean(.))) %>% slice(1) %>% as.numeric(); par_ind_res[upper.tri(par_ind_res)] <- t(par_ind_res)[upper.tri(par_ind_res)]; dimnames(par_ind_res)[[1]] <- gsub(":","",gsub("trait","",gsub("[^:]+$","",colnames(vcv)[1:n_resp])));dimnames(par_ind_res)[[2]] <- gsub(":","",gsub("trait","",gsub("[^:]+$","",colnames(vcv)[1:n_resp])))
  
  
  result <- list(posteriors = list(phy_cor=phy_cor,par_phy_cor=par_phy_cor,
                                   ind_cor=ind_cor,par_ind_cor=par_ind_cor), 
                 matrices = list(phy_mat = phy_res, par_phy_mat = par_phy_res,
                                 ind_mat = ind_res, par_ind_mat = par_ind_res))
  
  return(result)
  
}

# function to extract ancestral state estimates from a fitted MR-PMM
pred_states_mcmcglmm <- function(mod, mod_traits, phy, data, slice = 1, use.obs = T, post.mean = F){ 
  
  nodes <- tibble(des=c(phy$tip.label, phy$node.label), node.index=1:length(des))
  d.trans <- data.frame(anc=phy$edge[,1],des=phy$edge[,2]) %>% as_tibble() 
  d.trans <- d.trans %>% mutate(des.phy.label = nodes[match(d.trans$des, nodes$node.index),]$des)
  
  for (k in 1:length(mod_traits)) {
    
    anc.states <- mod$Sol %>% as_tibble %>% dplyr::select(contains("node")&contains(mod_traits[k], ignore.case=F)) %>% mutate(!! paste0("trait",mod_traits[k],".phylo.root") := 0) %>% relocate(paste0("trait",mod_traits[k],".phylo.root")) %>% slice(slice) %>% unlist()
    tip.states <- mod$Sol %>% as_tibble %>% dplyr::select(contains("phylo")&contains(mod_traits[k], ignore.case=F)&!contains("node")) %>% slice(slice) %>% unlist()
    node.vals<-phy$edge
    node.vals[]<-c(tip.states,anc.states)[phy$edge]
    node.vals <- node.vals + mod$Sol %>% as_tibble %>% dplyr::select(paste0("trait",mod_traits[k])) %>% slice(slice) %>% unlist() # add intercept for trait
    head(node.vals)
    
    # add predicted states for each trait as new columns in df
    d.trans <- d.trans %>% mutate(anc.trait = node.vals[,1],
                                  des.trait = node.vals[,2])
    
    
    if (post.mean == T) {
      
      anc.states <- mod$Sol %>% as_tibble %>% dplyr::select(contains("node")&contains(mod_traits[k], ignore.case=F)) %>% mutate(!! paste0("trait",mod_traits[k],".phylo.root") := 0) %>% relocate(paste0("trait",mod_traits[k],".phylo.root")) %>% colMeans() %>% unlist()
      tip.states <- mod$Sol %>% as_tibble %>% dplyr::select(contains("phylo")&contains(mod_traits[k], ignore.case=F)&!contains("node")) %>% colMeans() %>% unlist()
      node.vals<-phy$edge
      node.vals[]<-c(tip.states,anc.states)[phy$edge]
      node.vals <- node.vals + mod$Sol %>% as_tibble %>% dplyr::select(paste0("trait",mod_traits[k])) %>% colMeans() %>% unlist() # add intercept for trait
      head(node.vals)
      
      # add predicted states for each trait as new columns in df
      d.trans <- d.trans %>% mutate(anc.trait = node.vals[,1],
                                    des.trait = node.vals[,2])
      
    }
    
    
    if (use.obs == T) {
      
      # use only observed data at tips (NAs added - use estimate instead?)
      data_mean <- data %>% mutate(trait = data %>% pull(mod_traits[k])) %>% group_by(phylo) %>% summarise(trait = mean(trait, na.rm = T))
      d.trans <- d.trans %>% mutate(des.trait = ifelse(des.phy.label %in% data_mean$phylo, data_mean[match(d.trans$des.phy.label, data_mean$phylo),]$trait, des.trait))
      
    }
    
    d.trans <- d.trans %>% 
      mutate(obs.trait = ifelse(des.phy.label %in% data$phylo, des.trait, NA)) %>% 
      dplyr::rename(!! paste0("anc.",mod_traits[[k]]) := "anc.trait",
                    !! paste0("des.",mod_traits[[k]]) := "des.trait",
                    !! paste0("obs.",mod_traits[[k]]) := "obs.trait") %>% 
      relocate(des.phy.label, .after = des)
    
  }
  
  return(d.trans)
}

# function to adjust BM (mu=0) ancestral state estimates according to a shift to a BM trend model (mu!=0) 
# at a given depth in the tree. The value of mu (trend) is determined by subtracting the supplied root state 
# by the root state according to BM (offset) and dividing by the shift depth.
adjust_ancestral_states <- function(tree, trait, root.state, shift_depth, estimates = NULL) {
  
  if(is.null(estimates)){
    bm_estimates <- fastAnc(tree, trait)   # ASR according to BM
    root.BM <- bm_estimates[1]
  } else {
    bm_estimates <- estimates
    root.BM <- bm_estimates[1]
  }
  
  # calculate mu as difference between supplied root and BM root divided by the supplied shift depth 
  offset = root.state - root.BM
  # mu = offset / max(node.depth.edgelength(tree))
  mu = offset / shift_depth
  
  # calculate node depths for internal nodes
  node_depths <- abs(node.depth.edgelength(tree) - max(node.depth.edgelength(tree)))[(Ntip(tree)+1):(Ntip(tree)+Nnode(tree))]
  
  # total time (root depth)
  total_time <- max(node_depths)
  
  # adjust the BM estimates by offset and mu (for nodes with depth < shift_depth)
  adjusted_estimates <- bm_estimates
  
  for (node in 1:length(bm_estimates)) {
    if (node_depths[node] < shift_depth) {
      adjustment <- ((shift_depth - node_depths[node]) * mu)*-1 # reverse sign because adjusting back in time
      adjusted_estimates[node] <- bm_estimates[node] + offset + adjustment
    } else {
      adjusted_estimates[node] <- bm_estimates[node] + offset
    }
  }
  
  return(adjusted_estimates)
}

# function to perform leave-one-species-out cross-validation with parallel processing
loo_cv_species <- function(fit, data, trees, family, prior, n_resp, resp_names, cores = 4, n_species = 1,
                               nitt = 11000, burnin = 1000, thin = 10, calc_cor = TRUE, res_cor = FALSE, 
                               part.1 = "phylo", part.2 = "taxon_sp") {
  
  # define model
  formula  = fit$Fixed$formula
  random   = fit$Random$formula
  rcov     = fit$Residual$formula
  
  # species to leave out
  species_levels <- data %>% pull(taxon_sp) %>% unique
  species_levels <- species_levels[1:n_species]
  
  # result list
  loo_pred <- list()
  
  ## PARALLELISE
  # set up cluster to run models in parallel
  result_list <- list();gc() # list for mod results
  cl <- makeCluster(cores, outfile="")
  clusterEvalQ(cl,  library(tidyverse))
  clusterEvalQ(cl,  library(MCMCglmm))
  registerDoParallel(cl)
  { # run
    result_list <- foreach(i=1:length(species_levels)) %dopar% {
      
      # define function to calc correlations
      # function to calculate correlations from fitted MR-PMM
      level_cor_vcv <- function(vcv, n_resp, part.1 = "phylo", part.2 =  "taxon", lambda = NULL) {
        
        # index corrmat elements for output
        len_cor <- choose(n_resp,2)
        m <- matrix(1:(n_resp^2),n_resp,n_resp)
        for (i in 1:nrow(m)) {
          for (j in i:(ncol(m))) {
            m[i,j] <- 0
          }
        }
        cols <- unique(as.vector(m));cols <- cols[cols!=0]
        
        # part 1
        draws <- vcv %>% as_tibble() %>% rename_with(tolower) %>% rename_with(~str_remove_all(., 'trait'))
        cor_draws <- draws %>% dplyr::select(all_of(contains(part.1))) %>% rename_with(~str_remove_all(., paste0('.',part.1)))
        phy_cor <- matrix(nrow = nrow(cor_draws), ncol = ncol(cor_draws))
        par_phy_cor <- matrix(nrow = nrow(cor_draws), ncol = ncol(cor_draws))
        for (i in 1:nrow(cor_draws)){
          cor <- unlist(cor_draws[i,])
          phy_res <- matrix(0, n_resp, n_resp)
          phy_res[] <- cor
          phy_res <- cov2cor(phy_res) # compute correlation
          par_phy_res <- if(is.null(lambda)) corpcor::pcor.shrink(phy_res, verbose = F) # compute partial correlation
          else(corpcor::pcor.shrink(phy_res, lambda = lambda, verbose = F))
          phy_cor[i,] <- as.vector(phy_res)
          par_phy_cor[i,] <- as.vector(par_phy_res)
        }
        phy_cor <- suppressMessages({phy_cor %>% as_tibble(.name_repair = "unique") %>% dplyr::select(all_of(cols)) %>% setNames(names(cor_draws %>% dplyr::select(all_of(cols))))})
        par_phy_cor <- suppressMessages({par_phy_cor %>% as_tibble(.name_repair = "unique") %>% dplyr::select(all_of(cols)) %>% setNames(names(cor_draws %>% dplyr::select(all_of(cols))))})
        phy_res[lower.tri(phy_res)] <- phy_cor %>% summarise(across(everything(), ~ mean(.))) %>% slice(1) %>% as.numeric(); phy_res[upper.tri(phy_res)] <- t(phy_res)[upper.tri(phy_res)]; dimnames(phy_res)[[1]] <- gsub(":","",gsub("trait","",gsub("[^:]+$","",colnames(vcv)[1:n_resp])));dimnames(phy_res)[[2]] <- gsub(":","",gsub("trait","",gsub("[^:]+$","",colnames(vcv)[1:n_resp])))
        par_phy_res[lower.tri(par_phy_res)] <- par_phy_cor %>% summarise(across(everything(), ~ mean(.))) %>% slice(1) %>% as.numeric(); par_phy_res[upper.tri(par_phy_res)] <- t(par_phy_res)[upper.tri(par_phy_res)]; dimnames(par_phy_res)[[1]] <- gsub(":","",gsub("trait","",gsub("[^:]+$","",colnames(vcv)[1:n_resp])));dimnames(par_phy_res)[[2]] <- gsub(":","",gsub("trait","",gsub("[^:]+$","",colnames(vcv)[1:n_resp])))
        
        # part 2
        draws <- vcv %>% as_tibble() %>% rename_with(tolower) %>% rename_with(~str_remove_all(., 'trait'))
        cor_draws <- draws %>% dplyr::select(all_of(contains(paste0('.',part.2)))) %>% rename_with(~str_remove_all(., paste0('.',part.2)))
        ind_cor <- matrix(nrow = nrow(cor_draws), ncol = ncol(cor_draws))
        par_ind_cor <- matrix(nrow = nrow(cor_draws), ncol = ncol(cor_draws))
        for (i in 1:nrow(cor_draws)){
          cor <- unlist(cor_draws[i,])
          ind_res <- matrix(0, n_resp, n_resp)
          ind_res[] <- cor
          ind_res <- cov2cor(ind_res)
          par_ind_res <- if(is.null(lambda)) corpcor::pcor.shrink(ind_res, verbose = F)
          else(corpcor::pcor.shrink(ind_res, lambda = lambda, verbose = F))
          ind_cor[i,] <- as.vector(ind_res)
          par_ind_cor[i,] <- as.vector(par_ind_res)
        }
        ind_cor <- suppressMessages({ind_cor %>% as_tibble(.name_repair = "unique") %>% dplyr::select(all_of(cols)) %>% setNames(names(cor_draws %>% dplyr::select(all_of(cols))))})
        par_ind_cor <- suppressMessages({par_ind_cor %>% as_tibble(.name_repair = "unique") %>% dplyr::select(all_of(cols)) %>% setNames(names(cor_draws %>% dplyr::select(all_of(cols))))})
        ind_res[lower.tri(ind_res)] <- ind_cor %>% summarise(across(everything(), ~ mean(.))) %>% slice(1) %>% as.numeric(); ind_res[upper.tri(ind_res)] <- t(ind_res)[upper.tri(ind_res)]; dimnames(ind_res)[[1]] <- gsub(":","",gsub("trait","",gsub("[^:]+$","",colnames(vcv)[1:n_resp])));dimnames(ind_res)[[2]] <- gsub(":","",gsub("trait","",gsub("[^:]+$","",colnames(vcv)[1:n_resp])))
        par_ind_res[lower.tri(par_ind_res)] <- par_ind_cor %>% summarise(across(everything(), ~ mean(.))) %>% slice(1) %>% as.numeric(); par_ind_res[upper.tri(par_ind_res)] <- t(par_ind_res)[upper.tri(par_ind_res)]; dimnames(par_ind_res)[[1]] <- gsub(":","",gsub("trait","",gsub("[^:]+$","",colnames(vcv)[1:n_resp])));dimnames(par_ind_res)[[2]] <- gsub(":","",gsub("trait","",gsub("[^:]+$","",colnames(vcv)[1:n_resp])))
        
        
        result <- list(posteriors = list(phy_cor=phy_cor,par_phy_cor=par_phy_cor,
                                         ind_cor=ind_cor,par_ind_cor=par_ind_cor), 
                       matrices = list(phy_mat = phy_res, par_phy_mat = par_phy_res,
                                       ind_mat = ind_res, par_ind_mat = par_ind_res))
        
        return(result)
        
      }
      
      # create training data
      print(i)
      species <- species_levels[i] # left out species
      train_data <- data
      train_data[train_data$taxon_sp == species, resp_names] <- NA # delete response data for left out species but retain grouping variables to predict
      
      # fit model
      C <- MCMCglmm::inverseA(trees[[1]])$Ainv
      fit <- MCMCglmm(formula, 
                      random = random, 
                      rcov = rcov,
                      ginv = list(phylo = C),
                      family = family,
                      data = train_data, 
                      prior = prior,
                      nitt = nitt, 
                      burnin = burnin, 
                      thin = thin,
                      verbose = FALSE,
                      pr=TRUE)
      
      # rows left out
      loo_rows <- as.integer(rownames(data[data$taxon_sp == species,]))
      
      # calculate correlations
      if (calc_cor == TRUE) {
        
        if (is.null(part.2)) {
          
          cor_list <- level_cor_vcv(fit$VCV, n_resp, part.1 = part.1, part.2 = part.1, lambda = NULL)
          phy_cor_post <- cor_list$posteriors[["par_phy_cor"]]
          ind_cor_post <- NULL
          
        } else {
          
          cor_list <- level_cor_vcv(fit$VCV, n_resp, part.1 = part.1, part.2 = part.2, lambda = NULL)
          phy_cor_post <- cor_list$posteriors[["par_phy_cor"]]
          ind_cor_post <- cor_list$posteriors[["par_ind_cor"]]
          
        }
        
      } else {
        
        phy_cor_post <- NULL
        ind_cor_post <- NULL
        
      }
      
      # re-fit to predict left-out species
      test_data <- data
      pred <- predict(fit, newdata = test_data, marginal = NULL, type = "response")
      pred <- matrix(pred, ncol = n_resp)
      pred <- matrix(nrow=length(loo_rows), data=pred[loo_rows,])
      dimnames(pred) <- list(NULL, resp_names)
      
      # observed data
      observed = test_data[as.integer(rownames(data[data$taxon_sp == species,])), resp_names]
      
      # predict with confidence interval
      conf.interval <- predict(fit, newdata = test_data, marginal = NULL, type = "response", interval = "confidence") %>% as_tibble
      conf.interval <- conf.interval %>% 
        mutate(trait = rep(resp_names, each = nrow(train_data)),
               Species = rep(train_data$taxon_sp, n_resp)) %>% 
        filter(Species == species) %>% 
        group_by(trait) %>% 
        summarise(mean = mean(fit),
                  lwr = min(lwr),
                  upr = max(upr)) %>% 
        mutate(species = species)
      
      # predict with prediction interval
      pred.interval <- predict(fit, newdata = test_data, marginal = NULL, type = "response", interval = "prediction") %>% as_tibble
      pred.interval <- pred.interval %>% 
        mutate(trait = rep(resp_names, each = nrow(train_data)),
               Species = rep(train_data$taxon_sp, n_resp)) %>% 
        filter(Species == species) %>% 
        group_by(trait) %>% 
        summarise(mean = mean(fit),
                  lwr = min(lwr),
                  upr = max(upr)) %>% 
        mutate(species = species)
      
      # residual
      if (res_cor == TRUE) {
        # extract the posterior samples of the residual covariance matrix
        Sigma_posterior <- fit$VCV[, paste0("trait", rep(resp_names, each = n_resp), ":", "trait", rep(resp_names, times = n_resp), ".units")]
        Sigma <- matrix(Sigma_posterior %>% colMeans(), n_resp, n_resp)
      } else {
        # define VCV as I%*%Ve
        Sigma = diag(n_resp) * fit$VCV[,paste0("trait",resp_names,".units")] %>% colMeans()
      }
      
      # calculate predictive log likelihoods for multivariate trait draw for left-out species (integrate out missing values)
      pld_values <- c()
      for (i in 1:nrow(observed)){
        idx <- !is.na(observed[i,])
        if (sum(idx)>1){
          pld_values[i] <- mvtnorm::dmvnorm(pred[i,idx]- observed[i,idx], mean = rep(0, sum(idx)), sigma = Sigma[idx,idx], log = T) 
        } else {
          pld_values[i] <- dnorm(pred[i,idx]- observed[i,idx], mean = rep(0, sum(idx)), sd = Sigma[idx,idx], log = T) 
        }
      }
      
      # store result
      list(observed = observed,
           predicted = pred,
           predictive_interval = pred.interval,
           confidence_interval = conf.interval,
           sigma = Sigma,
           pld = pld_values,
           par_phy_cor = phy_cor_post,
           par_ind_cor = ind_cor_post)
    }
  }
  
  loo_pred <- result_list
  names(loo_pred) <- species_levels
  
  return(loo_pred)
}

# function to perform posterior predictive checks
ppc <- function(fit, data, n_resp, resp_names, n_samples = 100, res_cor = FALSE) {
  
  # remove fixed residual variances for species-level traits (PH, and enviro vars)
  fit$VCV[,str_detect(colnames(fit$VCV),"units")][,6:10] <- 0
  
  # number of observations
  n <- nrow(data)
  
  # Create result list
  y_pred <- list()
  
  for (i in 1:n_samples) {
    # Compute the predicted mean for each response variable
    W.1 <- cbind(fit$X, fit$Z)
    y_hat <- W.1 %*% fit$Sol[i, ] # multiply design matrix by posterior coefficient estimates
    
    if (res_cor == TRUE) {
      # extract the posterior samples of the residual covariance matrix
      Sigma_posterior <- fit$VCV[i, paste0("trait", rep(resp_names, each = n_resp), ":", "trait", rep(resp_names, times = n_resp), ".units")]
      Sigma <- matrix(Sigma_posterior, n_resp, n_resp)
    } else {
      # define VCV as I%*%Ve
      Sigma = diag(n_resp) * rep(fit$VCV[i,paste0("trait",resp_names,".units")])
    }
    
    # Create block diagonal covariance matrix for the entire dataset
    y_hat_mat <- matrix(y_hat, ncol = n_resp) # separate conditional means for each trait into separate columns
    y_sig <- MASS::mvrnorm(n = n, # simulate residual
                           mu = rep(0,n_resp),
                           Sigma = Sigma)
    y_pred[[i]] <- y_sig + y_hat_mat # combine for simulation
  }
  
  # Convert predictions to a dataframe for easy plotting
  y_pred_df <- data.frame(sim = rep(1:n_samples, each = nrow(data)),
                          y_pred = do.call(rbind, y_pred))
  response_vars <- colnames(data[, resp_names])
  colnames(y_pred_df)[2:(n_resp + 1)] <- response_vars
  
  return(list(y_pred_df = y_pred_df, response_vars = response_vars))
}

# function to create and arrange posterior predictive check plots in a grid
create_ppc_plots <- function(ppc_result, data) {
  pp_plots <- list()
  
  for (var in ppc_result$response_vars) {
    ppc_plot <- ggplot(ppc_result$y_pred_df, aes(x = !!sym(var))) +
      geom_density(alpha = 0.5, colour = 'royalblue', aes(group = sim)) +
      geom_density(data = data.frame(x = data[[var]]), aes(x = x), color = 'black', linewidth = 1)
    
    pp_plots[[var]] <- ppc_plot
  }
  return(pp_plots)
}
```
